# Reconciling Kinetic and Equilibrium Models of Bacterial Transcription

**DOI:** 10.1101/2020.06.13.150292

**Authors:** Muir Morrison, Manuel Razo-Mejia, Rob Phillips

## Abstract

The study of transcription remains one of the centerpieces of modern biology with implications in settings from development to metabolism to evolution to disease. Precision measurements using a host of different techniques including fluorescence and sequencing readouts have raised the bar for what it means to quantitatively understand transcriptional regulation. In particular our understanding of the simplest genetic circuit is sufficiently refined both experimentally and theoretically that it has become possible to carefully discriminate between different conceptual pictures of how this regulatory system works. This regulatory motif, originally posited by Jacob and Monod in the 1960s, consists of a single transcriptional repressor binding to a promoter site and inhibiting transcription. In this paper, we show how seven distinct models of this so-called simple-repression motif, based both on equilibrium and kinetic thinking, can be used to derive the predicted levels of gene expression and shed light on the often surprising past success of the equilibrium models. These different models are then invoked to confront a variety of different data on mean, variance and full gene expression distributions, illustrating the extent to which such models can and cannot be distinguished, and suggesting a two-state model with a distribution of burst sizes as the most potent of the seven for describing the simple-repression motif.

## 1 Introduction

Gene expression presides over much of the most important dynamism of living organisms. The level of expression of batteries of different genes is altered as a result of spatiotemporal cues that integrate chemical, mechanical and other types of signals. The original repressor-operator model conceived by Jacob and Monod in the context of bacterial metabolism has now been transformed into the much broader subject of gene regulatory networks in living organisms of all kinds [1]–[3]. One of the remaining outstanding challenges to have emerged in the genomic era is our continued inability to predict the regulatory consequences of different regulatory architectures, i.e. the arrangement and affinity of binding sites for transcription factors and RNA polymerases on the DNA. This challenge stems first and foremost from our ignorance about what those architectures even are, with more than 60% of the genes even in an ostensibly well understood organism such as *E. coli* having no regulatory insights at all [4]–[7]. But even once we have established the identity of key transcription factors and their binding sites of a given promoter architecture, there remains the predictive challenge of understanding its input-output properties, an objective that can be met by a myriad of approaches using the tools of statistical physics [8]–[25]. One route to such predictive understanding is to focus on the simplest regulatory architecture and to push the theory-experiment dialogue as far and as hard as it can be pushed [26], [27]. If we demonstrate that we can pass that test by successfully predicting both the means and variance in gene expression at the mRNA level, then that provides a more solid foundation upon which to launch into more complex problems - for instance, some of the previously unknown architectures uncovered in [5] and [28].

To that end, in this paper we examine a wide variety of distinct models for the simple repression regulatory architecture. This genetic architecture consists of a DNA promoter regulated by a transcriptional repressor that binds to a single binding site as developed in pioneering early work on the quantitative dissection of transcription [29], [30]. All of the proposed models coarse-grain away some of the important microscopic features of this architecture that have been elucidated by generations of geneticists, molecular biologists and biochemists. One goal in exploring such coarse-grainings is to build towards the future models of regulatory response that will be able to serve the powerful predictive role needed to take synthetic biology from a brilliant exercise in enlightened empiricism to a rational design framework as in any other branch of engineering. More precisely, we want phenomenology in the sense of coarse-graining away atomistic detail, but still retaining biophysical meaning. For example, we are not satisfied with the strictly phenomenological approach offered by the commonly used Hill functions. As argued in [31], Hill functions are ubiquitous precisely because they coarse-grain away all biophysical details into inscrutable parameters. Studies like [32] have demonstrated that Hill functions are clearly insufficient since each new situation requires a completely new set of parameters. Such work requires a quantitative theory of how biophysical changes at the molecular level propagate to input-output functions at the genetic circuit level. In particular a key question is: at this level of coarse-graining, what microscopic details do we need to explicitly model, and how do we figure that out? For example, do we need to worry about all or even any of the steps that individual RNA polymerases go through each time they make a transcript? Turning the question around, can we see any imprint of those processes in the available data? If the answer is no, then those processes are irrelevant for our purposes. Forward modeling and inverse (statistical inferential) modeling are necessary to tackle such questions.

Figure 1(A) shows the qualitative picture of simple repression that is implicit in the repressor-operator model. An operator, the binding site on the DNA for a repressor protein, may be found occupied by a repressor, in which case transcription is blocked from occurring. Alternatively, that binding site may be found unoccupied, in which case RNA polymerase (RNAP) may bind and transcription can proceed. The key assumption we make in this simplest incarnation of the repressor-operator model is that binding of repressor and RNAP in the promoter region of interest is exclusive, meaning that one or the other may bind, but never may both be simultaneously bound. It is often imagined that when the repressor is bound to its operator, RNAP is sterically blocked from binding to its promoter sequence. Current evidence suggests this is sometimes, but not always the case, and it remains an interesting open question precisely how a repressor bound far upstream is able to repress transcription [4]. Suggestions include “action-at-a-distance” mediated by kinks in the DNA, formed when the repressor is bound, that prevent RNAP binding. Nevertheless, our modeling in this work is sufficiently coarse-grained that we simply assume exclusive binding and leave explicit accounting of these details out of the problem.

**Figure 1.**
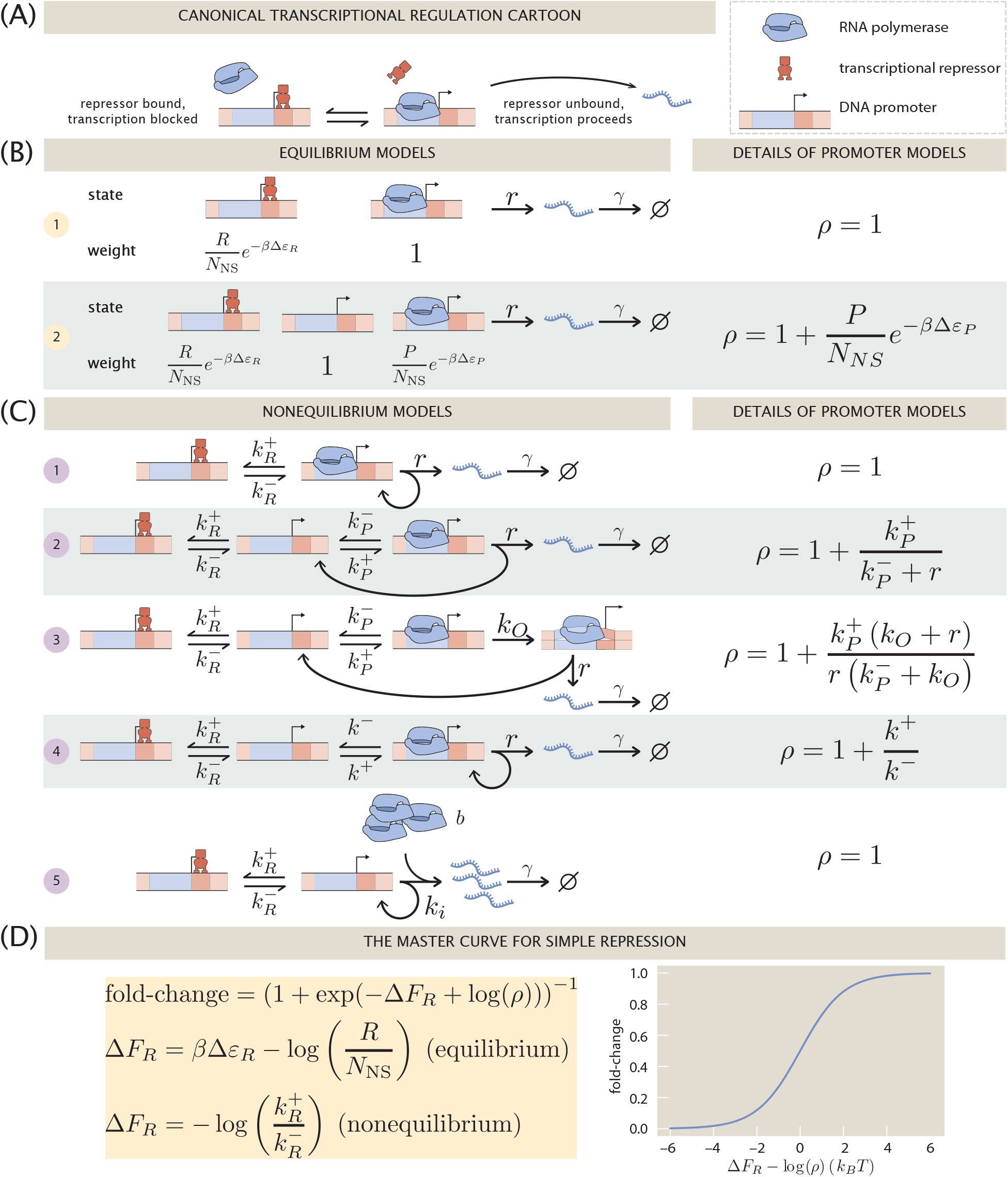
An overview of the simple repression motif at the level of means. (A) Schematic of the qualitative biological picture of the simple repression genetic architecture. (B) and (C) A variety of possible mathematicized cartoons of simple repression, along with the effective parameter *ρ* which subsumes all regulatory details of the architecture that do not directly involve the repressor. (B) Simple repression models from an equilibrium perspective. (C) Equivalent models cast in chemical kinetics language. (D) The “master curve” to which all cartoons in (B) and (C) collapse.

The logic of the remainder of the paper is as follows. In section 2, we show how both thermodynamic models and kinetic models based upon the chemical master equation all culminate in the same underlying functional form for the fold-change in the average level of gene expression as shown in Figure 1(D). Section 3 goes beyond an analysis of the mean gene expression by asking how the same models presented in Figure 1(C) can be used to explore noise in gene expression. To make contact with experiment, all of these models must make a commitment to some numerical values for the key parameters found in each such model. Therefore in Section 4 we explore the use of Bayesian inference to establish these parameters and to rigorously answer the question of how to discriminate between the different models.

## 2 Mean Gene Expression

As noted in the previous section, there are two broad classes of models in play for computing the input-output functions of regulatory architectures as shown in Figure 1. In both classes of model, the promoter is imagined to exist in a discrete set of states of occupancy, with each such state of occupancy accorded its own rate of transcription – including no transcription for many of these states. The models are probabilistic with each state assigned some probability and the overall rate of transcription given by

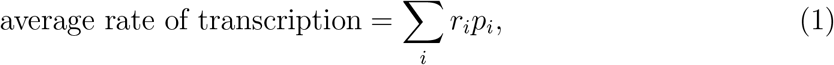

where *i* labels the distinct states, *p_i_* is the probability of the *i*^th^ state, and *r_i_* is the rate of transcription of that state. Ultimately, the different models differ along several key axes: what states to consider and how to compute the probabilities of those states.

The first class of models that are the focus of the present section on predicting the mean level of gene expression, sometimes known as thermodynamic models, invoke the tools of equilibrium statistical mechanics to compute the probabilities [8]–[17]. As seen in Figure 1(B), even within the class of thermodynamic models, we can make different commitments about the underlying microscopic states of the promoter. Indeed, the list of options considered here does not at all exhaust the suite of different microscopic states we can assign to the promoter.

The second class of models that allow us to access the mean gene expression use chemical master equations to compute the probabilities of the different microscopic states [18]–[25].

As seen in Figure 1(C), we consider a host of different nonequilibrium models, each of which will have its own result for both the mean (this section) and noise (next section) in gene expression.

### 2.1 Fold-changes are indistinguishable across models

As a first stop on our search for the “right” model of simple repression, let us consider what we can learn from theory and experimental measurements on the average level of gene expression in a population of cells. One experimental strategy that has been particularly useful (if incomplete since it misses out on gene expression dynamics) is to measure the fold-change in mean expression. The fold-change is defined as

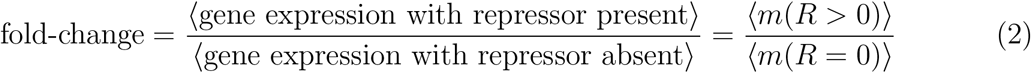

where angle brackets 〈·〉 denote the average over a population of cells and mean mRNA 〈*m*〉 is viewed as a function of repressor copy number *R*. What this means is that the fold-change in gene expression is a relative measurement of the effect of the transcriptional repressor (*R* > 0) on the gene expression level compared to an unregulated promoter (*R* = 0). The second equality in Eq. 2 follows from assuming that the translation efficiency, i.e., the number of proteins translated per mRNA, is the same in both conditions. In other words, we assume that mean protein level is proportional to mean mRNA level, and that the proportionality constant is the same in both conditions and therefore cancels out in the ratio. This is reasonable since the cells in the two conditions are identical except for the presence of the transcription factor, and the model assumes that the transcription factor has no direct effect on translation.

Fold-change has proven a very convenient observable in past work [32]–[35]. Part of its utility in dissecting transcriptional regulation is its ratiometric nature, which removes many secondary effects that are present when making an absolute gene expression measurement. Also, by measuring otherwise identical cells with and without a transcription factor present, any biological noise common to both conditions can be made to cancel away.

Figure 1 depicts a smorgasbord of mathematicized cartoons for simple repression using both thermodynamic and kinetic models that have appeared in previous literature. For each cartoon, we calculate the fold-change in mean gene expression as predicted by that model, deferring some algebraic details to Appendix S1. What we will find is that all cartoons collapse to a single master curve, shown in Figure 1(D), which contains just two parameters. We label the parameters Δ*F_R_*, an effective free energy parametrizing the repressor-DNA interaction, and *ρ*, which subsumes all details of transcription in the absence of repressors. We will offer some intuition for why this master curve exists and discuss why at the level of the mean expression, we are unable to discriminate “right” from “wrong” cartoons given only measurements of fold-changes in expression.

#### 2.1.1 The Two-State Equilibrium Model

In this simplest model, depicted as (1) in Figure 1(B), the promoter is idealized as existing in one of two states, either repressor bound or repressor unbound. The rate of transcription is assumed to be proportional to the fraction of time spent in the repressor unbound state. From the relative statistical weights listed in Figure 1, the probability *p_U_* of being in the unbound state is

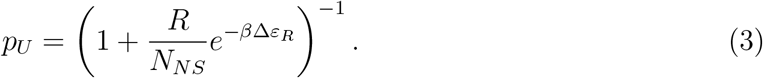

The mean rate of transcription is then given by *rp_U_*, as assumed by Eq. 1. The mean number of mRNA is set by the balance of average mRNA transcription and degradation rates, so it follows that the mean mRNA level is given by

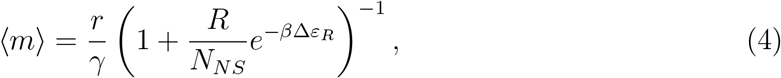

where *r* is the transcription rate from the repressor unbound state, *γ* is the mRNA degradation rate, *R* is repressor copy number, *N_NS_* is the number of nonspecific binding sites in the genome where repressors spend most of their time when not bound to the operator, *β* ≡ 1/*k_B_T*, and Δ*ε_R_* is the binding energy of a repressor to its operator site. The derivation of this result is deferred to Appendix S1.

The fold-change is found as the ratio of mean mRNA with and without repressor as introduced in Eq. 2. Invoking that definition results in

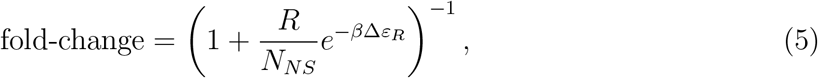

which matches the form of the master curve in Figure 1(D) with *ρ* = 1 and Δ*F_R_* = *β*Δ*ε_r_* − log(*R/N_NS_*).

In fact it was noted in [35] that this two-state model can be viewed as the coarse-graining of any equilibrium promoter model in which no transcriptionally active states have transcription factor bound, or put differently, when there is no overlap between transcription factor bound states and transcriptionally active states. We will see this explicitly in the 3-state equilibrium model below, but perhaps surprising is that an analogous result carries over even to the nonequilibrium models we consider later.

#### 2.1.2 The Three-State Equilibrium Model

Compared to the previous model, here we fine-grain the repressor unbound state into two separate states: empty, and RNAP bound as shown in (2) in Figure 1(B). This picture was used in [33] as we use it here, and in [32] and [35] it was generalized to incorporate small-molecule inducers that bind the repressor. The effect of this generalization is, roughly speaking, simply to rescale *R* from the total number of repressors to a smaller effective number of available repressors which are unbound by inducers. We point out that the same generalization can be incorporated quite easily into any of our models in Figure 1 by simply rescaling the repressor copy number *R* in the equilibrium models, or equivalently *k*^+^ in the nonequilibrium models.

The mean mRNA copy number, as derived in Appendix S1 from a similar enumeration of states and weights as the previous model, is

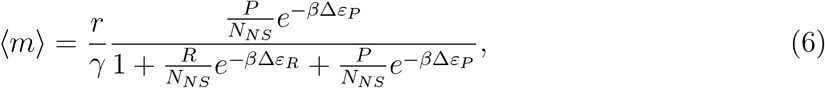

where the new variables are Δ*ε_P_*, the difference in RNAP binding energy to its specific site (the promoter) relative to an average nonspecific background site, and the RNAP copy number, *P*. The fold-change again follows immediately as

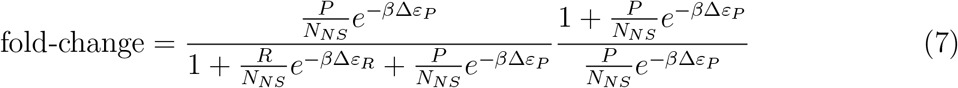

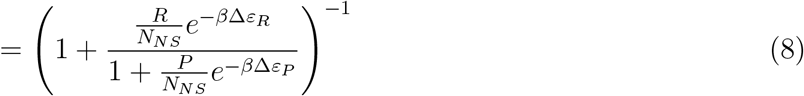

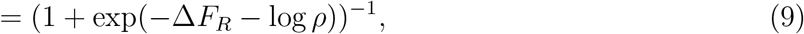

with Δ*F_R_* = *β*Δ*ε_R_* − log(*R/N_NS_*) and 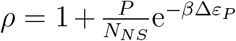 as shown in Figure 1(B). Thus far, we see that the two thermodynamic models, despite making different coarse-graining commitments, result in the same functional form for the fold-change in mean gene expression. We now explore how kinetic models fare when faced with computing the same observable.

#### 2.1.3 The Poisson Promoter Nonequilibrium Model

For our first kinetic model, we imitate the states considered in the Two-State Equilibrium Model and consider the simplest possible picture with only two states, repressor bound and unbound. This is exactly the model used for the main results of [36]. In this picture, repressor association and dissociation rates from its operator site, 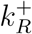 and 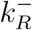, respectively, govern transitions between the two states. When the system is in the unbound state, transcription initiates at rate *r*, which represents a coarse-graining of all the downstream processes into a single effective rate. mRNA is degraded at rate *γ* as already exploited in the previous models.

Let *p_R_*(*m, t*) denote the joint probability of finding the system in the repressor bound state *R* with *m* mRNA molecules present at time *t*. Similarly define *p_U_* (*m, t*) for the repressor unbound state *U*. This model is governed by coupled master equations giving the time evolution of *p_R_*(*m, t*) and *p_U_* (*m, t*) [22], [24], [27] which we can write as

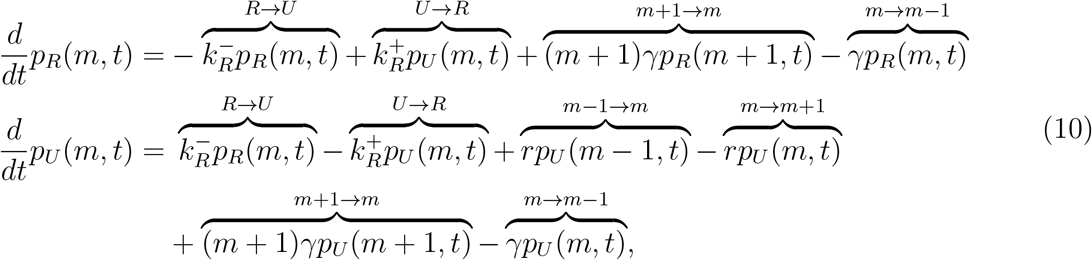

where each term on the right corresponds to a transition between two states of the promoter as indicated by the overbrace label. In each equation, the first two terms describe transitions between promoter states due to repressors unbinding and binding, respectively. The final two terms describe degradation of mRNA, decreasing the copy number by one, and the terms with coefficient *r* describe transcription initiation increasing the mRNA copy number by one. We direct the reader to Appendix S1.1 for a careful treatment showing how the form of this master equation follows from the corresponding cartoon in Figure 1.

We can greatly simplify the notation, which will be especially useful for the more complicated models yet to come, by re-expressing the master equation in vector form [37]. The promoter states are collected into a vector and the rate constants are collected into matrices as

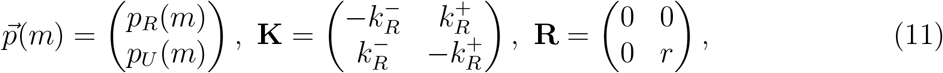

so that the master equation may be condensed as

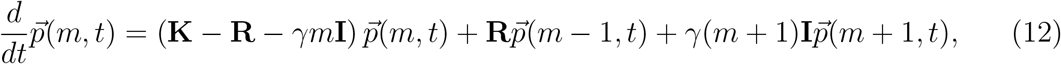

where **I** is the identity matrix. Taking steady state by setting time derivatives to zero, the mean mRNA can be found to be

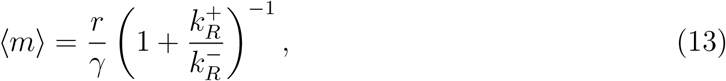

with the algebra details again deferred to Appendix S1. Recall 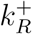 is proportional to the repressor copy number, so in computing fold-change, absence of repressor corresponds to 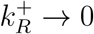. Therefore fold-change in this model is simply

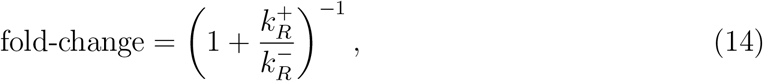

again matching the master curve of Figure 1(D) with *ρ* = 1.

#### 2.1.4 Nonequilibrium Model Two - RNAP Bound and Unbound States

Our second kinetic model depicted in Figure 1(C) mirrors the second equilibrium model of Figure 1(B) by fine-graining the repressor unbound state of nonequilibrium model 1, resolving it into an empty promoter state and an RNAP-bound state. Note in this picture, in contrast with model 4 below, transcription initiation is accompanied by a promoter state change, in keeping with the interpretation as RNAP-bound and empty states: if an RNAP successfully escapes the promoter and proceeds to elongation of a transcript, clearly it is no longer bound at the promoter. Therefore another RNAP must bind before another transcript can be initiated.

The master equation governing this model is analogous to Eqs. 11-12 for model 1 above. The main subtlety arises since transcription initiation accompanies a promoter state change. This can be understood by analogy to **K**. The off-diagonal and diagonal elements of **K** correspond to transitions arriving at or departing from, respectively, the promoter state of interest. If transcription initiation is accompanied by promoter state changes, we must have separate matrices for arriving and departing transcription events since the arriving and departing transitions have different initial copy numbers of mRNA, unlike for **K** where they are the same (see Appendix S1). The master equation for this model is

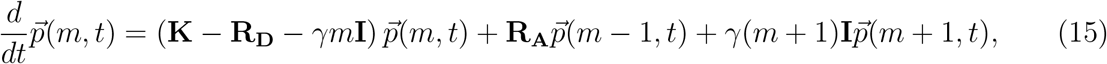

with the state vector and promoter transition matrix defined as

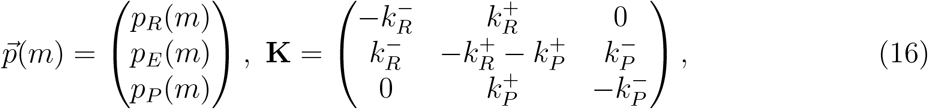

and the initiation matrices given by

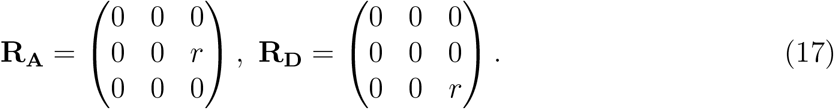

The elements of 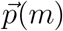 encode the probabilities of having *m* mRNA present along with the promoter having repressor bound (*R*), being empty (*E*), or having RNAP bound (*P*), respectively. **R_A_** describes probability flux *arriving* at the state 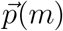 from a state with one fewer mRNA, namely 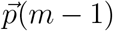, and **R_D_** describes probability flux *departing* from the state 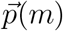 for a state with one more mRNA, namely 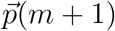. **K** is closely analogous to model 1.

Mean mRNA at steady state is found analogously to model 1, with the result

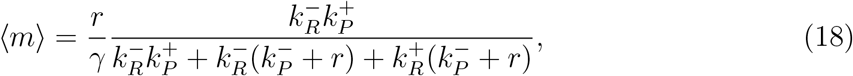

and with details again deferred to Appendix S1. Fold-change is again found from the ratio prescribed by Eq. 2, from which we have

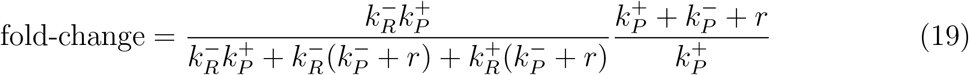

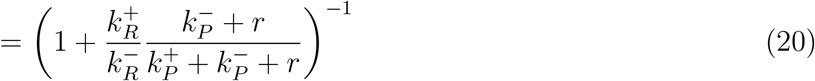

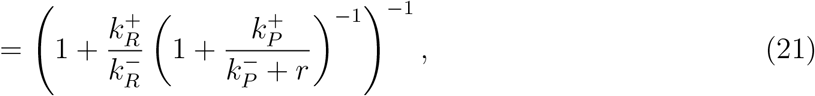

which follows the master curve of Figure 1(D) with 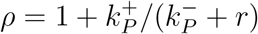 as claimed.

#### 2.1.5 Nonequilibrium Model Three - Multistep Transcription Initiation and Escape

One might reasonably complain that the first two “nonequilibrium” models we have considered are straw men. Their steady states necessarily satisfy detailed balance which is equivalent to thermodynamic equilibrium. Why is this the case? At steady state there is by definition no net probability flux in or out of each promoter state, but since the promoter states form a linear chain, there is only one way in or out of the repressor bound and RNAP bound states, implying each edge must actually have a net zero probability flux, which is the definition of detailed balance (usually phrased as equality of forward and reverse transition fluxes).

Now we consider model 3 in Figure 1(C) which allows the possibility of true nonequilibrium steady-state fluxes through the promoter states. We point out that this model was considered previously in [38] where a comparison was made with model 1 as used in [36]. The authors of [38] argued that the additional complexity is essential to properly account for the noise in the mRNA distribution. We will weigh in on both models later when we consider observables beyond fold-change.

The master equation governing this model is identical in form to model 2 above, namely

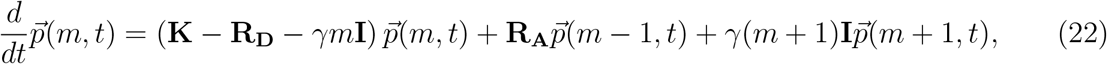

but with a higher-dimensional state space and different matrices. The state vector and promoter transition matrix are now

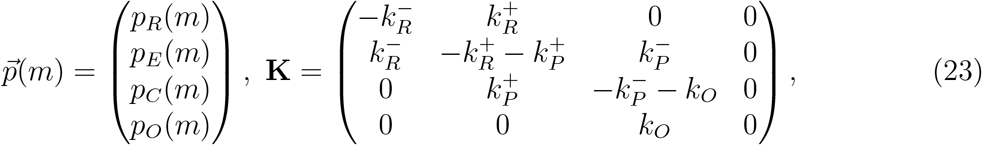

with the four promoter states, in order, being repressor bound (*R*), empty (*E*), RNAP closed complex (*C*), and RNAP open complex (*O*). Besides increasing dimension by one, the only new feature in **K** is the rate *k_O_*, representing the rate of open complex formation from the closed complex, which we assume for simplicity to be irreversible in keeping with some [38] but not all [39] past literature. The initiation matrices are given by

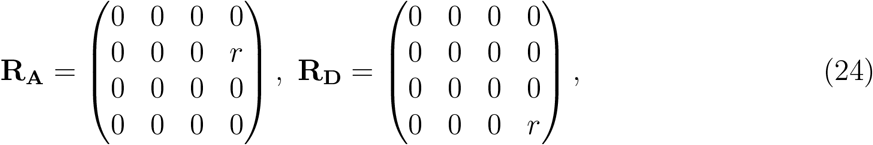

again closely analogous to nonequilibrium model 2.

The expression for mean mRNA is substantially more complicated now, as worked out in Appendix S1 where we find

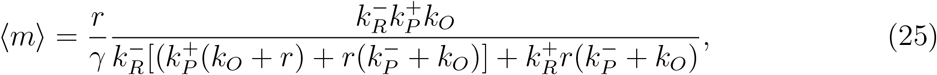

which can be simplified to

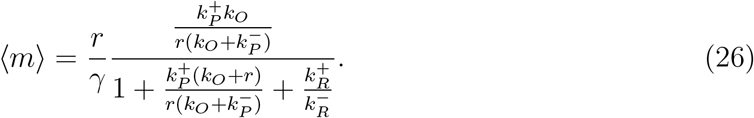

The strategy is to isolate the terms involving the repressor, so that now the fold-change is seen to be simply

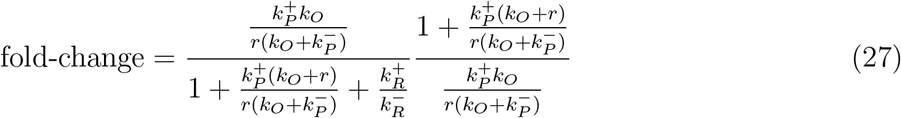

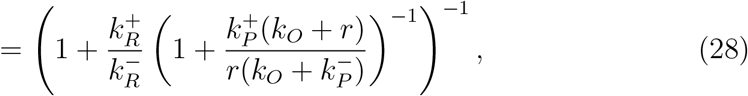

surprisingly reducing to the master curve of Figure 1(D) once again, with 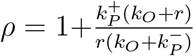.

This example hints that an arbitrarily fine-grained model of downstream transcription steps may still be collapsed to the form of the master curve for the means given in Figure 1(D), so long as the repressor binding is exclusive with transcriptionally active states. We offer this as a conjecture, and we suspect that a careful argument using the King-Altman diagram method [40], [41] might furnish a “proof.” Our focus here is not on full generality but rather to survey an assortment of plausible models for simple repression that have been proposed in the literature.

#### 2.1.6 Nonequilibrium Model Four - “Active” and “Inactive” States

Model 4 in Figure 1(C) is at the core of the theory in [42]. At a glance the cartoon for this model may appear very similar to model 2, and mathematically it is, but the interpretation is rather different. In model 2, we interpreted the third state literally as an RNAP-bound promoter and modeled initiation of a transcript as triggering a promoter state change, making the assumption that an RNAP can only make one transcript at a time. In contrast, in the present model the promoter state does *not* change when a transcript is initiated. So we no longer interpret these states as literally RNAP bound and unbound but instead as coarse-grained “active” and “inactive” states, the details of which we leave unspecified for now. We will comment more on this model below when we discuss Fano factors of models.

Mathematically this model is very similar to models 1 and 2. Like model 1, the matrix *R* describing transcription initiation is diagonal, namely

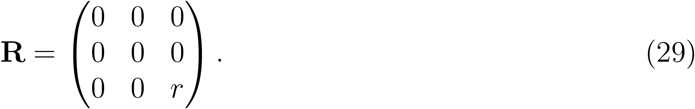

The master equation takes verbatim the same form as it did for model 1, Eq. 12. Meanwhile the promoter transition matrix **K** is the same as Eq. 16 from model 2, although we relabel the rate constants from 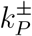 to *k^±^* to reiterate that these are not simply RNAP binding and unbinding rates.

Carrying out the algebra, the mean mRNA can be found to be

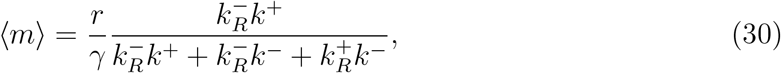

and the fold-change readily follows,

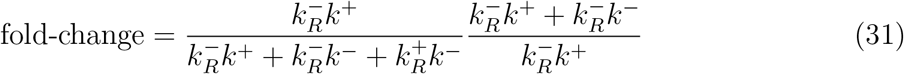

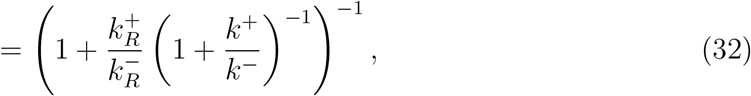

from which we see *ρ* = 1 + *k*^+^/*k^−^* as shown in Figure 1(C).

#### 2.1.7 Nonequilibrium Model Five - Bursty Promoter

The final model we consider shown in Figure 1(C) is an intuitive analog to model 1, with just two states, repressor bound or unbound, and transition rates between them of 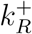 and 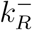. In model 1, when in the unbound state, single mRNA transcripts are produced as a Poisson process with some characteristic rate *r*. The current model by contrast produces, at some Poisson rate *k_i_*, *bursts* of mRNA transcripts. The burst sizes are assumed to be geometrically distributed with a mean burst size *b*, which we will motivate in Section 3 when we derive this model as a certain limiting case of model 4.

From this intuitive picture and by analogy to model 1, then, it should be plausible that the mean mRNA level is

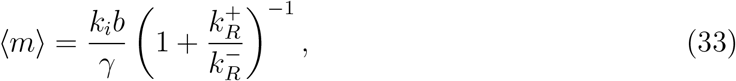

which will turn out to be correct from a careful calculation. For now, we simply note that just like model 1, the fold-change becomes

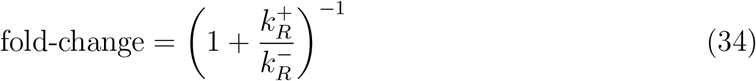

with *ρ* = 1 also like model 1. We will also see later how this model emerges as a natural limit of model 4.

### 2.2 Discussion of Results Across Models for Fold-Changes in Mean Expression

The key outcome of our analysis of the models in Figure 1 is the existence of a master curve shown in Figure 1(D) to which the fold-change predictions of all the models collapse. This master curve is parametrized by only two effective parameters: Δ*F_R_*, which characterizes the number of repressors and their binding strength to the DNA, and *ρ*, which characterizes all other features of the promoter architecture. The key assumption underpinning this result is that no transcription occurs when a repressor is bound to its operator. Note, however, that we are agnostic about the molecular mechanism which achieves this; steric effects are one plausible mechanism, but, for instance, “action-at-a-distance” mediated by kinked DNA due to repressors bound tens or hundreds of nucleotides upstream of a promoter is plausible as well.

Why does the master curve of Figure 1(D) exist at all? This brings to mind the deep questions posed in, e.g., [31] and [43], suggesting we consider multiple plausible models of a system and search for their common patterns to tease out which broad features are and are not important. In our case, the key feature seems to be the exclusive nature of repressor and RNAP binding, which allows the parameter describing the repressor, Δ*F_R_*, to cleanly separate from all other details of the promoter architecture, which are encapsulated in *ρ*. Arbitrary nonequilibrium behavior can occur on the rest of the promoter state space, but it may all be swept up in the effective parameter *ρ*, to which the repressor makes no contribution. We point the interested reader to [44] and [45] for an interesting analysis of similar problems using a graph-theoretic language.

As suggested in [35], we believe this master curve should generalize to architectures with multiple repressor binding sites, as long as the exclusivity of transcription factor binding and transcription initiation is maintained. The interpretation of Δ*F_R_* is then of an effective free energy of all repressor bound states. In an equilibrium picture this is simply given by the log of the sum of Boltzmann weights of all repressor bound states, which looks like the log of a partition function of a subsystem. In a nonequilibrium picture, while we can still mathematically gather terms and give the resulting collection the label Δ*F_R_*, it is unclear if the physical interpretation as an effective free energy makes sense. The problem is that free energies cannot be assigned unambiguously to states out of equilibrium because the free energy change along a generic path traversing the state space is path dependent, unlike at equilibrium. A consequence of this is that, out of equilibrium, Δ*F_R_* is no longer a simple sum of Boltzmann weights. Instead it resembles a restricted sum of King-Altman diagrams [40], [41]. Following the work of Hill [46], it may yet be possible to interpret this expression as an effective free energy, but this remains unclear to us. We leave this an open problem for future work.

If we relax the requirement of exclusive repressor-RNAP binding, one could imagine models in which repressor and RNAP doubly-bound states are allowed, where the repressor’s effect is to reduce the transcription rate rather than setting it to zero. Our results do not strictly apply to such a model, although we note that if the repressor’s reduction of the transcription rate is substantial, such a model might still be well-approximated by one of the models in Figure 1.

One may worry that our “one curve to rule them all” is a mathematical tautology. In fact we *agree* with this criticism if Δ*F_R_* is “just a fitting parameter” and cannot be meaningfully interpreted as a real, physical free energy. An analogy to Hill functions is appropriate here. One of their great strengths and weaknesses, depending on the use they are put to, is that their parameters coarse-grain many details and are generally not interpretable in terms of microscopic models, for deep reasons discussed at length in [31]. By contrast, our master curve claims to have the best of both worlds: a coarse-graining of all details besides the repressor into a single effective parameter *ρ*, while simultaneously retaining an interpretation of Δ*F_R_* as a physically meaningful and interpretable free energy. Our task, then, is to prove or disprove this claim.

How do we test this and probe the theory with fold-change measurements? There is a fundamental limitation in that the master curve is essentially a one-parameter function of Δ*F_R_* + log *ρ*. Worse, there are many *a priori* plausible microscopic mechanisms that could contribute to the value of *ρ*, such as RNAP binding and escape kinetics [38], [39], and/or supercoiling accumulation and release [47], [48], and/or, RNAP clusters analogous to those observed in eukaryotes [49], [50] and recently also observed in bacteria [51]. Even if Δ*F_R_* is measured to high precision, inferring the potential microscopic contributions to *ρ*, buried inside a log no less, from fold-change measurements seems beyond reach. As a statistical inference problem it is entirely nonidentifiable, in the language of [52], Section 4.3.

If we cannot simply infer values of *ρ* from measurements of fold-change, can we perturb some of the parameters that make up *ρ* and measure the change? Unfortunately we suspect this is off-limits experimentally: most of the potential contributors to *ρ* are global processes that affect many or all genes. For instance, changing RNAP association rates by changing RNAP copy numbers, or changing supercoiling kinetics by changing topoisomerase copy numbers, would massively perturb the entire cell’s physiology and confound any determination of *ρ*.

One might instead imagine a bottom-up modeling approach, where we mathematicize a model of what we hypothesize the important steps are and are not, use *in vitro* data for the steps deemed important, and *predict* what *ρ* should be. But again, because of the one-parameter nature of the master curve, many different models will likely make indistinguishable predictions, and without any way to experimentally perturb *in vivo*, there is no clear way to test whether the modeling assumptions are correct.

In light of this, we prefer the view that parameters and rates are not directly comparable between cartoons in Figure 1. Rather, parameters in the simpler cartoons represent coarse-grained combinations of parameters in the finer-grained models. For instance, by equating *ρ* between any two models, one can derive various possible correspondences between the two models’ parameters. Note that these correspondences are clearly not unique, since many possible associations could be made. It then is a choice as to what microscopic interpretations the model-builder prefers for the parameters in a particular cartoon, and as to which coarse-grainings lend intuition and which seem nonsensical. Indeed, since it remains an open question what microscopic features dominate *ρ* (as suggested above, perhaps RNAP binding and escape kinetics [38], [39], or supercoiling accumulation and release [47], [48], or, something more exotic like RNAP clusters [49]–[51]), we are hesitant to put too much weight on any one microscopic interpretation of model parameters that make up *ρ*.

One possible tuning knob to probe *ρ* that would not globally perturb the cell’s physiology is to manipulate RNAP binding sites. Work such as [53] has shown that models of sequence-dependent RNAP affinity can be inferred from data, and the authors of [54] showed that the model of [53] has predictive power by using the model to *design* binding sites of a desired affinity. But for our purposes, this begs the question: the authors of [53] *assumed* a particular model (essentially our 3-state equilibrium model but without the repressor), so it is unclear how or if such methods can be turned around to *compare* different models of promoter function.

Another possible route to dissect transcription details without a global perturbation would be to use phage polymerase with phage-specific promoters. While such results would carry some caveats, e.g., whether the repression of the phage polymerase is a good analog to the repression of the native RNAP, it could nevertheless be worthy of consideration.

We have already pointed out that the master curve of Figure 1 is essentially a one-parameter model, the one parameter being Δ*F_R_* + log *ρ*. By now the reader may be alarmed as to how can we even determine Δ*F_R_* and *ρ* independently of each other, never mind shedding a lens on the internal structure of *ρ* itself. A hint is provided by the weak promoter approximation, invoked repeatedly in prior studies [14], [32], [33] of simple repression using the 3-state equilibrium model in Figure 1(B). In that picture, the weak promoter approximation means 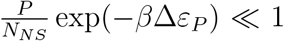, meaning therefore *ρ* ≈ 1. This approximation can be well justified on the basis of the number of RNAP and *σ* factors per cell and the strength of binding of RNAP to DNA at weak promoters. This is suggestive, but how can we be sure that *ρ* is not, for instance, actually 10^2^ and that Δ*F_R_* hides a compensatory factor? A resolution is offered by an independent inference of *ρ* in the absence of repressors. This was done in [42] by fitting nonequilibrium model 4 in Figure 1(C), with zero repressor (looking ahead, this is equivalent to model 4 in Figure 2(A)), to single-cell mRNA counts data from [34]. This provided a determination of *k*^+^ and *k^−^*, from which their ratio is estimated to be no more than a few 10^−1^ and possibly as small as 10^−2^.

**Figure 2.**
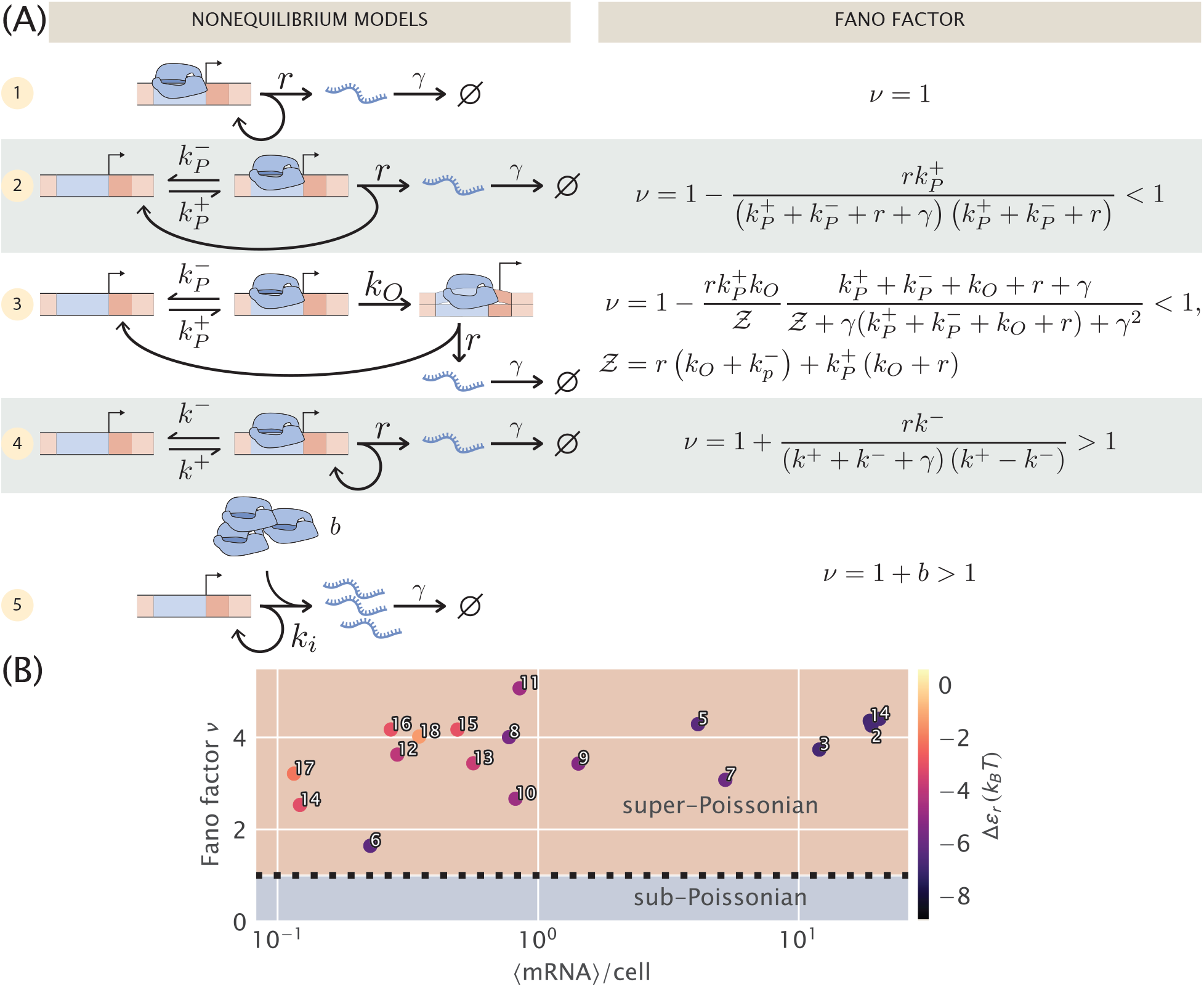
Comparison of different models for noise in the constitutive promoter. (A) The left column depicts various plausible models for the dynamics of constitutive promoters. In model (1), transcripts are produced in a Poisson process [36], [57]. Model (2) features explicit modeling of RNAP binding/unbinding kinetics [58]. Model (3) is a more detailed generalization of model (2), treating transcription initiation as a multi-step process proceeding through closed and open complexes [38]. Model (4) is somewhat analogous to (2) except with the precise nature of active and inactive states left ambiguous [19], [23], [42]. Finally, model (5) can be viewed as a certain limit of model (4) in which transcripts are produced in bursts, and initiation of bursts is a Poisson process. The right column shows the Fano factor *ν* (variance/mean) for each model. Note especially the crucial diagnostic: (2) and (3) have *ν* strictly below 1, while only for (4) and (5) can *ν* exceed 1. Models with Fano factors ≤ 1 cannot produce the single-cell data observed in part (B) without introducing additional assumptions and model complexity. (B) Data from [36]. Mean mRNA count vs. Fano factor (variance/mean) for different promoters as determined with single-molecule mRNA Fluorescence *in situ* Hybridization. The colorbar indicates the predicted binding affinity of RNAP to the promoter sequence as determined in [54]. Numbers serve for cross comparison with data presented in Figure 3.

The realization that *ρ* ≈ 1 to an excellent approximation, *independent* of which model in Figure 1 one prefers, goes a long way towards explaining the surprising success of equilibrium models of simple repression. Even though our 2- and 3-state models get so many details of transcription wrong, it does not matter because fold-change is a cleverly designed ratio. Since *ρ* subsumes all details except the repressor, and log *ρ* ≈ 0, fitting these simple models to fold-change measurements can still give a surprisingly good estimate of repressor binding energies. So the ratiometric construction of fold-change fulfills its intended purpose of canceling out all features of the promoter architecture except the repressor itself. Nevertheless it is perhaps surprising how effectively it does so: *a priori*, one might not have expected *ρ* to be quite so close to 1.

We would also like to highlight the relevance of [55] here. Landman et. al. reanalyzed and compared *in vivo* and *in vitro* data on the lacI repressor’s binding affinity to its various operator sequences. (The *in vivo* data was from, essentially, fitting our master curve to expression measurements.) They find broad agreement between the *in vitro* and *in vivo* values. This reinforces the suspicion that the equilibrium Δ*ε_R_* repressor binding energies do in fact represent real physical free energies. Again, *a priori* this did not have to be the case, even knowing that *ρ* ≈ 1.

In principle, if Δ*F_R_* can be measured to sufficient precision, then deviations from *ρ* = 1 become a testable matter of experiment. In practice, it is probably unrealistic to measure repressor rates 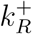 or 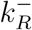 or fold-changes in expression levels (and hence Δ*ε_R_*) precisely enough to detect the expected tiny deviations from *ρ* = 1. We can estimate the requisite precision in Δ*F_R_* to resolve a given Δ*ρ* by noting, since *ρ* ≈ 1, that log(1 + Δ*ρ*) ≈ Δ*ρ*, so Δ(Δ*F_R_*) ≈ Δ*ρ*. Suppose we are lucky and Δ*ρ* is ~ 0.1, on the high end of our range estimated above. A determination of Δ*ε_R_/k_B_T* with an uncertainty of barely 0.1 was achieved in the excellent measurements of [32], so this requires a very difficult determination of Δ*F_R_* for a very crude determination of *ρ*, which suggests, to put it lightly, this is not a promising path to pursue experimentally. It is doubtful that inference of repressor kinetic rates would be any easier.

Moving forward, we have weak evidence supporting the interpretation of Δ*F_R_* as a physically real free energy [55] and other work casting doubt [56]. How might we resolve the confusion? If there is no discriminatory power to test the theory and distinguish the various models with measurements of fold-changes in means, how do we probe the theory? Clearly to discriminate between the nonequilibrium models in Figure 1, we need to go beyond means to ask questions about kinetics, noise and even full distributions of mRNA copy numbers over a population of cells. If the “one-curve-to-rule-them-all” is more than a mathematical tautology, then the free energy of repressor binding inferred from fold-change measurements should agree with repressor binding and unbinding rates. In other words, the equilibrium and nonequilibrium definitions of Δ*F_R_* must agree, meaning

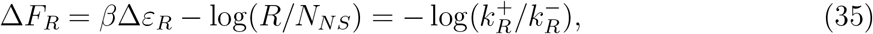

must hold, where *β*Δ*ε_R_* is inferred from the master curve fit to fold-change measurements, and 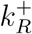 and 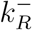 are inferred in some orthogonal manner. Single molecule measurements such as from [56] have directly observed these rates, and in the remainder of this work we explore a complementary approach: inferring repressor binding and unbinding rates 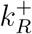 and 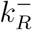 from single-molecule measurements of mRNA population distributions.

## 3 Beyond Means in Gene Expression

In this section, our objective is to explore the same models considered in the previous section, but now with reference to the the question of how well they describe the distribution of gene expression levels, with special reference to the variance in these distributions. To that end, we repeat the same pattern as in the previous section by examining the models one by one. In particular we will focus on the Fano factor, defined as the variance/mean. This metric serves as a powerful discriminatory tool from the null model that the steady-state mRNA distribution must be Poisson, giving a Fano factor of one.

### 3.1 Kinetic models for unregulated promoter noise

Before we can tackle simple repression, we need an adequate phenomenological model of constitutive expression. The literature abounds with options from which we can choose, and we show several potential kinetic models for constitutive promoters in Figure 2(A). Let us consider the suitability of each model for our purposes in turn.

#### 3.1.1 Noise in the Poisson Promoter Model

The simplest model of constitutive expression that we can imagine is shown as model 1 in Figure 2(A) and assumes that transcripts are produced as a Poisson process from a single promoter state. This is the picture from Jones et. al. [36] that was used to interpret a systematic study of gene expression noise over a series of promoters designed to have different strengths. This model insists that the “true” steady-state mRNA distribution is Poisson, implying the Fano factor *ν* must be 1. In [36], the authors carefully attribute measured deviations from Fano = 1 to intensity variability in fluorescence measurements, gene copy number variation, and copy number fluctuations of the transcription machinery, e.g., RNAP itself. In this picture, the master equation makes no appearance, and all the corrections to Poisson behavior are derived as additive corrections to the Fano factor. For disproving the “universal noise curve” from So et. al. [59], this picture was excellent. It is appealing in its simplicity, with only two parameters, the initiation rate *r* and degradation rate *γ*. Since *γ* is independently known from other experiments, and the mean mRNA copy number is *r/γ*, *r* is easily inferred from data. In other words, the model is not excessively complex for the data at hand. But for many interesting questions, for instance in the recent work [42], knowledge of means and variances alone is insufficient, and a description of the full distribution of molecular counts is necessary. For this we need a (slightly) more complex model than model 1 that would allow us to incorporate the non-Poissonian features of constitutive promoters directly into a master equation formulation.

#### 3.1.2 Noise in the Two-State Promoter, RNAP Bound or Unbound

Our second model of constitutive transcription posits an architecture in which the promoter is either empty or bound by RNAP [27], [58]. Here, as shown in model 2 of Figure 2(A), transcription initiation results in a state transition from the bound to the unbound state, reflecting the microscopic reality that an RNAP that has begun to elongate a transcript is no longer available at the start site to begin another. As shown in Appendix S1, the Fano factor in this model is given by

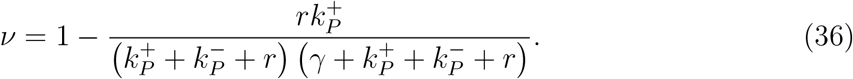

The problem with this picture is that the Fano factor is always < 1. To make contact with the experimental reality of *ν* > 1 as shown in Figure 2(B), clearly some corrections will be needed. While this model adds an appealing element of microscopic reality, we are forced to reject it as the additional complexity is unable to capture the phenomenology of interest. Obviously the promoter state does in fact proceed through cycles of RNAP binding, initiating, and elongating, but it seems that the super-Poissonian noise in mRNA copy number we want to model must be governed by other features of the system.

#### 3.1.3 Noise in the Three-State Promoter, Multistep Transcription Initiation and Escape

How might we remedy the deficits of model 2? It is known [39] that once RNAP initially binds the promoter region, a multitude of distinct steps occur sequentially before RNAP finally escapes into the elongation phase. Perhaps adding some of this mechanistic detail as shown in model 3 of Figure 2(A) might rescue the previous model. The next simplest refinement of that model could consider open complex formation and promoter escape as separate steps rather than as a single effective step. In other words, we construct model 3 by adding a single extra state to model 2, and we will label the two RNAP-bound states as the closed and open complexes, despite the true biochemical details certainly being more complex. For example, earlier work extended this model by adding an additional repressor bound state and did not explicitly consider the limit with no repressor that we analyze here [38]. Again, our goal here is not a complete accounting of all the relevant biochemical detail; this is an exploratory search for the important features that a model needs to include to square with the known experimental reality of constitutive expression.

Unfortunately, as hinted at in earlier work [38], this model too has Fano factor *ν* < 1. We again leave the algebraic details for Appendix S1 and merely state the result that

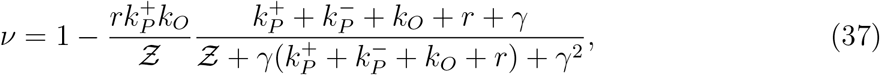

where we defined 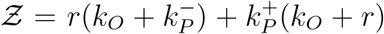 for notational tidiness. This is necessarily less than 1 for arbitrary rate constants.

In fact, we suspect *any* model in which transcription proceeds through a multistep cycle must necessarily have *ν* < 1. The intuitive argument compares the waiting time distribution to traverse the cycle with the waiting time for a Poisson promoter (model 1) with the same mean time. The latter is simply an exponential distribution. The former is a convolution of multiple exponentials, and intuitively the waiting time distribution for a multistep process should be more peaked with a smaller fractional width than a single exponential with the same mean. A less disperse waiting time distribution means transcription initations are more uniformly distributed in time relative to a Poisson process. Hence the distribution of mRNA over a population of cells should be less variable compared to Poisson, giving *ν* < 1. (In Appendix S1 we present a more precise version of the intuitive arguments in this paragraph.) Regardless of the merits of this model in describing the noise properties of constitutive transcription initiation, it ultimately fails the simplest quantitative feature of the data, namely that the Fano factor > 1 and hence we must discard this mechanistic picture and search elsewhere.

#### 3.1.4 Noise in a Two-State Promoter with “Active” and “Inactive” States

Inspired by [42], we next revisit an analog of model 2 in Figure 2(A), but as with the analogous models considered in Section 2, the interpretation of the two states is changed. Rather than explicitly viewing them as RNAP bound and unbound, we view them as “active” and “inactive,” which are able and unable to initiate transcripts, respectively. We are noncommittal as to the microscopic details of these states.

One interpretation [47], [48] for the active and inactive states is that they represent the promoter’s supercoiling state: transitions to the inactive state are caused by accumulation of positive supercoiling, which inhibits transcription, and transitions back to “active” are caused by gyrase or other topoisomerases relieving the supercoiling. This is an interesting possibility because it would mean the timescale for promoter state transitions is driven by topoisomerase kinetics, not by RNAP kinetics. From in vitro measurements, the former are suggested to be of order minutes [47]. Contrast this with model 2, where the state transitions are assumed to be governed by RNAP, which, assuming a copy number per cell of order 10^3^, has a diffusion-limited association rate *k_on_* ~ 10^2^ s^−1^ to a target promoter. Combined with known *K_d_*’s of order *μ*M, this gives an RNAP dissociation rate *k_off_* of order 10^2^ s^−1^. As we will show below, however, there are some lingering puzzles with interpreting this supercoiling hypothesis, so we leave it as a speculation and refrain from assigning definite physical meaning to the two states in this model.

Intuitively one might expect that, since transcripts are produced as a Poisson process only when the promoter is in one of the two states in this model, transcription initiations should now be “bunched” in time, in contrast to the “anti-bunching” of models 2 and 3 above. One might further guess that this bunching would lead to super-Poissonian noise in the mRNA distribution over a population of cells. Indeed, as shown in Appendix S1, a calculation of the Fano factor produces

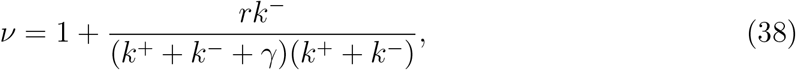

which is strictly greater than 1, verifying the above intuition. Note we have dropped the *P* label on the promoter switching rates to emphasize that these very likely do not represent kinetics of RNAP itself. This calculation can also be sidestepped by noting that the model is mathematically equivalent to the simple repression model from [36], with states and rates relabeled and reinterpreted.

How does this model compare to model 1 above? In model 1, all non-Poisson features of the mRNA distribution were handled as extrinsic corrections. By contrast, here the 3 parameter model is used to fit the full mRNA distribution as measured in mRNA FISH experiments. In essence, all variability in the mRNA distribution is regarded as “intrinsic,” arising either from stochastic initiation or from switching between the two coarse-grained promoter states. The advantage of this approach is that it fits neatly into the master equation picture, and the parameters thus inferred can be used as input for more complicated models with regulation by transcription factors.

While this seems promising, there is a major drawback for our purposes which was already uncovered by the authors of [42]: the statistical inference problem is nonidentifiable, in the sense described in Section 4.3 of [52]. What this means is that it is impossible to infer the parameters *r* and *k^−^* from the single-cell mRNA counts data of [36] (as shown in Fig. S2 of [42]). Rather, only the ratio *r/k^−^* could be inferred. In that work, the problem was worked around with an informative prior on the ratio *k^−^/k*^+^. That approach is unlikely to work here, as, recall, our entire goal in modeling constitutive expression is to use it as the basis for a yet more complicated model, when we add on repression. But adding more complexity to a model that is already poorly identified is a fool’s errand, so we will explore one more potential model.

#### 3.1.5 Noise Model for One-State Promoter with Explicit Bursts

The final model we consider is inspired by the failure mode of model 4. The key observation above was that, as found in [42], only two parameters, *k*^+^ and the ratio *r/k^−^*, could be directly inferred from the single-cell mRNA data from [36]. So let us take this seriously and imagine a model where these are the only two model parameters. What would this model look like?

To develop some intuition, consider model 4 in the limit *k*^+^ ≪ *k^−^* ≲ *r*, which is roughly satisfied by the parameters inferred in [42]. In this limit, the system spends the majority of its time in the inactive state, occasionally becoming active and making a burst of transcripts. This should call to mind the well-known phenomenon of transcriptional bursting, as reported in, e.g., [47], [48], [60]. Let us make this correspondence more precise. The mean dwell time in the active state is 1/*k^−^*. While in this state, transcripts are produced at a rate *r* per unit time. So on average, *r/k^−^* transcripts are produced before the system switches to the inactive state. Once in the inactive state, the system dwells there for an average time 1/*k*^+^ before returning to the active state and repeating the process. *r/k^−^* resembles an average burst size, and 1/*k*^+^ resembles the time interval between burst events. More precisely, 1/*k*^+^ is the mean time between the end of one burst and the start of the next, whereas 1/*k*^+^ +1/*k^−^* would be the mean interval between the start of two successive burst events, but in the limit *k*^+^ ≪ *k^−^*, 1/*k*^+^ + 1/*k^−^* ≈ 1/*k*^+^. Note that this limit ensures that the waiting time between bursts is approximately exponentially distributed, with mean set by the only timescale left in the problem, 1/*k*^+^.

Let us now verify this intuition with a precise derivation to check that *r/k^−^* is in fact the mean burst size and to obtain the full burst size distribution. Consider first a constant, known dwell time *T* in the active state. Transcripts are produced at a rate *r* per unit time, so the number of transcripts *n* produced during *T* is Poisson distributed with mean *rT*, i.e.,

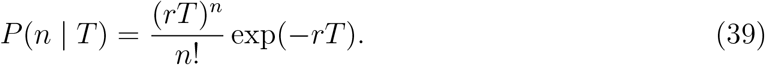

Since the dwell time *T* is unobservable, we actually want *P* (*n*), the dwell time distribution with no conditioning on *T*. Basic rules of probability theory tell us we can write *P* (*n*) in terms of *P* (*n* | *T*) as

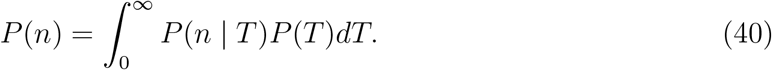

But we know the dwell time distribution *P* (*T*), which is exponentially distributed according to

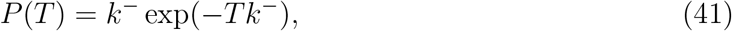

so *P* (*n*) can be written as

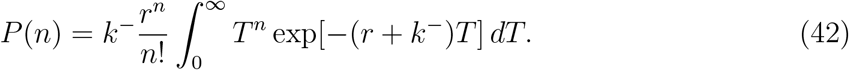

A standard integral table shows 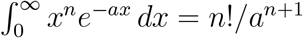, so

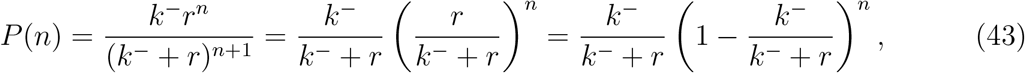

which is exactly the geometric distribution with standard parameter *θ* ≡ *k^−^/*(*k^−^* + *r*) and domain *n* ∈ {0, 1, 2,…}. The mean of the geometric distribution, with this convention, is

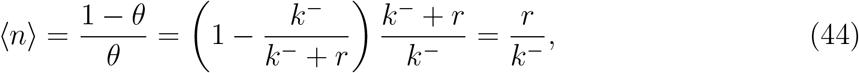

exactly as we guessed intuitively above.

So in taking the limit *r, k^−^* → ∞, *r/k^−^* ≡ *b*, we obtain a model which effectively has only a single promoter state, which initiates bursts at rate *k*^+^ (transitions to the active state, in the model 4 picture). The master equation for mRNA copy number *m* as derived in Appendix S1 takes the form

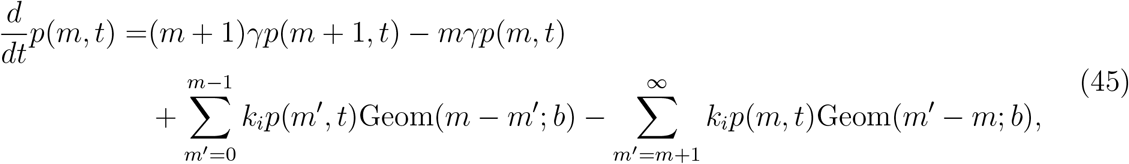

where we use *k_i_* to denote the burst initiation rate, Geom(*n*; *b*) is the geometric distribution with mean *b*, i.e., 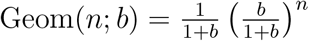 (with domain over nonnegative integers as above). The first two terms are the usual mRNA degradation terms. The third term enumerates all ways the system can produce a burst of transcripts and arrive at copy number *m*, given that it had copy number *m′* before the burst. The fourth term allows the system to start with copy number *m*, then produce a burst and end with copy number *m′*. In fact this last sum has trivial *m′* dependence and simply enforces normalization of the geometric distribution. Carrying it out we have

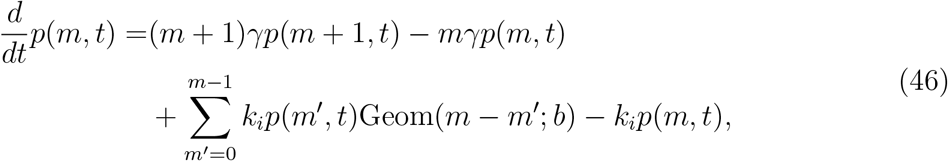

We direct readers again to Appendix S2 for further details. This improves on model 4 in that now the parameters are easily inferred, as we will see later, and have clean interpretations. The non-Poissonian features are attributed to the empirically well-established phenomenological picture of bursty transcription.

The big approximation in going from model 4 to 5 is that a burst is produced instantaneously rather than over a finite time. If the true burst duration is not short compared to transcription factor kinetic timescales, this could be a problem in that mean burst size in the presence and absence of repressors could change, rendering parameter inferences from the constitutive case inappropriate. Let us make some simple estimates of this.

Consider the time delay between the first and final RNAPs in a burst initiating transcription (*not* the time to complete transcripts, which potentially could be much longer.) If this timescale is short compared to the typical search timescale of transcription factors, then all is well. The estimates from deHaseth et. al. [39] put RNAP’s diffusion-limited on rate around ~ few × 10^−2^ nM^−1^ s^−1^ and polymerase loading as high as 1 s^−1^. Then for reasonable burst sizes of < 10, it is reasonable to guess that bursts might finish initiating on a timescale of tens of seconds or less (with another 30-60 sec to finish elongation, but that does not matter here). A transcription factor with typical copy number of order 10 (or less) would have a diffusion-limited association rate of order (10 sec)^−1^ [56]. Higher copy number TFs tend to have many binding sites over the genome, which should serve to pull them out of circulation and keep their effective association rates from rising too large. Therefore, there is *perhaps* a timescale separation possible between transcription factor association rates and burst durations, but this assumption could very well break down, so we will have to keep it in mind when we infer repressor rates from the Jones et. al. single-cell mRNA counts data later [36].

In reflecting on these 5 models, the reader may feel that exploring a multitude of potential models just to return to a very minimal phenomenological model of bursty transcription may seem highly pedantic. But the purpose of the exercise was to examine a host of models from the literature and understand why they are insufficient, one way or another, for our purposes. Along the way we have learned that the detailed kinetics of RNAP binding and initiating transcription are probably irrelevant for setting the population distribution of mRNA. The timescales are simply too fast, and as we will see later in Figures 3 and 4, the noise seems to be governed by slower timescales. Perhaps in hindsight this is not surprising: intuitively, the degradation rate *γ* sets the fundamental timescale for mRNA dynamics, and any other processes that substantially modulate the mRNA distribution should not differ from *γ* by orders of magnitude.

**Figure 3.**
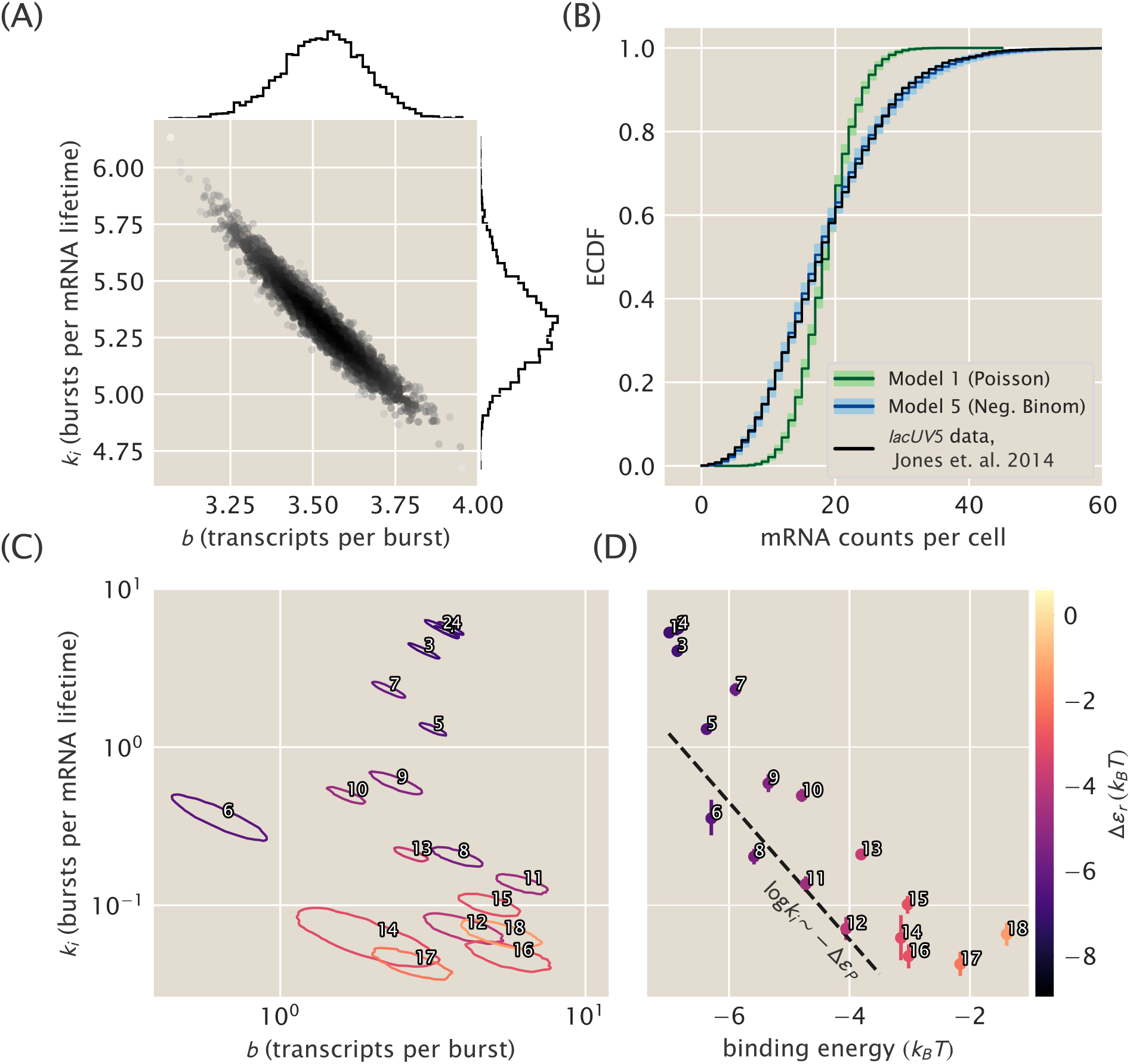
Constitutive promoter posterior inference and model comparison. (A) The joint posterior density of model 5, the bursty promoter with negative binomially-distributed steady state, is plotted with MCMC samples. 1D marginal probability densities are plotted as flanking histograms. The model was fit on *lacUV5* data from [36]. (B) The empirical distribution function (ECDF) of the observed population distribution of mRNA transcripts under the control of a constitutive *lacUV5* promoter is shown in black. The median posterior predictive ECDFs for models (1), Poisson, and (5), negative binomial, are plotted in dark green and dark blue, respectively. Lighter green and blue regions enclose 95% of all posterior predictive samples from their respective models. Model (1) is in obvious contradiction with the data while model (5) is not. Single-cell mRNA count data is again from [36]. (C) Joint posterior distributions for burst rate *k_i_* and mean burst size *b* for 18 unregulated promoters from [36]. Each contour indicates the 95% highest posterior probability density region for a particular promoter. Note that the vertical axis is shared with (D). (D) Plots of the burst rate *k_i_* vs. the binding energy for each promoter as predicted in [54]. The dotted line shows the predicted slope according to Eq. 55, described in text. Each individual promoter is labeled with a unique number in both (C) and (D) for cross comparison and for comparison with Figure 2(B).

**Figure 4.**
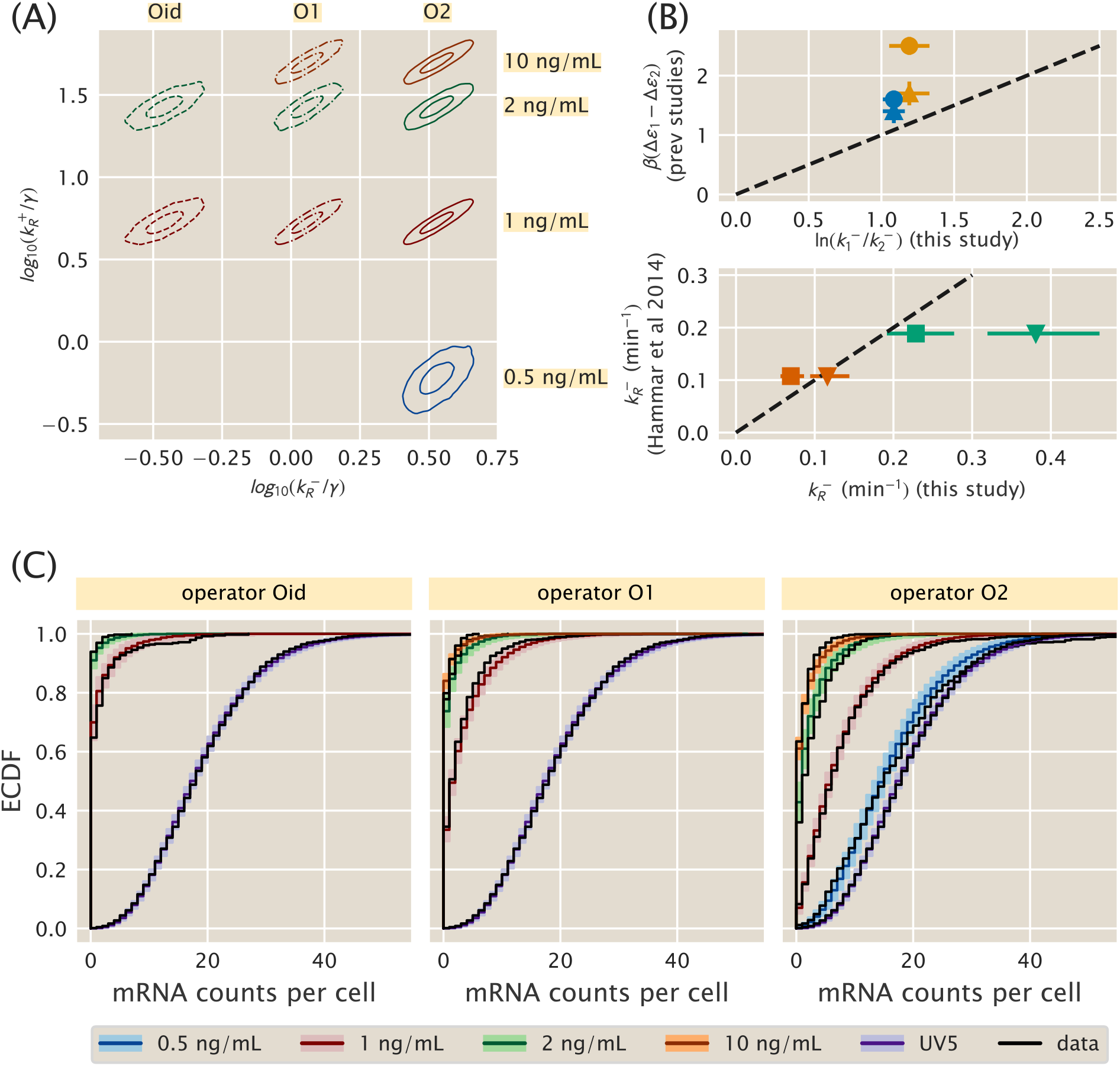
Simple repression parameter inference and comparison. (A) Contours which enclose 50% and 95% of the posterior probability mass are shown for each of several 2D slices of the 9D posterior distribution. The model assumes one unbinding rate for each operator (Oid, O1, O2) and one binding rate for each aTc induction concentration (corresponding to an unknown mean repressor copy number). (B, upper) Ratios of our inferred unbinding rates are compared with operator binding energy differences measured by Garcia and Phillips [33] (triangles) and Razo-Mejia et. al. [32] (circles). Blue glyphs compare O2-O1, while orange compare O1-Oid. Points with perfect agreement would lie on the dotted line. (B, lower) Unbinding rates for O1 (cyan) and Oid (red) inferred in this work are compared with single-molecule measurements from Hammar et. al. [56]. We plot the comparison assuming illustrative mRNA lifetimes of *γ^−^*^1^ = 3 min (triangles) or *γ^−^*^1^ = 5 min (squares). Dotted line is as in upper panel. (C) Theory-experiment comparison are shown for each of the datasets used in the inference of the model in (A). Observed single-molecule mRNA counts data from [36] are plotted as black lines. The median of the randomly generated samples for each condition is plotted as a dark colored line. Lighter colored bands enclose 95% of all samples for a given operator/repressor copy number pair. The unregulated promoter, *lacUV5*, is shown with each as a reference.

## 4 Finding the “right” model: Bayesian parameter inference

In this section of the paper, we continue our program of providing one complete description of the entire broad sweep of studies that have been made in the context of the repressor-operator model, dating all the way back to the original work of Jacob and Monod and including the visionary quantitative work of Müller-Hill and collaborators [30] and up to more recent studies [33]. In addition, the aim is to reconcile the equilibrium and non-equilibrium perspectives that have been brought to bear on this problem. From Section 2, this reconciliation depends on a key quantitative question as codified by Eq. 35: does the free energy of repressor binding, as described in the equilibrium models and indirectly inferred from gene expression measurements, agree with the corresponding values of repressor binding and unbinding rates in the non-equilibrium picture, measured or inferred more directly? In this section we tackle the statistical inference problem of inferring these repressor rates from single-cell mRNA counts data. But before we can turn to the full case of simple repression, we must choose an appropriate model of the constitutive promoter and infer the parameter values in that model. This is the problem we address first.

### 4.1 Parameter inference for constitutive promoters

From consideration of Fano factors in the previous section, we suspect that model 5 in Figure 2(A), a one-state bursty model of constitutive promoters, achieves the right balance of complexity and simplicity, by allowing both Fano factor *ν* > 1, but also by remedying, by design, the problems of parameter degeneracy that model 4 in Figure 2 suffered [42]. Does this stand up to closer scrutiny, namely, comparison to full mRNA distributions rather than simply their moments? We will test this thoroughly on single-cell mRNA counts for different unregulated promoters from Jones et. al. [36].

It will be instructive, however, to first consider the Poisson promoter, model 1 in Figure 2. As we alluded to earlier, since the Poisson distribution has a Fano factor *ν* strictly equal to 1, and all of the observed data in Figure 2(B) has Fano factor *ν* > 1, we might already suspect that this model is incapable of fitting the data. We will verify that this is in fact the case. Using the same argument we can immediately rule out models 2 and 3 from Figure 2(A). These models have Fano factors *ν* ≤ 1 meaning they are underdispersed relative to the Poisson distribution. We will also not explicitly consider model 4 from Figure 2(A) since it was already thoroughly analyzed in [42], and since model 5 can be viewed as a special case of it.

Our objective for this section will then be to assess whether or not model 5 is quantitatively able to reproduce experimental data. In other words, if our claim is that the level of coarse graining in this model is capable of capturing the relevant features of the data, then we should be able to find values for the model parameters that can match theoretical predictions with single-molecule mRNA count distributions. A natural language for this parameter inference problem is that of Bayesian probability. We will then build a Bayesian inference pipeline to fit the model parameters to data. To gain intuition on how this analysis is done we will begin with the “wrong” model 1 in Figure 2(A). We will use the full dataset of single-cell mRNA counts from [36] used in Figure 2(B).

#### 4.1.1 Model 1: Poisson promoter

For this model the master equation of interest is Eq. 10 with repressor set to zero, i.e.,

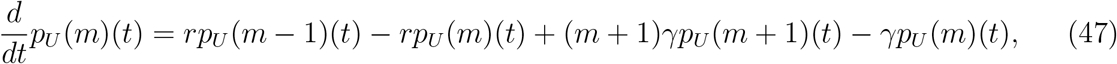

whose steady-state solution is given by a Poisson distribution with parameter *λ* ≡ *r/γ* [57]. The goal of our inference problem is then to find the probability distribution for the parameter value *λ* given the experimental data. By Bayes’ theorem this can be written as

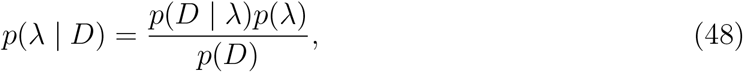

where *D* = {*m*_1_, *m*_2_,…, *m_N_*} are the single-cell mRNA experimental counts. As is standard we will neglect the denominator *p*(*D*) on the right hand side since it is independent of *λ* and serves only as a normalization factor.

The steady-state solution for the master equation defines the likelihood term for a single cell *p*(*m* | *λ*). What this means is that for a given choice of parameter *λ*, under model 1 of Figure 2(A), we expect to observe *m* mRNAs in a single cell with probability

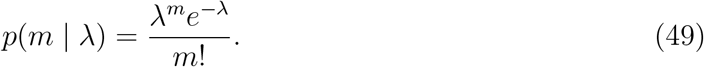

Assuming each cell’s mRNA count in our dataset is independent of others, the likelihood of the full inference problem *p*(*D* | *λ*) is simply a product of the single cell likelihoods given by Eq. 49 above, so

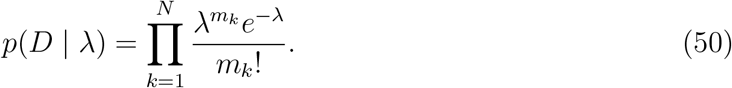

To proceed we need to specify a prior distribution *p*(*λ*). In this case we are extremely data-rich, as the dataset from Jones et. al [36] has of order 1000-3000 single-cell measurements for each promoter, so our choice of prior matters little here, as long as it is sufficiently broad. For details on the prior selection we refer the reader to Appendix S3. For our purpose here it suffices to specify that we use as prior a Gamma distribution. This particular choice of prior introduces two new parameters, *α* and *β*, which parametrize the gamma distribution itself, which we use to encode the range of *λ* values we view as reasonable. Recall *λ* is the mean steady-state mRNA count per cell, which *a priori* could plausibly be anywhere from 0 to a few hundred. *α* = 1 and *β* = 1/50 achieve this, since the gamma distribution is strictly positive with mean *α/β* and standard deviation 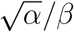.

As detailed in Appendix S3 this particular choice of prior is known as the *conjugate* prior for a Poisson likelihood. Conjugate priors have the convenient properties that a closed form exists for the posterior distribution *p*(*λ* | *D*) - unusual in Bayesian inference problems - and the closed form posterior takes the same form as the prior. For our case of a Poisson distribution likelihood with its Gamma distribution conjugate prior, the posterior distribution is also a Gamma distribution [52]. Specifically the two parameters *α′* and *β′* for this posterior distribution take the form 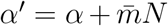 and *β′* = *β* + *N*, where we defined the sample mean 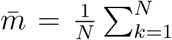 for notational convenience, and *N* is the number of cells in our dataset. Furthermore, given that *N* is 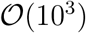 and 〈*m*〉 ≳ 0.1 for all promoters measured in [36] our data easily overwhelms the choice of prior, and allows us to approximate the Gamma distribution with a Gaussian distribution with mean 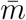 and variance 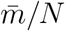 with marginal errors. As an example with real numbers, for the *lacUV5* promoter, Jones et. al [36] measured 2648 cells with an average mRNA count per cell of 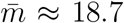. For this case our posterior distribution *P* (*λ* | *D*) would be a Gaussian distribution with mean *μ* = 18.7, and a standard deviation *σ* ≈ 0.08. This suggests we have inferred our model’s one parameter to a precision of order 1%.

We remind the reader that we began this section claiming that the Poisson model was “wrong” since it could not reproduce features of the data such as a Fano factor > 1. The fact that we obtain such a narrow posterior distribution for our parameter *P* (*λ* | *D*) does not equate to the model being adequate to describe the data. What this means is that given the data *D*, only values in a narrow range are remotely plausible for the parameter *λ*, but a narrow posterior distribution does not necessarily mean the model accurately depicts reality. As we will see later in Figure 3 after exploring the bursty promoter model, indeed the correspondence when contrasting the Poisson model with the experimental data is quite poor.

#### 4.1.2 Model 5 - Bursty promoter

Let us now consider the problem of parameter inference for model five from Figure 1(C). As derived in Appendix S2, the steady-state mRNA distribution in this model is a negative binomial distribution, given by

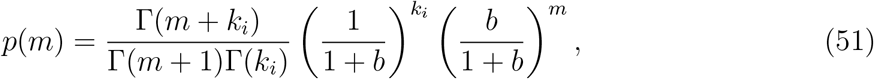

where *b* is the mean burst size and *k_i_* is the burst rate in units of the mRNA degradation rate *γ*. As sketched earlier, to think of the negative binomial distribution in terms of an intuitive “story,” in the precise meaning of [61], we imagine the arrival of bursts as a Poisson process with rate *k_i_*, where each burst has a geometrically-distributed size with mean size *b*.

As for the Poisson promoter model, this expression for the steady-state mRNA distribution is exactly the likelihood we want to use when stating Bayes theorem. Again denoting the single-cell mRNA count data as *D* = {*m*_1_, *m*_2_,…, *m_N_*}, here Bayes’ theorem takes the form

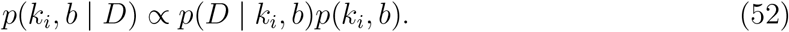

We already have our likelihood – the product of *N* negative binomials as Eq. 51 – so we only need to choose priors on *k_i_* and *b*. For the datasets from [36] that we are analyzing, as for the Poisson promoter model above we are still data-rich so the prior’s influence remains weak, but not nearly as weak because the dimensionality of our model has increased from one parameter to two. Details on the arguments behind our prior distribution selection are left for Appendix S3. We state here that the natural scale to explore these parameters is logarithmic. This is commonly the case for parameters for which our previous knowledge based on our domain expertise spans several orders of magnitude. For this we chose log-normal distributions for both *k_i_* and *b*. Details on the mean and variance of these distributions can be found in Appendix S3.

We carried out Markov-Chain Monte Carlo (MCMC) sampling on the posterior of this model, starting with the constitutive *lacUV5* dataset from [36]. The resulting MCMC samples are shown in Figure 3(A). In contrast to the active/inactive constitutive model considered in [42] (nonequilibrium model 4 in Figure 2(A)), this model is well-identified with both parameters determined to a fractional uncertainty of 5-10%. The strong correlation reflects the fact that their product sets the mean of the mRNA distribution, which is tightly constrained by the data, but there is weak “sloppiness” [62] along a set of values with a similar product.

Having found the model’s posterior to be well-identified as with the Poisson promoter, the next step is to compare both models with experimental data. To do this for the case of the bursty promoter, for each of the parameter samples shown in Figure 3(A) we generated negative bionomial-distributed mRNA counts. As MCMC samples parameter space proportionally to the posterior distribution, this set of random samples span the range of possible values that we would expect given the correspondence between our theoretical model and the experimental data. A similar procedure can be applied to the Poisson promoter. To compare so many samples with the actual observed data, we can use empirical cumulative distribution functions (ECDF) of the distribution quantiles. This representation is shown in Figure 3(B). In this example, the median for each possible mRNA count for the Poisson distribution is shown as a dark green line, while the lighter green contains 95% of the randomly generated samples. This way of representing the fit of the model to the data gives us a sense of the range of data we might consider plausible, under the assumption that the model is true. For this case, as we expected given our premise of the Poisson promoter being wrong, it is quite obvious that the observed data, plotted in black is not consistent with the Poisson promoter model. An equivalent plot for the bursty promoter model is shown in blue. Again the darker tone shows the median, while the lighter color encompasses 95% of the randomly generated samples. Unlike the Poisson promoter model, the experimental ECDF closely tracks the posterior predictive ECDF, indicating this model is actually able to generate the observed data and increasing our confidence that this model is sufficient to parametrize the physical reality of the system.

The commonly used promoter sequence *lacUV5* is our primary target here, since it forms the core of all the simple repression constructs of [36] that we consider in Section 4.2. Nevertheless, we thought it wise to apply our bursty promoter model to the other 17 unregulated promoters available in the single-cell mRNA count dataset from [36] as a test that the model is capturing the essential phenomenology. If the model fit well to all the different promoters, this would increase our confidence that it would serve well as a foundation for inferring repressor kinetics later in Section 4.2. Conversely, were the model to fail on more than a couple of the other promoters, it would give us pause.

Figure 3(C) shows the results, plotting the posterior distribution from individually MCMC sampling all 18 constitutive promoter datasets from [36]. To aid visualization, rather than plotting samples for each promoter’s posterior as in Figure 3(A), for each posterior we find and plot the curve that surrounds the 95% highest probability density region. What this means is that each contour encloses approximately 95% of the samples, and thus 95% of the probability mass, of its posterior distribution. Theory-experiment comparisons, shown in Figure S5 in Appendix S3, display a similar level of agreement between data and predictive samples as for the bursty model with *lacUV5* in Figure 3(B).

One interesting feature from Figure 3(C) is that burst rate varies far more widely, over a range of ~ 10^2^, than burst size, confined to a range of ≲ 10^1^ (and with the exception of promoter 6, just a span of 3 to 5-fold). This suggests that *k_i_*, not *b*, is the key dynamic variable that promoter sequence tunes.

#### 4.1.3 Connecting inferred parameters to prior work

It is interesting to connect these inferences on *k_i_* and *b* to the work of [54], where these same 18 promoters were considered through the lens of the three-state equilibrium model (model 2 in Figure 1(B)) and binding energies Δ*ε_P_* were predicted from an energy matrix model derived from [53]. As previously discussed the thermodynamic models of gene regulation can only make statements about the mean gene expression. This implies that we can draw the connection between both frameworks by equating the mean mRNA 〈*m*〉. This results in

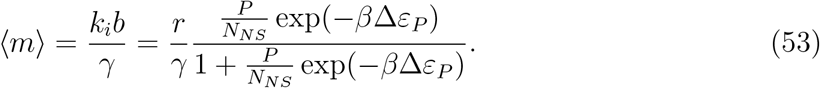

By taking the weak promoter approximation for the equilibrium model (*P/N_NS_* exp(−*β*Δ*ε_r_*) ≪ 1) results in

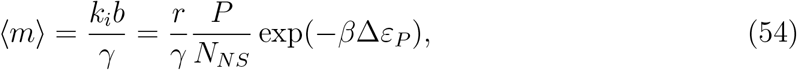

valid for all the binding energies considered here.

Given this result, how are the two coarse-grainings related? A quick estimate can shed some light. Consider for instance the *lacUV5* promoter, which we see from Figure 3(A) has *k_i_/γ* ~ *b* ~ few, from Figure 3(B) has 〈*m*〉 ~ 20, and from [54] has *β*Δ*ε_P_* ~ −6.5. Further we generally assume *P/N_NS_* ~ 10^−3^ since *N_NS_* ≈ 4.6 × 10^6^ and *P* ~ 10^3^. After some guess-and-check with these values, one finds the only association that makes dimensional sense and produces the correct order-of-magnitude for the known parameters is to take

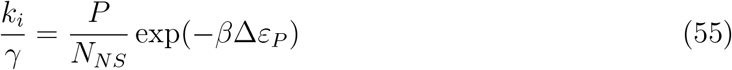

and

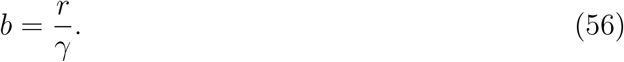

Figure 3(D) shows that this linear scaling between ln *k_i_* and −*β*Δ*ε_P_* is approximately true for all 18 constitutive promoters considered. The plotted line is simply Eq. 55 and assumes *P* ≈ 5000.

While the associations represented by Eq. 55 and Eq. 56 appear to be borne out by the data in Figure 3, we do not find the association of parameters they imply to be intuitive. We are also cautious to ascribe too much physical reality to the parameters. Indeed, part of our point in comparing the various constitutive promoter models is to demonstrate that these models each provide an internally self-consistent framework that adequately describes the data, but attempting to translate between models reveals the dubious physical interpretation of their parameters.

We mention one further comparison, between our inferred parameters and the work of Chong et. al. [47], which is interesting and puzzling. Beautiful experiments in [47] convincingly argue that supercoiling accumulated from the production of mRNA transcripts is key in setting the burstiness of mRNA production. In their model, this supercoiling occurs on the scale of 100 kb domains of DNA. This suggests that all genes on a given domain should burst in synchrony, and that the difference between highly and lowly expressed genes is the *size* of transcriptional bursts, not the *time between* bursts. But here, all burst sizes we infer in Figure 3(C) are comparable and burst rates vary wildly. It is not immediately clear how to square this circle. Furthermore, Figure 7E in [47] reports values of the quantity they label *β/α* and we label *k*^+^/*k^−^* in model 4 from Figure 2. In contrast to the findings of [42], Chong et. al. do not find *k*^+^/*k^−^* ≪ 1 for most of the genes they consider. This begs the question: is the *galK* chromosomal locus used for the reporter constructs in [42] and [36] merely an outlier, or is there a deeper puzzle here waiting to be resolved? Without more apples-to-apples data we can only speculate, and we leave it as an intriguing open question for the field.

Despite such puzzles, our goal here is not to unravel the mysterious origins of burstiness in transcription. Our remaining task in this work is a determination of the physical reality of equilibrium binding energies in Figure 1, as codified by the equilibrium-nonequilibrium equivalence of Eq. 35. For our phenomenological needs here model 5 in Figure 2 is more than adequate: the posterior distributions in Figure 3(C) are cleanly identifiable and the predictive checks in Figure S5 indicate no discrepancies between the model and the mRNA single-moleucle count data of [36]. Of the models we have considered it is unique in satisfying both these requirements. So we will happily use it as a foundation to build upon in the next section when we add regulation.

### 4.2 Transcription factor kinetics can be inferred from single-cell mRNA distribution measurements

#### 4.2.1 Building the model and performing parameter inference

Now that we have a satisfactory model in hand for constitutive promoters, we would like to return to the main thread: can we reconcile the equilibrium and nonequilibrium models by putting to the test Eq. 35, the correspondence between indirectly inferred equilibrium binding energies and nonequilibrium kinetic rates? To make this comparison, is it possible to infer repressor binding and unbinding rates from mRNA distributions over a population of cells as measured by single-molecule Fluorescence *in situ* Hybridization in [36]? If so, how do these inferred rates compare to direct single-molecule measurements such as from [56] and to binding energies such as from [33] and [32], which were inferred under the assumptions of the equilibrium models in Figure 1(B)? And can this comparison shed light on the unreasonable effectiveness of the equilibrium models, for instance, in their application in [35], [63]?

As we found in Section 3, for our purposes the “right” model of a constitutive promoter is the bursty picture, model five in Figure 2(A). Therefore our starting point here is the analogous model with repressor added, model 5 in Figure 1(C). For a given repressor binding site and copy number, this model has four rate parameters to be inferred: the repressor binding and unbinding rates 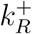, and 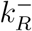, the initiation rate of bursts, *k_i_*, and the mean burst size *b* (we nondimensionalize all of these by the mRNA degradation rate *γ*).

Before showing the mathematical formulation of our statistical inference model, we would like to sketch the intuitive structure. The dataset from [36] we consider consists of single-cell mRNA counts data of nine different conditions, spanning several combinations of three unique repressor binding sites and four unique repressor copy numbers. We assume that the values of *k_i_* and *b* are known, since we have already cleanly inferred them from constitutive promoter data, and further we assume that these values are the same across datasets with different repressor binding sites and copy numbers. In other words, we assume that the regulation of the transcription factor does not affect the mean burst size nor the burst initiation rate. The regulation occurs as the promoter is taken away from the transcriptionally active state when the promoter is bound by repressor. We assume that there is one unbinding rate parameter for each repressor binding site, and likewise one binding rate for each unique repressor copy number. This makes our model seven dimensional, or nine if one counts *k_i_* and *b* as well. Note that we use only a subset of the datasets from Jones et. al. [36], as discussed more in Appendix S3.

Formally now, denote the set of seven repressor rates to be inferred as

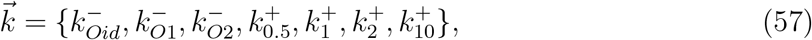

where subscripts for dissociation rates *k^−^* indicate the different repressor binding sites, and subscripts for association rates *k*^+^ indicate the concentration of the small-molecule that controlled the expression of the LacI repressor (see Appendix S3). This is because for this particular dataset the repressor copy numbers were not measured directly, but it is safe to assume that a given concentration of the inducer resulted in a specific mean repressor copy number [63]. Also note that the authors of [36] report estimates of LacI copy number per cell rather than direct measurements. However, these estimates were made assuming the validity of the equilibrium models in Figure 1, and since testing these models is our present goal, it would be circular logic if we were to make the same assumption. Therefore we will make no assumptions about the LacI copy number for a given inducer concentrations.

Having stated the problem, Bayes’ theorem reads

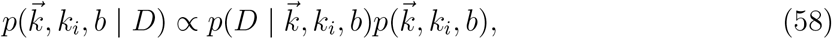

where *D* is again the set of all *N* observed single-cell mRNA counts across the various conditions. We assume that individual single-cell measurements are independent so that the likelihood factorizes as

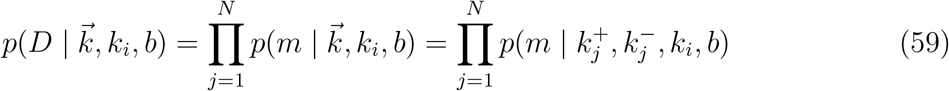

where 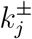 represent the appropriate binding and unbinding rates out of 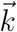 for the *j*-th measured cell. The probability 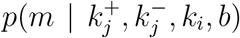 appearing in the last expression is exactly Eq. S184, the steady-state distribution for our bursty model with repression derived in Section S2, which for completeness we reproduce here as

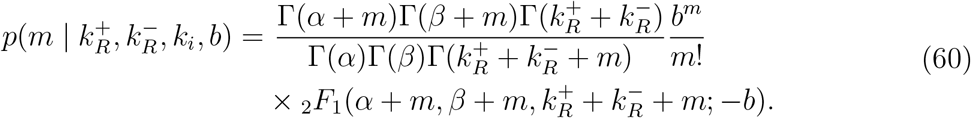

where _2_*F*_1_ is the confluent hypergeometric function of the second kind and *α* and *β*, defined for notational convenience, are

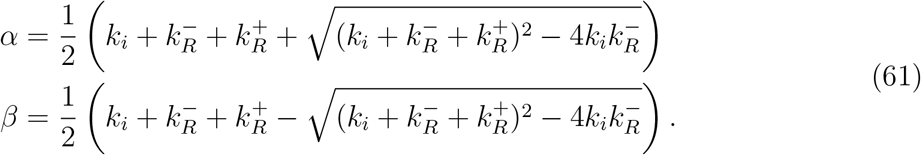

This likelihood is rather inscrutable. We did not find any of the known analytical approximations for _2_*F*_1_ terribly useful in gaining intuition, so we instead resorted to numerics. One insight we found was that for very strong or very weak repression, the distribution in Eq. 60 is well approximated by a negative binomial with burst size *b* and burst rate *k_i_* equal to their constitutive *lacUV5* values, except with *k*_i_ multiplied by the fold-change 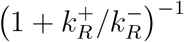. In other words, once again only the ratio 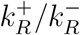 detectable. But for intermediate repression, the distribution was visibly broadened with Fano factor greater than 1 + *b*, the value for the corresponding constitutive case. This indicates that the repressor rates had left an imprint on the distribution, and perhaps intuitively, this intermediate regime occurs for values of 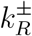 comparable to the burst rate *k_i_*. Put another way, if the repressor rates are much faster or much slower than *k_i_*, then there is a timescale separation and effectively only one timescale remains, 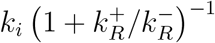. Only when all three rates in the problem are comparable does the mRNA distribution retain detectable information about them.

Next we specify priors. As for the constitutive model, weakly informative log-normal priors are a natural choice for all our rates. We found that if the priors were too weak, our MCMC sampler would often become stuck in regions of parameter space with very low probability density, unable to move. We struck a balance in choosing our prior widths between helping the sampler run while simultaneously verifying that the marginal posteriors for each parameter were not artificially constrained or distorted by the presence of the prior. All details for our prior distributions are listed in Appendix S3.

We ran MCMC sampling on the full nine dimensional posterior specified by this model. To attempt to visualize this object, in Figure 4(A) we plot several two-dimensional slices as contour plots, analogous to Figure 3(C). Each of these nine slices corresponds to the 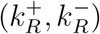 pair of rates for one of the conditions from the dataset used to fit the model and gives a sense of the uncertainty and correlations in the posterior. We note that the 95% uncertainties of all the rates span about ~ 0.3 log units, or about a factor of two, with the exception of 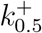, the association rate for the lowest repressor copy number which is somewhat larger.

#### 4.2.2 Comparison with prior measurements of repressor binding energies

Our primary goal in this work is to reconcile the kinetic and equilibrium pictures of simple repression. Towards this end we would like to compare the repressor kinetic rates we have inferred with the repressor binding energies inferred through multiple methods in [33] and [32]. If the agreement is close, then it suggests that the equilibrium models are not wrong and the repressor binding energies they contain correspond to physically real free energies, not mere fit parameters.

Figure 4(B) shows both comparisons, with the top panel comparing to equilibrium binding energies and the bottom panel comparing to single-molecule measurements. First consider the top panel and its comparison between repressor kinetic rates and binding energies. As described in section 2, if the equilibrium binding energies from [33] and [32] indeed are the physically real binding energies we believe them to be, then they should be related to the repressor kinetic rates via Eq. 35, which we restate here,

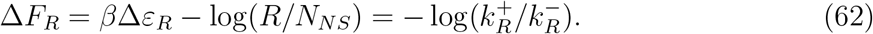

Assuming mass action kinetics implies that 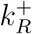 is proportional to repressor copy number *R*, or more precisely, it can be thought of as repressor copy number times some intrinsic per molecule association rate. But since *R* is not directly known for our data from [36], we cannot use this equation directly. Instead we can consider two different repressor binding sites and compute the *difference* in binding energy between them, since this difference depends only on the unbinding rates and not on the binding rates. This can be seen by evaluating Eq. 62 for two different repressor binding sites, labeled (1) and (2), but with the same repressor copy number *R*, and taking the difference to find

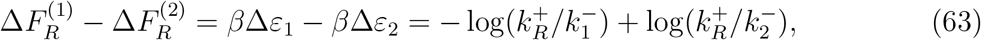

or simply

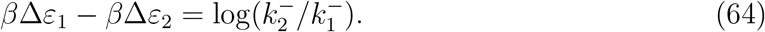

The left and right hand sides of this equation are exactly the horizontal and vertical axes of the top panel of Figure 4. Since we inferred rates for three repressor binding sites (O1, O2, and Oid), there are only two independent differences that can be constructed, and we arbitrarily chose to plot O2-O1 and O1-Oid in Figure 4(B). Numerically, we compute values of 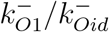 and 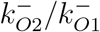 directly from our full posterior samples, which conveniently provides uncertainties as well, as detailed in Appendix S3. We then compare these log ratios of rates to the binding energy differences Δ*ε_O_*_1_ − Δ*ε_Oid_* and from Δ*ε_O_*_2_ −Δ*ε_O_*_1_ as computed from the values from both [33] and [32]. Three of the four values are within ~ 0.5 *k_B_T* of the diagonal representing perfect agreement, which is comparable to the ~ few × 0.1 *k_B_T* variability between the independent determinations of the same quantities between [33] and [32]. The only outlier involves Oid measurements from [32], and as the authors of [32] note, this is a difficult measurement of low fluorescence signal against high background since Oid represses so strongly. We are therefore inclined to regard the failure of this point to fall near the diagonal as a testament to the difficulty of the measurement and not as a failure of our theory.

On the whole then, we regard this as striking confirmation of the validity of the equilibrium models. Their lynchpin parameter is a phenomenological free energy of repressor binding that has previously only been inferred indirectly. Our result shows that the microscopic interpretation of this free energy, as the log of a ratio of transition rates, does indeed hold true to within the inherent uncertainties that remain in the entire theory-experiment dialogue.

#### 4.2.3 Comparison with prior measurements of repressor kinetics

In the previous section we established the equivalence between the equilibrium models’ binding energies and the repressor kinetics we infer from mRNA population distributions. But one might worry that the repressor rates we infer from mRNA distributions are *themselves* merely fitting parameters and that they do not actually correspond to the binding and unbinding rates of the repressor in vivo. To verify that this is not the case, we next compare our kinetic rates with a different measurement of the same rates using a radically different method: single molecule measurements as performed in Hammar et. al. [56]. This is plotted in the lower panel of Figure 4(B).

Since we do not have access to repressor copy number for either the single-cell mRNA data from [36] or the single-molecule data from [56], we cannot make an apples-to-apples comparison of association rates 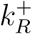. Further, while Hammar et. al. directly measure the dissociation rates 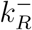, our inference procedure returns 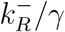, i.e., the repressor dissociation rate nondimensionalized by the mRNA degradation rate *γ*. So to make the comparison, we must make an assumption for the value of *γ* since it was not directly measured. For most mRNAs in *E. coli*, quoted values for the typical mRNA lifetime *γ^−^*^1^ range between about 2.5 min [64] to 8 min. We chose *γ^−^*^1^ = 3 min and *γ^−^*^1^ = 5 min as representative values and plot a comparison of 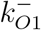 and 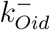 from our inference with corresponding values reported in [56] for both these choices of *γ*.

The degree of quantitative agreement in the lower panel of Figure 4(B) clearly depends on the precise choice of *γ*. Nevertheless we find this comparison very satisfying, when two wildly different approaches to a measurement of the same quantity yield broadly compatible results. We emphasize the agreement between our rates and the rates reported in [56] for any reasonable *γ*: values differ by at most a factor of 2 and possibly agree to within our uncertainties of 10-20%. From this we feel confident asserting that the parameters we have inferred from Jones et. al.’s single-cell mRNA counts data do in fact correspond to repressor binding and unbinding rates, and therefore our conclusions on the agreement of these rates with binding energies from [33] and [32] are valid.

#### 4.2.4 Model checking

In Figure 3(B) we saw that the simple Poisson model of a constitutive promoter, despite having a well behaved posterior, was clearly insufficient to describe the data. It behooves us to carry out a similar check for our model of simple repression, codified by Eq. 60 for the steady-state mRNA copy number distribution. As derived in Sections 2 and 3, we have compelling theoretical reasons to believe it is a good model, but if it nevertheless turned out to be badly contradicted by the data we should like to know.

The details are deferred to Appendix S3, and here we only attempt to summarize the intuitive ideas, as detailed at greater length by Jaynes [65] as well as Gelman and coauthors [52], [66]. From our samples of the posterior distribution, plotted in Figure 4(A), we generate many replicate data using a random number generator. In Figure 4(C), we plot empirical cumulative distribution functions of the middle 95% quantiles of these replicate data with the actual experimental data from Jones et. al. [36] overlaid, covering all ten experimental conditions spanning repressor binding sites and copy numbers (as well as the constitutive baseline UV5).

The purpose of Figure 4(C) is simply a graphical, qualitative assessment of the model: do the experimental data systematically disagree with the simulated data, which would suggest that our model is missing important features? A further question is not just whether there is a detectable difference between simulated and experimental data, but whether this difference is likely to materially affect the conclusions we draw from the posterior in Figure 4(A). More rigorous and quantitative statistical tests are possible [52], but their quantitativeness does not necessarily make them more useful. As stated in [66], we often find this graphical comparison more enlightening because it better engages our intuition for the model, not merely telling *if* the model is wrong but suggesting *how* the model may be incomplete.

Our broad brush takeaway from Figure 4(C) is overall of good agreement. There some oddities, in particular the long tails in the data for Oid, 1 ng/mL, and O2, 0.5 ng/mL. The latter is especially odd since it extends beyond the tail of the unregulated UV5 distribution. This is a relatively small number of cells, however, so whether this is a peculiarity of the experimental data, a statistical fluke of small numbers, or a real biological effect is unclear. It is conceivable that there is some very slow timescale switching dynamics that could cause this bimodality, although it is unclear why it would only appear for specific repressor copy numbers. There is also a small offset between experiment and simulation for O2 at the higher repressor copy numbers, especially at 2 and 10 ng/mL. From the estimate of repressor copy numbers from [36], it is possible that the repressor copy numbers here are becoming large enough to partially invalidate our assumption of a separation of timescales between burst duration and repressor association rate. Another possibility is that the very large number of zero mRNA counts for Oid, 2 ng/mL is skewing its partner datasets through the shared association rate. None of these fairly minor quibbles cause us to seriously doubt the overall correctness of our model, which further validates its use to compare the equilibrium models’ binding energies to the nonequilibrium models’ repressor kinetics, as we originally set out to do.

## 5 Discussion and future work

The study of gene expression is one of the dominant themes of modern biology, made all the more urgent by the dizzying pace at which genomes are being sequenced. But there is a troubling Achilles heel buried in all of that genomic data, which is our inability to find and interpret regulatory sequence. In many cases, this is not possible even qualitatively, let alone the possibility of quantitative dissection of the regulatory parts of genomes in a predictive fashion. Other recent work has tackled the challenge of finding and annotating the regulatory part of genomes [5], [28]. Once we have determined the architecture of the regulatory part of the genome, we are then faced with the next class of questions which are sharpened by formulating them in mathematical terms, namely, what are the input-output properties of these regulatory circuits and what knobs control them?

The present work has tackled that question in the context of the first regulatory architecture hypothesized in the molecular biology era, namely, the repressor-operator model of Jacob and Monod [1]. Regulation in that architecture is the result of a competition between a repressor which inhibits transcription and RNAP polymerase which undertakes it. Through the labors of generations of geneticists, molecular biologists and biochemists, an overwhelming amount of information and insight has been garnered into this simple regulatory motif, licensing it as what one might call the “hydrogen atom” of regulatory biology. It is from that perspective that the present paper explores the extent to which some of the different models that have been articulated to describe that motif allow us to understand both the average level of gene expression found in a population of cells, the intrinsic cell-to-cell variability, and the full gene expression distribution found in such a population as would be reported in a single molecule mRNA Fluorescence *in situ* Hybridization experiment, for example.

Our key insights can be summarized as follows. First, as shown in Figure 1, the mean expression in the simple repression architecture is captured by a master curve in which the action of repressor and the details of the RNAP interaction with the promoter appear separately and additively in an effective free energy. Interestingly, as has been shown elsewhere in the context of the Monod-Wyman-Changeux model, these kinds of coarse-graining results are an exact mathematical result and do not constitute hopeful approximations or biological naivete [32], [35]. To further dissect the relative merits of the different models, we must appeal to higher moments of the gene expression probability distribution. To that end, our second set of insights focus on gene expression noise, where it is seen that a treatment of the constitutive promoter already reveals that some models have Fano factors (variance/mean) that are less than one, at odds with any and all experimental data that we are aware of [36], [59]. This theoretical result allows us to directly discard a subset of the models (models 1-3 in Figure 2(A)) since they cannot be reconciled with experimental observations. The two remaining models (models 4 and 5 in Figure 2) appear to contain enough microscopic realism to be able to reproduce the data. A previous exploration of model 4 demonstrated the “sloppy” [62] nature of the model in which data on single-cell mRNA counts alone cannot constrain the value of all parameters simultaneously [42]. Here we demonstrate that the proposed one-state bursty promoter model (model 5 in Figure 2(A)) emerges as a limit of the commonly used two-state promoter model [19], [23], [36], [57], [59]. We put the idea to the test that this level of coarse-graining is rich enough to reproduce previous experimental observations. In particular we perform Bayesian inference to determine the two parameters describing the full steady-state mRNA distribution, finding that the model is able to provide a quantitative description of a plethora of promoter sequences with different mean levels of expression and noise.

With the results of the constitutive promoter in hand, we then fix the parameters associated with this class of promoters and use them as input for evaluating the noise in gene expression for the simple repression motif itself. This allows us to provide a single overarching analysis of both the constitutive and simple repression architectures using one simple model and corresponding set of self-consistent parameters, demonstrating not only a predictive framework, but also reconciling the equilibrium and non-equilibrium views of the same simple repression constructs. More specifically, we obtained values for the transcription factor association and dissociation rates by performing Bayesian inference on the full mRNA distribution for data obtained from simple-repression promoters with varying number of transcription factors per cell and affinity of such transcription factors for the binding site. The free energy value obtained from these kinetic rates – computed as the log ratio of the rates – agrees with previous inferences performed only from mean gene expression measurements, that assumed an equilibrium rather than a kinetic framework [32], [33].

It is interesting to speculate what microscopic details are being coarse-grained by our burst rate and burst size in Figure 2, model 5. Chromosomal locus is one possible influence we have not addressed in this work, as all the single-molecule mRNA data from [36] that we considered was from a construct integrated at the *galK* locus. The results of [47] indicate that transcription-induced supercoiling contributes substantially in driving transcriptional bursting, and furthermore, their Figure 7 suggests that the parameters describing the rate, duration, and size of bursts vary substantially for transcription from different genomic loci. Although the authors of [67] do not address noise, they note enormous differences in mean expression levels when an identical construct is integrated at different genomic loci. The authors of [68] attribute noise and burstiness in their single-molecule mRNA data to the influence of different sigma factors, which is a reasonable conclusion from their data. Could the difference also be due to the different chromosomal locations of the two operons? What features of different loci are and are not important? Could our preferred coarse-grained model capture the variability across different loci? If so, and we were to repeat the parameter inference as done in this work, is there a simple theoretical model we could build to understand the resulting parameters?

In summary, this work took up the challenge of exploring the extent to which a single specific mechanistic model of the simple-repression regulatory architecture suffices to explain the broad sweep of experimental data for this system. Pioneering early experimental efforts from the Müller-Hill lab established the simple-repression motif as an arena for the quantitative dissection of regulatory response in bacteria, with similar beautiful work emerging in examples such as the *ara* and *gal* operons as well [29], [30], [69]–[73]. In light of a new generation of precision measurements on these systems, the definition of what it means to understand them can now be formulated as a rigorous quantitative question. In particular, we believe understanding of the simple repression motif has advanced sufficiently that the design of new versions of the architecture is now possible, based upon predictions about how repressor copy number and DNA binding site strength control expression. In our view, the next step in the progression is to first perform similar rigorous analyses of the fundamental “basis set” of regulatory architectures. Natural follow-ups to this work are explorations of motifs such as simple activation that is regulated by a single activator binding site, and the repressor-activator architecture, mediated by the binding of both a single activator and a single repressor, and beyond. With the individual input-output functions in hand, similar quantitative dissections including the rigorous analysis of their tuning parameters can be undertaken for the “basis set” of full gene-regulatory networks such as switches, feed-forward architectures and oscillators for example, building upon the recent impressive bonanza of efforts from systems biologists and synthetic biologists [74], [75].

## 6 Methods

All data and custom scripts were collected and stored using Git version control. Code for Bayesian inference and figure generation is available on the GitHub repository (https://github.com/RPGroup-PBoC/bursty_transcription).

## Acknowledgments

We thank Rob Brewster for providing the raw single-molecule mRNA FISH data. We thank Justin Bois for his key support with the Bayesian inference section. We would also like to thank Griffin Chure for invaluable feedback on the manuscript. This material is based upon work supported by the National Science Foundation Graduate Research Fellowship under Grant No. DGE-1745301. This work was also supported by La Fondation Pierre-Gilles de Gennes, the Rosen Center at Caltech, and the NIH 1R35 GM118043 (MIRA). M.R.M. was supported by the Caldwell CEMI fellowship.

