## Supplemental Information for "Reconciling Kinetic and Equilibrium Models of Bacterial Transcription"

### S1 Derivations for non-bursty promoter models

In this section we detail the calculation of mean mRNA levels, fold-changes in expression, and Fano factors for nonequilibrium promoter models 1 through 4 in Figure 1. These are the results that were quoted but not derived in Sections 2 and 3 of the main text. In each of these four models, the natural mathematicization of their cartoons is as a chemical master equation such as Eq. 12 for model 1. Before jumping into the derivations of the general computation of the mean mRNA level and the Fano factor we will work through the derivation of an example master equation. In particular we will focus on model 1 from Figure 1(C). The general steps are applicable to all other chemical master equations in this work.

#### S1.1 Derivation of chemical master equation

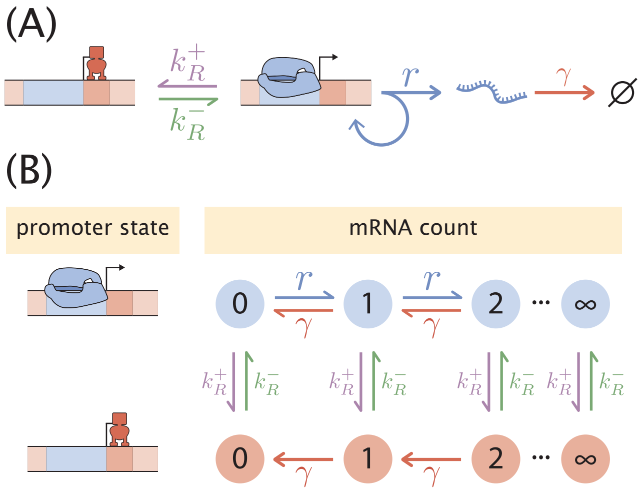

**Figure S1. Two-state promoter chemical master equation.** (A) Schematic of the two state promoter simple repression model. Rates $k_R^+$ and $k_R^-$ are the association and dissociation rates of the transcriptional repressor, respectively, $r$ is the transcription initiation rate, and $\gamma$ is the mRNA degradation rate. (B) Schematic depiction of the mRNA count state transitions. The model in (A) only allows for jumps in mRNA of size 1. The production of mRNA can only occur when the promoter is in the transcriptionally active state.

The chemical master equation describes the continuous time evolution of a continuous or discrete probability distribution function. In our specific case we want to describe the time evolution of the discrete mRNA distribution. What this means is that we want to compute the probability of having $m$ mRNA molecules at time $t + \Delta t$, where $\Delta t$ is a sufficiently small time interval such that only one of the possible reactions take place during that time interval.

For the example that we will work out here in detail we chose the two-state stochastic simple repression model schematized in Figure S1(A). To derive the master equation we will focus more on the representation shown in Figure S1(B), where the transitions between different mRNA counts and promoter states is more explicitly depicted. Given that the DNA promoter can exist in one of two states – transcriptionally active state, and with repressor bound – we will keep track not only of the mRNA count, but on which state the promoter is. For this we will keep track of two probability distributions: The probability of having $m$ mRNAs at time $t$ when the promoter is in the transcriptionally active state $A$, $p_A(m, t)$, and the equivalent probability but when the promoter is in the repressor bound state $R$, $p_R(m, t)$.

Since mRNA production can only take place in the transcriptionally active state we will focus on this function for our derivation. The repressor bound state will have an equivalent equation without terms involving the production of mRNAs. We begin by listing the possible state transitions that can occur for a particular mRNA count with the promoter in the active state. For state changes in a small time window $\Delta t$ that "jump into" state $m$ in the transcriptionally active state we have

- Produce an mRNA, jumping from $m - 1$ to $m$.
- Degrade an mRNA, jumping from $m + 1$ to $m$.
- Transition from the repressor bound state $R$ with $m$ mRNAs to the active state $A$ with $m$ mRNAs.

Likewise, for state transitions that "jump out" of state $m$ in the transcriptionally inactive state we have

- Produce an mRNA, jumping from $m$ to $m + 1$.
- Degrade an mRNA, jumping from $m$ to $m - 1$.
- Transition from the active state $A$ with $m$ mRNAs to the repressor bound state $R$ with $m$ mRNAs.

The mRNA production does not depend on the current number of mRNAs, therefore these state transitions occur with probability $r\Delta t$. The same is true for the promoter state transitions; each occurs with probability $k_R^{\pm}\Delta t$. As for the mRNA degradation events, these transitions depend on the current number of mRNA molecules since the more molecules of mRNA there are, the more will decay during a given time interval. Each molecule has a constant probability $\gamma\Delta t$ of being degraded, so the total probability for an mRNA degradation event to occur is computed by multiplying this probability by the current number of mRNAs.

To see these terms in action let us compute the probability of having $m$ mRNA at time

$t + \Delta t$ in the transcriptionally active state. This takes the form

$$
\begin{aligned}
p_A(m, t+\Delta t) = p_A(m,t) \\
&+ \overbrace{(r\Delta t) p_A(m-1,t)}^{m-1\to m} - \overbrace{(r\Delta t) p_A(m,t)}^{m\to m+1} \\
&+ \overbrace{(m+1)(\gamma\Delta t) p_A(m+1,t)}^{m+1\to m} - \overbrace{m(\gamma\Delta t) p_A(m,t)}^{m\to m-1} \\
&+ \overbrace{(k_R^-\Delta t) p_R(m,t)}^{R\to A} - \overbrace{(k_R^+\Delta t) p_A(m,t)}^{A\to R},
\end{aligned}
\tag{S1}
$$

where the overbrace indicates the corresponding state transitions. Notice that the second to last term on the right-hand side is multiplied by $p_R(m,t)$ since the transition from state $R$ to state $A$ depends on the probability of being in state $R$ to begin with. It is through this term that the dynamics of the two probability distribution functions ($p_R(m,t)$ and $p_A(m,t)$) are coupled. An equivalent equation can be written for the probability of having $m$ mRNA at time $t + \Delta t$ while in the repressor bound state, the only difference being that the mRNA production rates are removed, and the sign for the promoter state transitions are inverted. This is

$$
\begin{aligned}
p_R(m, t+\Delta t) = p_R(m,t) \\
&+ \overbrace{(m+1)(\gamma\Delta t) p_R(m+1,t)}^{m+1\to m} - \overbrace{m(\gamma\Delta t) p_R(m,t)}^{m\to m-1} \\
&- \overbrace{(k_R^-\Delta t) p_R(m,t)}^{R\to A} + \overbrace{(k_R^+\Delta t) p_A(m,t)}^{A\to R}.
\end{aligned}
\tag{S2}
$$

All we have to do now are simple algebraic steps in order to simplify the equations. Let us focus on the transcriptionally active state $A$. First we will send the term $p_A(m,t)$ to the right-hand side, and then we will divide both sides of the equation by $\Delta t$. This results in

$$
\begin{aligned}
\frac{p_A(m,t+\Delta t) - p_A(m,t)}{\Delta t} &= r p_A(m-1,t) - r p_A(m,t) \\
&+ (m+1)\gamma p_A(m+1,t) - m\gamma p_A(m,t) \\
&+ k_R^- p_R(m,t) - k_R^+ p_A(m,t).
\end{aligned}
\tag{S3}
$$

Upon taking the limit when $\Delta t \to 0$ we can transform the left-hand side into a derivative, obtaining the chemical master equation

$$
\begin{aligned}
\frac{dp_A(m,t)}{dt} &= r p_A(m-1,t) - r p_A(m,t) \\
&+ (m+1)\gamma p_A(m+1,t) - m\gamma p_A(m,t) \\
&+ k_R^- p_R(m,t) - k_R^+ p_A(m,t).
\end{aligned}
\tag{S4}
$$

Doing equivalent manipulations for the repressor bound state gives an ODE of the form

$$
\begin{aligned}
\frac{dp_R(m,t)}{dt} &= (m+1)\gamma p_R(m+1,t) - m\gamma p_R(m,t) \\
&- k_R^- p_R(m,t) + k_R^+ p_A(m,t).
\end{aligned}
\tag{S5}
$$

In the next section we will write these coupled ODEs in a more compact form using matrix notation.

#### S1.2 Matrix form of the multi-state chemical master equation

Having derived an example chemical master equation we now focus on writing a general matrix form for the kinetic models 1-4 shown in Figure 1(C) in the main text. In each of these four models, the natural mathematicization of their cartoons is as a chemical master equation such as Eq. 12 for model 1, which for completeness we reproduce here as

$$\begin{aligned}
\frac{d}{dt}p_R(m,t) =& -\overbrace{k_R^- p_R(m,t)}^{R\to U} + \overbrace{k_R^+ p_U(m,t)}^{U\to R} + \overbrace{(m+1)\gamma p_R(m+1,t)}^{m+1\to m} - \overbrace{\gamma m p_R(m,t)}^{m\to m-1} \\
\frac{d}{dt}p_U(m,t) =& \overbrace{k_R^- p_R(m,t)}^{R\to U} - \overbrace{k_R^+ p_U(m,t)}^{U\to R} + \overbrace{r p_U(m-1,t)}^{m-1\to m} - \overbrace{r p_U(m,t)}^{m\to m+1} \\
&+ \overbrace{(m+1)\gamma p_U(m+1,t)}^{m+1\to m} - \overbrace{\gamma m p_U(m,t)}^{m\to m-1}.
\end{aligned} \tag{S6}$$

Here $p_R(m,t)$ and $p_U(m,t)$ are the probabilities of finding the system with $m$ mRNA molecules at time $t$ either in the repressor bound or unbound states, respectively. $r$ is the probability per unit time that a transcript will be initiated when the repressor is unbound, and $\gamma$ is the probability per unit time for a given mRNA to be degraded. $k_R^-$ is the probability per unit time that a bound repressor will unbind, while $k_R^+$ is the probability per unit time that an unbound operator will become bound by a repressor. Assuming mass action kinetics, $k_R^+$ is proportional to repressor copy number $R$.

Next consider the cartoon for nonequilibrium model 2 in Figure 1(C). Now we must track probabilities $p_R$, $p_P$, and $p_E$ for the repressor bound, empty, and polymerase bound states, respectively. By analogy to Eq. S6, the master equation for model 2 can be written

$$\begin{aligned}
\frac{d}{dt}p_R(m,t) =& -\overbrace{k_R^- p_R(m,t)}^{R\to U} + \overbrace{k_R^+ p_E(m,t)}^{U\to R} + \overbrace{(m+1)\gamma p_R(m+1,t)}^{m+1\to m} - \overbrace{\gamma m p_R(m,t)}^{m\to m-1} \\
\frac{d}{dt}p_E(m,t) =& \overbrace{k_R^- p_R(m,t)}^{R\to U} - \overbrace{k_R^+ p_E(m,t)}^{U\to R} + \overbrace{(m+1)\gamma p_E(m+1,t)}^{m+1\to m} - \overbrace{\gamma m p_E(m,t)}^{m\to m-1}. \\
&+ \overbrace{k_P^- p_P(m,t)}^{A\to U} - \overbrace{k_P^+ p_E(m,t)}^{U\to A} + \overbrace{r p_P(m-1,t)}^{m-1\to m,\ A\to U} \\
\frac{d}{dt}p_P(m,t) =& -\overbrace{k_P^- p_P(m,t)}^{A\to U} + \overbrace{k_P^+ p_E(m,t)}^{U\to A} + \overbrace{(m+1)\gamma p_P(m+1,t)}^{m+1\to m} - \overbrace{\gamma m p_P(m,t)}^{m\to m-1}. \\
&- \overbrace{r p_P(m,t)}^{m\to m+1,\ A\to U}.
\end{aligned} \tag{S7}$$

$k_P^+$ and $k_P^-$ are defined in close analogy to $k_R^+$ and $k_R^-$, except for RNAP binding and unbinding instead of repressor. Similarly $p_P(m,t)$ is defined for the active (RNAP-bound) state exactly as are $p_R(m,t)$ and $p_E(m,t)$ for the repressor bound and unbound states, respectively. The new subtlety Eq. S7 introduces compared to Eq. S6 is that when mRNAs are produced, the promoter state also changes. This is encoded by the terms involving $r$, the last term in each of the equations for $p_E$ and $p_P$. The former accounts for arrivals in the unbound state and the latter accounts for departures from the RNAP-bound state.

To condense and clarify the unwieldy notation of Eq. S7, it can be rewritten in matrix form. We define the column vector $\vec{p}(m,t)$ as

$$\vec{p}(m,t) = \begin{pmatrix} p_R(m,t) \\ p_E(m,t) \\ p_P(m,t) \end{pmatrix} \tag{S8}$$

to gather, for a given $m$, the probabilities of finding the system in the three possible promoter states. Then all the transition rates may be condensed into matrices which multiply this vector. The first matrix is

$$\mathbf{K} = \begin{pmatrix} -k_R^- & k_R^+ & 0 \\ k_R^- & -k_R^+ - k_P^+ & k_P^- \\ 0 & k_P^+ & -k_P^- \end{pmatrix}, \tag{S9}$$

which tracks all promoter state changes in Eq. S7 that are *not* accompanied by a change in the mRNA copy number. The two terms accounting for transcription, the only transition that increases mRNA copy number, must be handled by two separate matrices given by

$$\mathbf{R_A} = \begin{pmatrix} 0 & 0 & 0 \\ 0 & 0 & r \\ 0 & 0 & 0 \end{pmatrix}, \ \mathbf{R_D} = \begin{pmatrix} 0 & 0 & 0 \\ 0 & 0 & 0 \\ 0 & 0 & r \end{pmatrix}. \tag{S10}$$

$\mathbf{R_A}$ accounts for transitions *arriving* in a given state while $\mathbf{R_D}$ tracks *departing* transitions. With these definitions, we can condense Eq. S7 into the single equation

$$\frac{d}{dt}\vec{p}(m,t) = (\mathbf{K} - \mathbf{R_D} - \gamma m \mathbf{I})\,\vec{p}(m,t) + \mathbf{R_A}\vec{p}(m-1,t) + \gamma(m+1)\mathbf{I}\vec{p}(m+1,t), \tag{S11}$$

which is just Eq. 15 in the main text. Straightforward albeit tedious algebra verifies that Eqs. S7 and S11 are in fact equivalent.

Although we derived Eq. S11 for the particular case of nonequilibrium model 2 in Figure 1, in fact the chemical master equations for all of the nonequilibrium models in Figure 1 except for model 5 can be cast in this form. (We treat model 5 separately in Appendix S2.) Model 3 introduces no new subtleties beyond model 2 and Eq. S11 applies equally well to it, simply with different matrices of dimension four instead of three. Models 1 and 4 are both handled by Eq. 12 in the main text, which is just Eq. S11 except in the special case of $\mathbf{R_D} = \mathbf{R_A} \equiv \mathbf{R}$, since in these two models transcription initiation events do not change promoter state.

Recalling that our goal in this section is to derive expressions for mean mRNA and Fano factor for nonequilibrium models 1 through four in Figure 1, and since all four of these models are described by Eq. S11, we can save substantial effort by deriving general formulas for mean mRNA and Fano factor from Eq. S11 once and for all. Then for each model we can simply plug in the appropriate matrices for $\mathbf{K}$, $\mathbf{R_D}$, and $\mathbf{R_A}$ and carry out the remaining algebra.

For our purposes it will suffice to derive the first and second moments of the mRNA distribution from this master equation, similar to the treatment in [24], but we refer the interested reader to [42] for an analogous treatment demonstrating an analytical solution for arbitrary moments.

#### S1.3 General forms for mean mRNA and Fano factor

Our task now is to derive expressions for the first two moments of the steady-state mRNA distribution from Eq. S11. Our treatment of this is analogous to that given in Refs. [24] and [42]. We first obtain the steady-state limit of Eq. S11 in which the time derivative vanishes, giving

$$0 = (\mathbf{K} - \mathbf{R_D} - \gamma m \mathbf{I})\, \vec{p}(m) + \mathbf{R_A} \vec{p}(m-1) + \gamma (m+1) \mathbf{I} \vec{p}(m+1), \tag{S12}$$

From this, we want to compute

$$\langle m \rangle = \sum_S \sum_{m=0}^{\infty} m\, p_S(m) \tag{S13}$$

and

$$\langle m^2 \rangle = \sum_S \sum_{m=0}^{\infty} m^2 p_S(m) \tag{S14}$$

which define the average values of $m$ and $m^2$ at steady state, where the averaging is over all possible mRNA copy numbers and promoter states $S$. For example, for model 1 in Figure 1(C), the sum on $S$ would cover repressor bound and unbound states ($R$ and $U$ respectively), for model 2, the sum would cover repressor bound, polymerase bound, and empty states ($R$, $P$, and $E$), and so on for the other models.

Along the way it will be convenient to define the following *conditional* moments as

$$\langle \vec{m} \rangle = \sum_{m=0}^{\infty} m \vec{p}(m), \tag{S15}$$

and

$$\langle \vec{m}^2 \rangle = \sum_{m=0}^{\infty} m^2 \vec{p}(m). \tag{S16}$$

These objects are vectors of the same size as $\vec{p}(m)$, and each component can be thought of as the expected value of the mRNA copy number, or copy number squared, conditional on the promoter being in a certain state. For example, for model 1 in Figure 1 which has repressor bound and unbound states labeled $R$ and $U$, $\langle \vec{m}^2 \rangle$ would be

$$\langle \vec{m}^2 \rangle = \begin{pmatrix} \sum_{m=0}^{\infty} m^2 p_R(m) \\ \sum_{m=0}^{\infty} m^2 p_U(m) \end{pmatrix}. \tag{S17}$$

Analogously to $\langle \vec{m} \rangle$ and $\langle \vec{m}^2 \rangle$, it is convenient to define the vector

$$\langle \vec{m}^0 \rangle = \sum_{m=0}^{\infty} \vec{p}(m), \tag{S18}$$

whose elements are simply the probabilities of finding the system in each of the possible promoter states. It will be convenient to denote by $\vec{1}^\dagger$ a row vector of the same length as $\vec{p}$ whose elements are all 1, such that, for instance, $\vec{1}^\dagger \cdot \langle \vec{m}^0 \rangle = 1$, $\vec{1}^\dagger \cdot \langle \vec{m} \rangle = \langle m \rangle$, etc.

##### S1.3.1 Promoter state probabilities $\langle \vec{m}^0 \rangle$

To begin, we will find the promoter state probabilities $\langle \vec{m}^0 \rangle$ from Eq. S12 by summing over all mRNA copy numbers $m$, producing

$$0 = \sum_{m=0}^{\infty} \left[ (\mathbf{K} - \mathbf{R_D} - \gamma m \mathbf{I}) \, \vec{p}(m) + \mathbf{R_A} \vec{p}(m-1) + \gamma(m+1)\mathbf{I}\vec{p}(m+1) \right] \tag{S19}$$

Using the definitions of $\langle \vec{m}^0 \rangle$ and $\langle \vec{m} \rangle$, and noting the matrices $\mathbf{K}$, $\mathbf{R_D}$, and $\mathbf{R_A}$ are all independent of $m$ and can be moved outside the sum, this simplifies to

$$0 = (\mathbf{K} - \mathbf{R_D})\langle \vec{m}^0 \rangle - \gamma \langle \vec{m} \rangle + \mathbf{R_A} \sum_{m=0}^{\infty} \vec{p}(m-1) + \gamma \sum_{m=0}^{\infty} (m+1)\vec{p}(m+1). \tag{S20}$$

The last two terms can be handled by reindexing the summations, transforming them to match the definitions of $\langle \vec{m}^0 \rangle$ and $\langle \vec{m} \rangle$. For the first, noting $\vec{p}(-1) = 0$ since negative numbers of mRNA are nonsensical, we have

$$\sum_{m=0}^{\infty} \vec{p}(m-1) = \sum_{m=-1}^{\infty} \vec{p}(m) = \sum_{m=0}^{\infty} \vec{p}(m) = \langle \vec{m}^0 \rangle. \tag{S21}$$

Similarly for the second,

$$\sum_{m=0}^{\infty} (m+1)\vec{p}(m+1) = \sum_{m=1}^{\infty} m\vec{p}(m) = \sum_{m=0}^{\infty} m\vec{p}(m) = \langle \vec{m} \rangle, \tag{S22}$$

which holds since in extending the lower limit from $m = 1$ to $m = 0$, the extra term we added to the sum is zero. Substituting these back in Eq. S20, we have

$$0 = (\mathbf{K} - \mathbf{R_D})\langle \vec{m}^0 \rangle - \gamma \langle \vec{m} \rangle + \mathbf{R_A} \langle \vec{m}^0 \rangle + \gamma \langle \vec{m} \rangle, \tag{S23}$$

or simply

$$0 = (\mathbf{K} - \mathbf{R_D} + \mathbf{R_A})\langle \vec{m}^0 \rangle. \tag{S24}$$

So given matrices $\mathbf{K}$, $\mathbf{R_D}$, and $\mathbf{R_A}$ describing a promoter, finding $\langle \vec{m}^0 \rangle$ simply amounts to solving this set of linear equations, subject to the normalization constraint $\vec{1}^\dagger \cdot \langle \vec{m}^0 \rangle = 1$. It will turn out to be the case that, if the matrix $\mathbf{K} - \mathbf{R_D} + \mathbf{R_A}$ is $n$ dimensional, it will always have only $n-1$ linearly independent equations. Including the normalization condition provides the $n$-th linearly independent equation, ensuring a unique solution. So when using this equation to solve for $\langle \vec{m}^0 \rangle$, we may always drop one row of the matrix equation at our convenience and supplement the system with the normalization condition instead. The reader may find it illuminating to skip ahead and see Eq. S24 in use with concrete examples, e.g., Eq. S52 and what follows it, before continuing on through the general formulas for moments.

##### S1.3.2 First moments $\langle \vec{m} \rangle$ and $\langle m \rangle$

By analogy to the above procedure for finding $\langle \vec{m}^0 \rangle$, we may find $\langle \vec{m} \rangle$ by first multiplying Eq. S12 by $m$ and then summing over $m$. Carrying out this procedure we have

$$0 = \sum_{m=0}^{\infty} m \left[ \left( \mathbf{K} - \mathbf{R_D} - \gamma m \mathbf{I} \right) \vec{p}(m) + \mathbf{R_A} \vec{p}(m-1) + \gamma(m+1) \mathbf{I} \vec{p}(m+1) \right], \tag{S25}$$

and now identifying $\langle \vec{m} \rangle$ and $\langle \vec{m}^2 \rangle$ gives

$$0 = (\mathbf{K} - \mathbf{R_D}) \langle \vec{m} \rangle - \gamma \langle \vec{m}^2 \rangle + \mathbf{R_A} \sum_{m=0}^{\infty} m \vec{p}(m-1) + \gamma \sum_{m=0}^{\infty} m(m+1) \vec{p}(m+1). \tag{S26}$$

The summations in the last two terms can be reindexed just as we did for $\langle \vec{m}^0 \rangle$, freely adding or removing terms from the sum which are zero. For the first term we find

$$\sum_{m=0}^{\infty} m \vec{p}(m-1) = \sum_{m=-1}^{\infty} (m+1) \vec{p}(m) = \sum_{m=0}^{\infty} (m+1) \vec{p}(m) = \langle \vec{m} \rangle + \langle \vec{m}^0 \rangle, \tag{S27}$$

and similarly for the second,

$$\sum_{m=0}^{\infty} m(m+1) \vec{p}(m+1) = \sum_{m=1}^{\infty} (m-1) m \vec{p}(m) = \sum_{m=0}^{\infty} (m-1) m \vec{p}(m) = \langle \vec{m}^2 \rangle - \langle \vec{m} \rangle. \tag{S28}$$

Substituting back in Eq. S26 then produces

$$0 = (\mathbf{K} - \mathbf{R_D}) \langle \vec{m} \rangle - \gamma \langle \vec{m}^2 \rangle + \mathbf{R_A} (\langle \vec{m} \rangle + \langle \vec{m}^0 \rangle) + \gamma (\langle \vec{m}^2 \rangle - \langle \vec{m} \rangle), \tag{S29}$$

or after simplifying

$$0 = (\mathbf{K} - \mathbf{R_D} + \mathbf{R_A} - \gamma) \langle \vec{m} \rangle + \mathbf{R_A} \langle \vec{m}^0 \rangle. \tag{S30}$$

So like $\langle \vec{m}^0 \rangle$, $\langle \vec{m} \rangle$ is also found by simply solving a set of linear equations after first solving for $\langle \vec{m}^0 \rangle$ from Eq. S24.

Next we can find the mean mRNA copy number $\langle m \rangle$ from $\langle \vec{m} \rangle$ according to

$$\langle m \rangle = \vec{1}^\dagger \cdot \langle \vec{m} \rangle, \tag{S31}$$

where $\vec{1}^\dagger$ is a row vector whose elements are all 1. Eq. S31 holds since the $i^{th}$ element of the column vector $\langle \vec{m} \rangle$ is the mean mRNA value conditional on the system occupying the $i^{th}$ promoter state, so the dot product with $\vec{1}^\dagger$ amounts to simply summing the elements of $\langle \vec{m} \rangle$.

Rather than solving Eq. S30 for $\langle \vec{m} \rangle$ and then computing $\vec{1}^\dagger \cdot \langle \vec{m} \rangle$, we may take a shortcut by multiplying Eq. S30 by $\vec{1}^\dagger$ first. The key observation that makes this useful is that

$$\vec{1}^\dagger \cdot (\mathbf{K} - \mathbf{R_D} + \mathbf{R_A}) = 0. \tag{S32}$$

Intuitively, this equality holds because each column of this matrix represents transitions to and from a given promoter state. In any given column, the diagonal encodes all departing transitions and off-diagonals encode all arriving transitions. Conservation of probability means that each column sums to zero, and summing columns is exactly the operation that multiplying by $\vec{1}^\dagger$ carries out.

Proceeding then in multiplying Eq. S30 by $\vec{1}^\dagger$ produces

$$0 = -\gamma \vec{1}^\dagger \cdot \langle \vec{m} \rangle + \vec{1}^\dagger \cdot \mathbf{R_A} \langle \vec{m}^0 \rangle, \tag{S33}$$

or simply

$$\langle m \rangle = \frac{1}{\gamma} \vec{1}^\dagger \cdot \mathbf{R_A} \langle \vec{m}^0 \rangle. \tag{S34}$$

We note that the in equilibrium models of transcription such as in Figure 1, it is usually *assumed* that the mean mRNA level is given by the ratio of initiation rate $r$ to degradation rate $\gamma$ multiplied by the probability of finding the system in the RNAP-bound state. Reassuringly, we have recovered exactly this result from the master equation picture: the product $\vec{1}^\dagger \cdot \mathbf{R_A} \langle \vec{m}^0 \rangle$ picks out the probability of the active promoter state from $\langle \vec{m}^0 \rangle$ and multiplies it by the initiation rate $r$.

##### S1.3.3 Second moment $\langle m^2 \rangle$ and Fano factor $\nu$

Continuing the pattern of the zeroth and first moments, we now find $\langle \vec{m}^2 \rangle$ by multiplying Eq. S12 by $m^2$ and then summing over $m$, which explicitly is

$$0 = \sum_{m=0}^{\infty} m^2 \left[ \left( \mathbf{K} - \mathbf{R_D} - \gamma m \mathbf{I} \right) \vec{p}(m) + \mathbf{R_A} \vec{p}(m-1) + \gamma (m+1) \mathbf{I} \vec{p}(m+1) \right]. \tag{S35}$$

Identifying the moments $\langle \vec{m}^2 \rangle$ and $\langle \vec{m}^3 \rangle$ in the first term simplifies this to

$$0 = (\mathbf{K} - \mathbf{R_D}) \langle \vec{m}^2 \rangle - \gamma \langle \vec{m}^3 \rangle + \mathbf{R_A} \sum_{m=0}^{\infty} m^2 \vec{p}(m-1) + \gamma \sum_{m=0}^{\infty} m^2 (m+1) \vec{p}(m+1). \tag{S36}$$

Reindexing the sums of the last two terms proceeds just as it did for the zeroth and first moments. Explicitly, we have

$$\sum_{m=0}^{\infty} m^2 \vec{p}(m-1) = \sum_{m=-1}^{\infty} (m+1)^2 \vec{p}(m) = \sum_{m=0}^{\infty} (m+1)^2 \vec{p}(m) = \langle \vec{m}^2 \rangle + 2 \langle \vec{m} \rangle + \langle \vec{m}^0 \rangle, \tag{S37}$$

for the first sum and

$$\sum_{m=0}^{\infty} m^2 (m+1) \vec{p}(m+1) = \sum_{m=1}^{\infty} (m-1)^2 m \vec{p}(m) = \sum_{m=0}^{\infty} (m-1)^2 m \vec{p}(m) = \langle \vec{m}^3 \rangle - 2 \langle \vec{m}^2 \rangle + \langle \vec{m} \rangle \tag{S38}$$

for the second. Substituting the results of the sums back in Eq. S36 gives

$$0 = (\mathbf{K} - \mathbf{R_D})\langle \vec{m}^2 \rangle - \gamma \langle \vec{m}^3 \rangle + \mathbf{R_A}(\langle \vec{m}^2 \rangle + 2\langle \vec{m} \rangle + \langle \vec{m}^0 \rangle) + \gamma(\langle \vec{m}^3 \rangle - 2\langle \vec{m}^2 \rangle + \langle \vec{m} \rangle), \quad \text{(S39)}$$

and after grouping like powers of $m$ we have

$$0 = (\mathbf{K} - \mathbf{R_D} + \mathbf{R_A} - 2\gamma)\langle \vec{m}^2 \rangle + (2\mathbf{R_A} + \gamma)\langle \vec{m} \rangle + \mathbf{R_A}\langle \vec{m}^0 \rangle. \quad \text{(S40)}$$

As we found when computing $\langle m \rangle$ from $\langle \vec{m} \rangle$, we can spare ourselves some algebra by multiplying Eq. S40 by $\vec{1}^\dagger$, which then reduces to

$$0 = -2\gamma \langle m^2 \rangle + \vec{1}^\dagger \cdot (2\mathbf{R_A} + \gamma)\langle \vec{m} \rangle + \vec{1}^\dagger \cdot \mathbf{R_A}\langle \vec{m}^0 \rangle, \quad \text{(S41)}$$

and noting from Eq. S34 that $\vec{1}^\dagger \cdot \mathbf{R_A}\langle \vec{m}^0 \rangle = \gamma \langle m \rangle$, we have the tidy result

$$\langle m^2 \rangle = \langle m \rangle + \frac{1}{\gamma}\vec{1}^\dagger \cdot \mathbf{R_A}\langle \vec{m} \rangle. \quad \text{(S42)}$$

Finally we have all the preliminary results needed to write a general expression for the Fano factor $\nu$. The Fano factor is defined as the ratio of variance to mean, which can be written as

$$\nu = \frac{\langle m^2 \rangle - \langle m \rangle^2}{\langle m \rangle} = \frac{\langle m \rangle + \frac{1}{\gamma}\vec{1}^\dagger \cdot \mathbf{R_A}\langle \vec{m} \rangle - \langle m \rangle^2}{\langle m \rangle} \quad \text{(S43)}$$

and simplified to

$$\nu = 1 - \langle m \rangle + \frac{\vec{1}^\dagger \cdot \mathbf{R_A}\langle \vec{m} \rangle}{\gamma \langle m \rangle}. \quad \text{(S44)}$$

Note a subtle notational trap here: $\langle m \rangle = \frac{1}{\gamma}\vec{1}^\dagger \cdot \mathbf{R_A}\langle \vec{m}^0 \rangle$ rather than the by-eye similar but wrong expression $\langle m \rangle \neq \frac{1}{\gamma}\vec{1}^\dagger \cdot \mathbf{R_A}\langle \vec{m} \rangle$, so the last term in Eq. S44 is in general quite nontrivial. For a generic promoter, Eq. S44 may be greater than, less than, or equal to one, as asserted in Section 3. We have not found the general form Eq. S44 terribly intuitive and instead defer discussion to specific examples.

##### S1.3.4 Summary of general results

For ease of reference, we collect and reprint here the key results derived in this section that are used in the main text and subsequent subsections. Mean mRNA copy number and Fano factor are given by Eqs. S34 and S44, which are

$$\langle m \rangle = \frac{1}{\gamma}\vec{1}^\dagger \cdot \mathbf{R_A}\langle \vec{m}^0 \rangle \quad \text{(S45)}$$

and

$$\nu = 1 - \langle m \rangle + \frac{\vec{1}^\dagger \cdot \mathbf{R_A}\langle \vec{m} \rangle}{\gamma \langle m \rangle}, \quad \text{(S46)}$$

respectively. To compute these two quantities, we need the expressions for $\langle \vec{m}^0 \rangle$ and $\langle \vec{m} \rangle$ given by solving Eqs. S24 and S30, respectively, which are

$$(\mathbf{K} - \mathbf{R_D} + \mathbf{R_A})\langle \vec{m}^0 \rangle = 0 \quad \text{(S47)}$$

and

$$(\mathbf{K} - \mathbf{R_D} + \mathbf{R_A} - \gamma \mathbf{I})\langle \vec{m} \rangle = -\mathbf{R_A} \langle \vec{m}^0 \rangle. \tag{S48}$$

Some comments are in order before we consider particular models. First, note that to obtain $\langle \vec{m} \rangle$ and $\nu$, we need not bother solving for all components of the vectors $\langle \vec{m}^0 \rangle$ and $\langle \vec{m} \rangle$, but only the components which are multiplied by nonzero elements of $\mathbf{R_A}$. The only component of $\langle \vec{m}^0 \rangle$ that ever survives is the transciptionally active state, and for the models we consider here, there is only ever one such state. This will save us some amount of algebra below.

Also note that we are computing Fano factors to verify the results of Section 3, concerning the constitutive promoter models in Figure 2 which are analogs of the simple repression models in Figure 1. We can translate the matrices from the simple repression models to the constitutive case by simply substituting all occurrences of repressor rates by zero and removing the row and column corresponding to the repressor bound state. The results for $\langle m \rangle$ computed in the repressed case can be easily translated to the constitutive case, rather than recalculating from scratch, by taking the limit $k_R^+ \to 0$, since this amounts to sending repressor copy number to zero.

Finally, we point out that it would be possible to compute $\langle \vec{m}^0 \rangle$ more simply using the diagram methods from King and Altman [40] (also independently discovered by Hill [41]). But to our knowledge this method cannot be applied to compute $\langle \vec{m} \rangle$ or $\nu$, so since we would need to resort to solving the matrix equations anyways for $\langle \vec{m} \rangle$, we choose not to introduce the extra conceptual burden of the diagram methods simply for computing $\langle \vec{m}^0 \rangle$.

#### S1.4 Nonequilibrium Model One - Poisson Promoter

##### S1.4.1 Mean mRNA

For nonequilibrium model 1 in Figure 1, we have already shown the full master equation in Eq. 10 and Eq. S6, but for completeness we reprint it again as

$$\begin{aligned}
\frac{d}{dt} p_R(m,t) = &- \overbrace{k_R^- p_R(m,t)}^{R\to U} + \overbrace{k_R^+ p_U(m,t)}^{U\to R} + \overbrace{(m+1)\gamma p_R(m+1,t)}^{m+1\to m} - \overbrace{\gamma m p_R(m,t)}^{m\to m-1} \\
\frac{d}{dt} p_U(m,t) = &\ \overbrace{k_R^- p_R(m,t)}^{R\to U} - \overbrace{k_R^+ p_U(m,t)}^{U\to R} + \overbrace{r p_U(m-1,t)}^{m-1\to m} - \overbrace{r p_U(m,t)}^{m\to m+1} \\
&+ \overbrace{(m+1)\gamma p_U(m+1,t)}^{m+1\to m} - \overbrace{\gamma m p_U(m,t)}^{m\to m-1}.
\end{aligned} \tag{S49}$$

This is a direct transcription of the states and rates in Figure 1. This may be converted to the matrix form of the master equation shown in Eq. S11 with matrices

$$\vec{p}(m) = \begin{pmatrix} p_R(m) \\ p_U(m) \end{pmatrix}, \ \mathbf{K} = \begin{pmatrix} -k_R^- & k_R^+ \\ k_R^- & -k_R^+ \end{pmatrix}, \ \mathbf{R} = \begin{pmatrix} 0 & 0 \\ 0 & r \end{pmatrix}, \tag{S50}$$

where $\mathbf{R_A}$ and $\mathbf{R_D}$ are equal, so we drop the subscript and denote both simply by $\mathbf{R}$. Since our interest is only in steady-state we dropped the time dependence as well.

First we need $\langle \vec{m}^0 \rangle$. Label its components as $p_R$ and $p_U$, the probabilities of finding the system in either promoter state, and note that only $p_U$ survives multiplication by $\mathbf{R}$, since

$$\mathbf{R}\langle \vec{m}^0 \rangle = \begin{pmatrix} 0 & 0 \\ 0 & r \end{pmatrix} \begin{pmatrix} p_R \\ p_U \end{pmatrix} = \begin{pmatrix} 0 \\ r p_U \end{pmatrix}, \tag{S51}$$

so we need not bother finding $p_R$. Then we have

$$(\mathbf{K} - \mathbf{R_D} + \mathbf{R_A})\langle \vec{m}^0 \rangle = \begin{pmatrix} -k_R^- & k_R^+ \\ k_R^- & -k_R^+ \end{pmatrix} \begin{pmatrix} p_R \\ p_U \end{pmatrix} = 0. \tag{S52}$$

As mentioned earlier in Section S1.3.1, the two rows are linearly dependent, so taking only the first row and using normalization to set $p_R = 1 - p_U$ gives

$$-k_R^-(1 - p_U) + k_R^+ p_U = 0, \tag{S53}$$

which is easily solved to find

$$p_U = \frac{k_R^-}{k_R^- + k_R^+}. \tag{S54}$$

Substituting this into Eq. S51, and the result of that into Eq. S45, we have

$$\langle m \rangle = \frac{r}{\gamma} \frac{k_R^-}{k_R^- + k_R^+} \tag{S55}$$

as asserted in Eq. 13 of the main text.

##### S1.4.2 Fano factor

To verify that the Fano factor for model 1 in Figure 2(A) is in fact 1 as claimed in the main text, note that in this limit $p_U = 1$ and $\langle m \rangle = r/\gamma$. All elements of $\mathbf{K}$ are zero, and $\mathbf{R_A} - \mathbf{R_D} = 0$, so Eq. S48 reduces to

$$-\gamma \langle \vec{m} \rangle = -r, \tag{S56}$$

or, in other words, since there is only one promoter state, $\langle \vec{m} \rangle = \langle m \rangle$. Then it follows that

$$\nu = 1 - \frac{r}{\gamma} + \frac{r\langle m \rangle}{\gamma \langle m \rangle} = 1 \tag{S57}$$

as claimed.

#### S1.5 Nonequilibrium Model Two - RNAP Bound and Unbound States

##### S1.5.1 Mean mRNA

As shown earlier, the full master equation for model 2 in Figure 1 is

$$
\begin{aligned}
\frac{d}{dt}p_R(m,t) &= -\overbrace{k_R^- p_R(m,t)}^{R\to U} + \overbrace{k_R^+ p_E(m,t)}^{U\to R} + \overbrace{(m+1)\gamma p_R(m+1,t)}^{m+1\to m} - \overbrace{\gamma m p_R(m,t)}^{m\to m-1} \\
\frac{d}{dt}p_E(m,t) &= \overbrace{k_R^- p_R(m,t)}^{R\to U} - \overbrace{k_R^+ p_E(m,t)}^{U\to R} + \overbrace{(m+1)\gamma p_E(m+1,t)}^{m+1\to m} - \overbrace{\gamma m p_E(m,t)}^{m\to m-1}. \\
&\quad + \overbrace{k_P^- p_P(m,t)}^{A\to U} - \overbrace{k_P^+ p_E(m,t)}^{U\to A} + \overbrace{r p_P(m-1,t)}^{m-1\to m,\ A\to U} \\
\frac{d}{dt}p_P(m,t) &= -\overbrace{k_P^- p_P(m,t)}^{A\to U} + \overbrace{k_P^+ p_E(m,t)}^{U\to A} + \overbrace{(m+1)\gamma p_P(m+1,t)}^{m+1\to m} - \overbrace{\gamma m p_P(m,t)}^{m\to m-1}. \\
&\quad - \overbrace{r p_P(m,t)}^{m\to m+1,\ A\to U},
\end{aligned}
\tag{S58}
$$

which can be condensed to the matrix form of Eq. S11 with matrices given by

$$
\mathbf{K} = \begin{pmatrix} -k_R^- & k_R^+ & 0 \\ k_R^- & -k_R^+ - k_P^+ & k_P^- \\ 0 & k_P^+ & -k_P^- \end{pmatrix},\ \mathbf{R_A} = \begin{pmatrix} 0 & 0 & 0 \\ 0 & 0 & r \\ 0 & 0 & 0 \end{pmatrix},\ \mathbf{R_D} = \begin{pmatrix} 0 & 0 & 0 \\ 0 & 0 & 0 \\ 0 & 0 & r \end{pmatrix}. \tag{S59}
$$

As for model 1, we must first find $\mathbf{R_A}\langle\vec{m}^0\rangle$. Denote its components as $p_R$, $p_E$, $p_P$, the probabilities of being found in repressor bound, empty, or RNAP-bound states, respectively. Only $p_P$ is necessary to find since

$$
\mathbf{R_A}\langle\vec{m}^0\rangle = \begin{pmatrix} 0 \\ r p_P \\ 0 \end{pmatrix}. \tag{S60}
$$

Then Eq. S47 for $\langle\vec{m}\rangle$ reads

$$
\begin{pmatrix} -k_R^- & k_R^+ & 0 \\ k_R^- & -k_R^+ - k_P^+ & k_P^- + r \\ 0 & k_P^+ & -k_P^- - r \end{pmatrix} \begin{pmatrix} p_R \\ p_E \\ p_P \end{pmatrix} = 0. \tag{S61}
$$

Discarding the middle row as redundant and incorporating the normalization condition leads to a set of three linearly independent equations, namely

$$
-k_R^- p_R + k_R^+ p_E = 0 \tag{S62}
$$

$$
k_P^+ p_E + (-k_P^- - r) p_P = 0 \tag{S63}
$$

$$
p_R + p_E + p_P = 1. \tag{S64}
$$

Using $p_R = 1 - p_E - p_P$ to eliminate $p_R$ in the first and solving the resulting equation for $p_E$ gives $p_E = (1-p_P)k_R^-/(k_R^- + k_R^+)$. Substituting this for $p_E$ in the second equation gives an equation in $p_P$ alone which is

$$k_P^+ k_R^-(1-p_P) - (k_P^- + r)(k_R^+ + k_R^-)p_P = 0 \tag{S65}$$

and solving for $p_P$ gives

$$p_P = \frac{k_P^+ k_R^-}{k_P^+ k_R^- + (k_P^- + r)(k_R^+ + k_R^-)}. \tag{S66}$$

Substituting this in Eq. S60 and multiplying by $\mathbf{R_A}$ produces

$$\mathbf{R_A}\langle \vec{m}^0 \rangle = r\frac{k_P^+ k_R^-}{k_P^+ k_R^- + (k_P^- + r)(k_R^+ + k_R^-)} \begin{pmatrix} 0 \\ 1 \\ 0 \end{pmatrix} \tag{S67}$$

from which $\langle m \rangle$ follows readily,

$$\langle m \rangle = \frac{r}{\gamma}\frac{k_P^+ k_R^-}{k_P^+ k_R^- + (k_P^- + r)(k_R^+ + k_R^-)}, \tag{S68}$$

as claimed in Eq. 18 in the main text.

##### S1.5.2 Fano factor

To compute the Fano factor, we first remove the repressor bound state from the matrices describing the model, which reduce to

$$\mathbf{K} = \begin{pmatrix} -k_P^+ & k_P^- \\ k_P^+ & -k_P^- \end{pmatrix},\ \mathbf{R_A} = \begin{pmatrix} 0 & r \\ 0 & 0 \end{pmatrix},\ \mathbf{R_D} = \begin{pmatrix} 0 & 0 \\ 0 & r \end{pmatrix}. \tag{S69}$$

Similarly we remove the repressor bound state from $\mathbf{R_A}\langle \vec{m}^0 \rangle$ and take the $k_R^+ \to 0$ limit, which simplifies to

$$\mathbf{R_A}\langle \vec{m}^0 \rangle = r\frac{k_P^+}{k_P^+ + k_P^- + r} \begin{pmatrix} 1 \\ 0 \end{pmatrix}. \tag{S70}$$

Then we must compute $\langle \vec{m} \rangle$ from Eq. S48, which with these matrices reads

$$(\mathbf{K} - \mathbf{R_D} + \mathbf{R_A} - \gamma\mathbf{I})\langle \vec{m} \rangle = \begin{pmatrix} -k_P^+ - \gamma & k_P^- + r \\ k_P^+ & -k_P^- - r - \gamma \end{pmatrix} \begin{pmatrix} m_E \\ m_P \end{pmatrix} = r\frac{k_P^+}{k_P^+ + k_P^- + r} \begin{pmatrix} 1 \\ 0 \end{pmatrix}, \tag{S71}$$

where we labeled the components of $\langle \vec{m} \rangle$ as $m_E$ and $m_P$, since they are the mean mRNA counts conditional upon the system residing in the empty or polymerase bound states, respectively. Unlike for $\langle \vec{m}^0 \rangle$, the rows of this matrix are linearly independent so we simply solve this matrix equation as is. We can immediately eliminate $m_E$ since $m_E = m_P(k_P^- + r + \gamma)/k_P^+$ from the second row, and substituting into the first row gives an equation for $m_P$ alone, which is

$$\left[-(k_P^+ + \gamma)(k_P^- + r + \gamma) + k_P^+(k_P^- + r)\right] m_P = -\frac{r(k_P^+)^2}{k_P^+ + k_P^- + r}. \tag{S72}$$

Expanding the products cancels several terms, and solving for $m_P$ gives

$$m_P = \frac{r(k_P^+)^2}{\gamma(k_P^+ + k_P^- + r)(k_P^+ + k_P^- + r + \gamma)}. \tag{S73}$$

Note then that $\vec{1}^\dagger \cdot \mathbf{R_A}\langle \vec{m} \rangle = r m_P$. We also need the constitutive limit of $\langle m \rangle$ from Eq. S68, again found by taking $k_R^+ \to 0$, which is

$$\langle m \rangle = \frac{r}{\gamma} \frac{k_P^+}{k_P^+ + k_P^- + r} \tag{S74}$$

and substituting this along with $\vec{1}^\dagger \cdot \mathbf{R_A}\langle \vec{m} \rangle = r m_P$ into Eq. S46 for the Fano factor $\nu$, we find

$$\nu = 1 - \frac{r}{\gamma}\frac{k_P^+}{k_P^+ + k_P^- + r} + \frac{r}{\gamma}\frac{r(k_P^+)^2}{\gamma(k_P^+ + k_P^- + r)(k_P^+ + k_P^- + r + \gamma)}\left(\frac{r}{\gamma}\frac{k_P^+}{k_P^+ + k_P^- + r}\right)^{-1}. \tag{S75}$$

This simplifies to

$$\nu = 1 - \frac{r}{\gamma}\left(\frac{k_P^+}{k_P^+ + k_P^- + r} - \frac{k_P^+}{k_P^+ + k_P^- + r + \gamma}\right), \tag{S76}$$

which further simplifies to

$$\nu = 1 - \frac{r k_P^+}{(k_P^+ + k_P^- + r)(k_P^+ + k_P^- + r + \gamma)}, \tag{S77}$$

exactly Eq. 36 in the main text.

#### S1.6 Nonequilibrium Model Three - Multistep Transcription Initiation and Escape

##### S1.6.1 Mean mRNA

In close analogy to model 2 above, nonequilibrium model 3 from Figure 1(C) can be described by our generic master equation Eq. S11 with promoter transition matrix given by

$$\mathbf{K} = \begin{pmatrix} -k_R^- & k_R^+ & 0 & 0 \\ k_R^- & -k_R^+ - k_P^+ & k_P^- & 0 \\ 0 & k_P^+ & -k_P^- - k_O & 0 \\ 0 & 0 & k_O & 0 \end{pmatrix} \tag{S78}$$

and transcription matrices given by

$$\mathbf{R_A} = \begin{pmatrix} 0 & 0 & 0 & 0 \\ 0 & 0 & 0 & r \\ 0 & 0 & 0 & 0 \\ 0 & 0 & 0 & 0 \end{pmatrix}, \ \mathbf{R_D} = \begin{pmatrix} 0 & 0 & 0 & 0 \\ 0 & 0 & 0 & 0 \\ 0 & 0 & 0 & 0 \\ 0 & 0 & 0 & r \end{pmatrix}. \tag{S79}$$

$\langle \vec{m}^0 \rangle$ is again given by Eq. S47, which in this case takes the form

$$(\mathbf{K} - \mathbf{R_D} + \mathbf{R_A})\langle \vec{m}^0 \rangle = \begin{pmatrix} -k_R^- & k_R^+ & 0 & 0 \\ k_R^- & -k_R^+ - k_P^+ & k_P^- & r \\ 0 & k_P^+ & -k_P^- - k_O & 0 \\ 0 & 0 & k_O & -r \end{pmatrix} \begin{pmatrix} p_R \\ p_E \\ p_C \\ p_O \end{pmatrix} = 0, \tag{S80}$$

where the four components of $\langle \vec{m}^0 \rangle$ correspond to the four promoter states repressor bound, empty, RNAP-bound closed complex, and RNAP-bound open complex. As explained in Section S1.3.1, we are free to discard one linearly dependent row from this matrix and replace it with the normalization condition $p_R + p_E + p_C + p_O = 1$. Using normalization to eliminate $p_R$ from the first row gives

$$p_E = (1 - p_C - p_O) \frac{k_R^-}{k_R^- + k_R^+}. \tag{S81}$$

If we substitute this in the third row, then the last two rows constitute two equations in $p_C$ and $p_O$ given by

$$k_P^+ k_R^- (1 - p_C - p_O) - (k_P^- + k_O)(k_R^+ + k_R^-) p_C = 0 \tag{S82}$$

$$k_O p_C - r p_O = 0. \tag{S83}$$

Solving for $p_C = p_O r / k_O$ in the second and substituting into the first gives us our desired single equation in the single variable $p_O$, which is

$$k_P^+ k_R^- - k_P^+ k_R^- \left(1 + \frac{r}{k_O}\right) p_O - (k_P^- + k_O)(k_R^+ + k_R^-) \frac{r}{k_O} p_O = 0, \tag{S84}$$

and solving for $p_O$ we find

$$p_O = \frac{k_P^+ k_R^- k_O}{k_P^+ k_R^- k_O + r k_P^+ k_R^- + r(k_P^- + k_O)(k_R^+ + k_R^-)}. \tag{S85}$$

Once again $p_O$, the transcriptionally active state, is the only component of $\langle \vec{m}^0 \rangle$ that survives multiplication by $\mathbf{R_A}$, and $\mathbf{R_A}\langle \vec{m}^0 \rangle = r p_O$. So

$$\langle m \rangle = \frac{1}{\gamma} \vec{1}^\dagger \cdot \mathbf{R_A} \langle \vec{m}^0 \rangle = \frac{r}{\gamma} \frac{k_P^+ k_R^- k_O}{k_P^+ k_R^- k_O + r k_P^+ k_R^- + r(k_P^- + k_O)(k_R^+ + k_R^-)}, \tag{S86}$$

which equals Eq. 25 in the main text.

##### S1.6.2 Fano factor

To compute the Fano factor of the analogous constitutive promoter, we first excise the repressor states and rates from the problem. More precisely, we construct the matrix $(\mathbf{K} - \mathbf{R_D} + \mathbf{R_A} - \gamma \mathbf{I})$ and substitute it into Eq. S48 which is now

$$(\mathbf{K} - \mathbf{R_D} + \mathbf{R_A} - \gamma \mathbf{I})\langle \vec{m} \rangle = \begin{pmatrix} -k_P^+ - \gamma & k_P^- & r \\ k_P^+ & -k_P^- - k_O - \gamma & 0 \\ 0 & k_O & -r - \gamma \end{pmatrix} \begin{pmatrix} m_E \\ m_C \\ m_O \end{pmatrix} = -r p_O \begin{pmatrix} 1 \\ 0 \\ 0 \end{pmatrix} \tag{S87}$$

where we labeled the unbound, closed complex, and open complex components of $\langle \vec{m} \rangle$ as $m_E$, $m_C$, and $m_O$, respectively. $p_O$ is given by the limit of Eq. S85 as $k_R^+ \to 0$, which is

$$p_O = \frac{k_P^+ k_O}{k_P^+(k_O + r) + r(k_P^- + k_O)} \equiv \frac{k_P^+ k_O}{\mathcal{Z}}, \tag{S88}$$

where we define $\mathcal{Z}$ for upcoming convenience as this sum of terms will appear repeatedly. We can use the second equation to eliminate $m_E$, finding $m_E = m_C(k_P^- + k_O + \gamma)/k_P^+$, and the third to eliminate $m_C$, which is simply $m_C = m_O(r + \gamma)/k_O$. Substituting these both into the first equation gives a single equation for the variable of interest, $m_O$,

$$-(k_P^+ + \gamma)(k_P^- + k_O + \gamma)(r + \gamma)m_O + k_P^- k_P^+(r + \gamma)m_O + r k_P^+ k_O m_O = -r k_P^+ k_O p_O, \tag{S89}$$

which is solved for $m_O$ to give

$$m_O = p_O \frac{r k_P^+ k_O}{(k_P^+ + \gamma)(k_P^- + k_O + \gamma)(r + \gamma) - r k_P^+ k_O - k_P^- k_P^+(r + \gamma)}. \tag{S90}$$

Expanding the denominator and canceling terms leads to

$$m_O = p_O \frac{r}{\gamma} \frac{k_P^+ k_O}{\mathcal{Z} + \gamma(k_P^+ + k_P^- + k_O + r) + \gamma^2}. \tag{S91}$$

Now $\vec{1}^\dagger \cdot \mathbf{R_A} \langle \vec{m} \rangle = r m_O$, and $\langle m \rangle = r p_O/\gamma$, so if we substitute these two quantities into Eq. S46, we will readily obtain the Fano factor as

$$\nu = 1 - \langle m \rangle + \frac{\vec{1}^\dagger \cdot \mathbf{R_A} \langle \vec{m} \rangle}{\gamma \langle m \rangle} = 1 - \frac{r}{\gamma} p_O + \frac{m_O}{p_O}. \tag{S92}$$

Substituting, we see that

$$\nu = 1 - \frac{r}{\gamma} \frac{k_P^+ k_O}{\mathcal{Z}} + \frac{r}{\gamma} \frac{k_P^+ k_O}{\mathcal{Z} + \gamma(k_P^+ + k_P^- + k_O + r) + \gamma^2}, \tag{S93}$$

and after simplifying, we obtain

$$\nu = 1 - \frac{r k_P^+ k_O}{\mathcal{Z}} \frac{k_P^+ + k_P^- + k_O + r + \gamma}{\mathcal{Z} + \gamma(k_P^+ + k_P^- + k_O + r) + \gamma^2}, \tag{S94}$$

as stated in Eq. 37 in the main text.

##### S1.6.3 Generalizing $\nu < 1$ to more fine-grained models

In the main text we argued that the convolution of multiple exponential distributions should be a distribution with a smaller fractional width than the corresponding exponential distribution with the same mean. This can be made more precise with a result from [76], who showed that the convolution of multiple gamma distributions (of which the exponential distribution is a special case) is, to a very good approximation, also gamma distributed. Using their Eq. 2 for the distribution of the convolution, with shape parameters set to 1 to give exponential distributions, the total waiting time distribution has a ratio of variance to squared mean $\sigma^2/\mu^2 = \sum_i k_i^2/\left(\sum_i k_i\right)^2$, where the $k_i$ are the rates of the individual steps. Clearly this is less than 1 and therefore the total waiting time distribution is narrower than the corresponding exponential.

We also claimed in the main text that for a process whose waiting time distribution is narrower than exponential, i.e., has $\sigma^2/\mu^2 < 1$, the distribution of counts should be less variable than a Poisson distribution, leading to a Fano factor $\nu < 1$. This we argue by analogy to photon statistics where it is known that "antibunched" arrivals, in other words more uniformly distributed in time relative to uncorrelated arrivals, generally gives rise to sub-Poissonian noise [77], [78]. Although loopholes to this result are known to exist, to our knowledge they appear to arise from uniquely quantum effects so we do not expect they apply for our problem. Nevertheless we refrain from elevating this equivalence of kinetic cycles with sub-Poissonian noise to a "theorem."

#### S1.7 Nonequilibrium Model Four - "Active" and "Inactive" States

##### S1.7.1 Mean mRNA

The mathematical specification of this model is almost identical to model 2. The matrix $\mathbf{K}$ is identical, as is $\mathbf{R_D}$. The only difference is that now $\mathbf{R_A} = \mathbf{R_D}$, i.e., both are diagonal, in contrast to model 2 where $\mathbf{R_A}$ has an off-diagonal element, as in Eq. S59. Then the analog of Eq. S61 for finding $\langle m^0 \rangle$ is

$$\begin{pmatrix} -k_R^- & k_R^+ & 0 \\ k_R^- & -k_R^+ - k^+ & k^- \\ 0 & k^+ & -k^- \end{pmatrix} \begin{pmatrix} p_R \\ p_I \\ p_A \end{pmatrix} = 0. \tag{S95}$$

In fact we need not do this calculation explicitly and can instead recycle the calculation of mean mRNA $\langle m \rangle$ from model 2. The matrices are identical except for the relabeling $k^- \longleftrightarrow (k_P^- + r)$, and a careful look through the derivation of $\langle m \rangle$ for model 2 shows that the parameters $k_P^-$ and $r$ only ever appear as the sum $k_P^- + r$. So taking $\langle m \rangle$ from model 2, Eq. S68, and relabeling $(k_P^- + r) \to k^-$ gives us our answer for model four, simply

$$\langle m \rangle = \frac{r}{\gamma} \frac{k^+ k_R^-}{k^+ k_R^- + k^-(k_R^+ + k_R^-)}. \tag{S96}$$

##### S1.7.2 Fano factor

Likewise, for computing the Fano factor of this model we may take a shortcut. Consider the constitutive model four from Figure 2 for which we want to compute the Fano factor and compare it to nonequilibrium model one of simple repression in Figure 1. Mathematically these are exactly the same model, just with rates labeled differently and the meaning of the promoter states interpreted differently. Furthermore, nonequilibrium model 1 from Figure 1 was the model considered by Jones et. al. [36], where they derived the Fano factor for that model to be

$$\nu = 1 + \frac{r k_R^+}{(k_R^+ + k_R^-)(k_R^+ + k_R^- + \gamma)}. \tag{S97}$$

So recognizing that the relabelings $k_R^+ \to k^-$ and $k_R^- \to k^+$ will translate this result to our model four from Figure 2, we can immediately write down our Fano factor as

$$\nu = 1 + \frac{rk^-}{(k^- + k^+)(k^- + k^+ + \gamma)}, \tag{S98}$$

as quoted in Eq. 38 and in Figure 2.

### S2 Bursty promoter models - generating function solutions and numerics

#### S2.1 Constitutive promoter with bursts

##### S2.1.1 From master equation to generating function

The objective of this section is to write down the steady-state mRNA distribution for model 5 in Figure 2. Our claim is that this model is rich enough that it can capture the expression pattern of bacterial constitutive promoters. Figure S2 shows two different schematic representations of the model. Figure S2(A) shows the promoter cartoon model with burst initiation rate $k_i$, mRNA degradation rate $\gamma$, and mean burst size $b$. For our derivation of the chemical master equation we will focus more on Figure S2(B). This representation is intended to highlight that bursty gene expression allows transitions between mRNA count $m$ and $m'$ even with $m - m' > 1$.

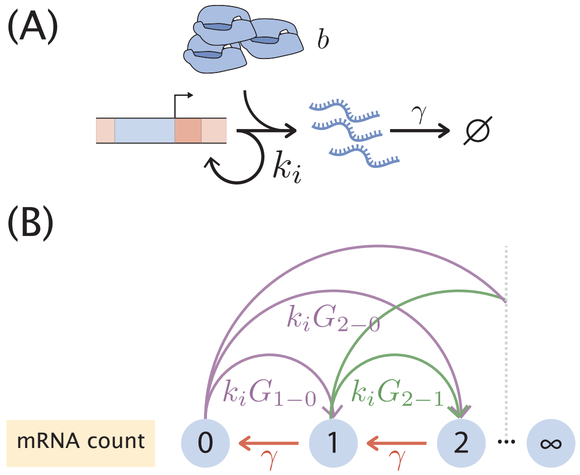

**Figure S2. Bursty transcription for unregulated promoter.** (A) Schematic of the one-state bursty transcription model. Rate $k_i$ is the bursty initiation rate, $\gamma$ is the mRNA degradation rate, and $b$ is the mean burst size. (B) Schematic depiction of the mRNA count state transitions. The model in (A) allows for transitions of $> 1$ mRNA counts with probability $G_{m-m'}$, where the state jumps from having $m'$ mRNA to having $m$ mRNA in a single burst of gene expression.

To derive the master equation we begin by considering the possible state transitions to "enter" state $m$. There are two possible paths to jump from an mRNA count $m' \neq m$ to a state $m$ in a small time window $\Delta t$:

1. By degradation of a single mRNA, jumping from $m+1$ to $m$.
2. By producing $m-m'$ mRNA for $m' \in \{0, 1, \ldots, m-1\}$.

For the "exit" states from $m$ into $m' \neq m$ during a small time window $\Delta t$ we also have two possibilities:

1. By degradation of a single mRNA, jumping from $m$ to $m-1$.
2. By producing $m'-m$ mRNA for $m'-m \in \{1, 2, \ldots\}$.

This implies that the probability of having $m$ mRNA at time $t+\Delta t$ can be written as

$$\begin{aligned} p(m, t+\Delta t) =& p(m, t) + \overbrace{\gamma \Delta t (m+1) p(m+1, t)}^{m+1 \to m} - \overbrace{\gamma \Delta t m p(m, t)}^{m \to m-1} \\ &+ \overbrace{k_i \Delta t \sum_{m'=0}^{m-1} G_{m-m'} p(m', t)}^{m' \in \{0,1,\ldots m-1\} \to m} - \overbrace{k_i \Delta t \sum_{m'=m+1}^{\infty} G_{m'-m} p(m, t)}^{m \to m' \in \{m+1, m+2, \ldots\}}, \end{aligned} \tag{S99}$$

where we indicate $G_{m'-m}$ as the probability of having a burst of size $m'-m$, i.e. when the number of mRNAs jump from $m$ to $m' > m$ due to a single mRNA transcription burst. We suggestively use the letter $G$ as we will assume that these bursts sizes are geometrically distributed with parameter $\theta$. This is written as

$$G_k = \theta(1-\theta)^k \text{ for } k \in \{0, 1, 2, \ldots\}. \tag{S100}$$

In Section 3 of the main text we derive this functional form for the burst size distribution. An intuitive way to think about it is that for transcription initiation events that take place instantaneously there are two competing possibilities: Producing another mRNA with probability $(1-\theta)$, or ending the burst with probability $\theta$. What this implies is that for a geometrically distributed burst size we have a mean burst size $b$ of the form

$$b \equiv \langle m' - m \rangle = \sum_{k=0}^{\infty} k\theta(1-\theta)^k = \frac{1-\theta}{\theta}. \tag{S101}$$

To clean up Equation S99 we can send the first term on the right hand side to the left, and divide both sides by $\Delta t$. Upon taking the limit where $\Delta t \to 0$ we can write

$$\frac{d}{dt} p(m,t) = (m+1)\gamma p(m+1,t) - m\gamma p(m,t) + k_i \sum_{m'=0}^{m-1} G_{m-m'} p(m',t) - k_i \sum_{m'=m+1}^{\infty} G_{m'-m} p(m,t). \tag{S102}$$

Furthermore, given that the timescale for this equation is set by the mRNA degradation rate $\gamma$ we can divide both sides by this rate, obtaining

$$\frac{d}{d\tau} p(m,\tau) = (m+1) p(m+1,\tau) - m p(m,\tau) + \lambda \sum_{m'=0}^{m-1} G_{m-m'} p(m',\tau) - \lambda \sum_{m'=m+1}^{\infty} G_{m'-m} p(m,\tau), \tag{S103}$$

where we defined $\tau \equiv t \times \gamma$, and $\lambda \equiv k_i/\gamma$. The last term in Eq. S103 sums all burst sizes except for a burst of size zero. We can re-index the sum to include this term, obtaining

$$\lambda \sum_{m'=m+1}^{\infty} G_{m'-m} p(m,\tau) = \lambda p(m,t) \left[ \underbrace{\sum_{m'=m}^{\infty} G_{m'-m}}_{\text{re-index sum to include burst size zero}} - \underbrace{G_0}_{\text{subtract extra added term}} \right]. \tag{S104}$$

Given the normalization constraint of the geometric distribution, adding the probability of all possible burst sizes – including size zero since we re-indexed the sum – allows us to write

$$\sum_{m'=m}^{\infty} G_{m'-m} - G_0 = 1 - G_0. \tag{S105}$$

Substituting this into Eq. S103 results in

$$\frac{d}{d\tau} p(m,\tau) = (m+1)p(m+1,\tau) - mp(m,\tau) + \lambda \sum_{m'=0}^{m-1} G_{m-m'} p(m',\tau) - \lambda p(m,\tau) \left[1 - G_0\right]. \tag{S106}$$

To finally get at a more compact version of the equation notice that the third term in Eq. S106 includes burst from size $m' - m = 1$ to size $m' - m = m$. We can include the term $p(m,t)G_0$ in the sum which allows bursts of size $m' - m = 0$. This results in our final form for the chemical master equation

$$\frac{d}{d\tau} p(m,\tau) = (m+1)p(m+1,\tau) - mp(m,\tau) - \lambda p(m,\tau) + \lambda \sum_{m'=0}^{m} G_{m-m'} p(m',\tau). \tag{S107}$$

In order to solve Eq. S107 we will use the generating function method [79]. The probability generating function is defined as

$$F(z,t) = \sum_{m=0}^{\infty} z^m p(m,t), \tag{S108}$$

where $z$ is just a dummy variable that will help us later on to obtain the moments of the distribution. Let us now multiply both sides of Eq. S107 by $z^m$ and sum over all $m$

$$\sum_m z^m \frac{d}{d\tau} p(m,\tau) = \sum_m z^m \left[ -mp(m,\tau) + (m+1)p(m+1,\tau) + \lambda \sum_{m'=0}^{m} G_{m-m'} p(m',\tau) - \lambda p(m,\tau) \right], \tag{S109}$$

where we use $\sum_m \equiv \sum_{m=0}^{\infty}$. We can distribute the sum and use the definition of $F(z,t)$ to obtain

$$\frac{dF(z,\tau)}{d\tau} = -\sum_m z^m mp(m,\tau) + \sum_m z^m (m+1)p(m+1,\tau) + \lambda \sum_m z^m \sum_{m'=0}^{m} G_{m-m'} p(m',\tau) - \lambda F(z,\tau). \tag{S110}$$

We can make use of properties of the generating function to write everything in terms of $F(z, \tau)$: the first term on the right hand side of Eq. S110 can be rewritten as

$$\sum_m z^m \cdot m \cdot p(m, \tau) = \sum_m z \frac{\partial z^m}{\partial z} p(m, \tau), \tag{S111}$$

$$= \sum_m z \frac{\partial}{\partial z} \left( z^m p(m, \tau) \right), \tag{S112}$$

$$= z \frac{\partial}{\partial z} \left( \sum_m z^m p(m, \tau) \right), \tag{S113}$$

$$= z \frac{\partial F(z, \tau)}{\partial z}. \tag{S114}$$

For the second term on the right hand side of Eq. S110 we define $k \equiv m + 1$. This allows us to write

$$\sum_{m=0}^{\infty} z^m \cdot (m+1) \cdot p(m+1, \tau) = \sum_{k=1}^{\infty} z^{k-1} \cdot k \cdot p(k, \tau), \tag{S115}$$

$$= z^{-1} \sum_{k=1}^{\infty} z^k \cdot k \cdot p(k, \tau), \tag{S116}$$

$$= z^{-1} \sum_{k=0}^{\infty} z^k \cdot k \cdot p(k, \tau), \tag{S117}$$

$$= z^{-1} \left( z \frac{\partial F(z)}{\partial z} \right), \tag{S118}$$

$$= \frac{\partial F(z)}{\partial z}. \tag{S119}$$

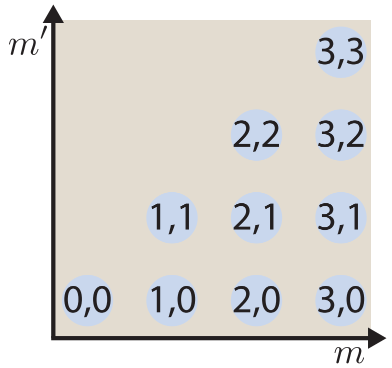

**Figure S3. Reindexing double sum.** Schematic for reindexing the sum $\sum_{m=0}^{\infty} \sum_{m'=0}^{m}$. Blue circles depict the 2D grid of nonnegative integers restricted to the lower triangular part of the $m, m'$ plane. The trick is that this double sum runs over all $(m, m')$ pairs with $m' \leq m$. Summing $m$ first instead of $m'$ requires determining the boundary: the upper boundary of the $m'$-first double sum becomes the lower boundary of the $m$-first double sum.

The third term in Eq. S110 is the most trouble. The trick is to reverse the default order of the sums as

$$\sum_{m=0}^{\infty} \sum_{m'=0}^{m} = \sum_{m'=0}^{\infty} \sum_{m=m'}^{\infty} . \tag{S120}$$

To see the logic of the sum we point the reader to Figure S3. The key is to notice that the double sum $\sum_{m=0}^{\infty} \sum_{m'=0}^{m}$ is adding all possible pairs $(m, m')$ in the lower triangle, so we can add the terms vertically as the original sum indexing suggests, i.e.

$$\sum_{m=0}^{\infty} \sum_{m'=0}^{m} x_{(m,m')} = x_{(0,0)} + x_{(1,0)} + x_{(1,1)} + x_{(2,0)} + x_{(2,1)} + x_{(2,2)} + \ldots, \tag{S121}$$

where the variable $x$ is just a placeholder to indicate the order in which the sum is taking place. But we can also add the terms horizontally as

$$\sum_{m'=0}^{\infty} \sum_{m=m'}^{\infty} x_{(m,m')} = x_{(0,0)} + x_{(1,0)} + x_{(2,0)} + \ldots + x_{(1,1)} + x_{(2,1)} + \ldots, \tag{S122}$$

which still adds all of the lower triangle terms. Applying this reindexing results in

$$\lambda \sum_{m} z^m \sum_{m'=0}^{m} G_{m-m'} p(m', \tau) = \lambda \sum_{m'=0}^{\infty} \sum_{m=m'}^{\infty} z^m \theta (1-\theta)^{m-m'} p(m', \tau), \tag{S123}$$

where we also substituted the definition of the geometric distribution $G_k = \theta(1-\theta)^k$. Redistributing the sums we can write

$$\lambda \sum_{m'=0}^{\infty} \sum_{m=m'}^{\infty} z^m \theta (1-\theta)^{m-m'} p(m', \tau) = \lambda\theta \sum_{n=0}^{\infty} (1-\theta)^{m'} P(m', \tau) \sum_{m=m'}^{\infty} [z(1-\theta)]^m . \tag{S124}$$

The next step requires us to look slightly ahead into what we expect to obtain. We are working on deriving an equation for the generating function $F(z, \tau)$ that when solved will allow us to compute what we care about, i.e. the probability function $p(m, \tau)$. Upon finding the function for $F(z, \tau)$, we will recover this probability distribution by evaluating derivatives of $F(z, \tau)$ at $z = 0$, whereas we can evaluate derivatives of $F(z, \tau)$ at $z = 1$ to instead recover the moments of the distribution. The point here is that when the dust settles we will evaluate $z$ to be less than or equal to one. Furthermore, we know that the parameter of the geometric distribution $\theta$ must be strictly between zero and one. With these two facts we can safely state that $|z(1-\theta)| < 1$. Defining $n \equiv m - m'$ we rewrite the last sum in Eq. S124 as

$$\sum_{m=m'}^{\infty} [z(1-\theta)]^m = \sum_{n=0}^{\infty} [z(1-\theta)]^{n+m'} \tag{S125}$$

$$= [z(1-\theta)]^{m'} \sum_{n=0}^{\infty} [z(1-\theta)]^n \tag{S126}$$

$$= [z(1-\theta)]^{m'} \left( \frac{1}{1 - z(1-\theta)} \right), \tag{S127}$$

where we use the geometric series since, as stated before, $|z(1-\theta)| < 1$. Putting these results together, the PDE for the generating function is

$$\frac{\partial F}{\partial \tau} = \frac{\partial F}{\partial z} - z\frac{\partial F}{\partial z} - \lambda F + \frac{\lambda \theta F}{1 - z(1-\theta)}. \tag{S128}$$

Changing variables to $\xi = 1 - \theta$ and simplifying gives

$$\frac{\partial F}{\partial \tau} + (z-1)\frac{\partial F}{\partial z} = \frac{(z-1)\xi}{1 - z\xi}\lambda F. \tag{S129}$$

##### S2.1.2 Steady-state

To get at the mRNA distribution at steady state we first must solve Eq. S129 setting the time derivative to zero. At steady-state, the PDE reduces to the ODE

$$\frac{dF}{dz} = \frac{\xi}{1 - z\xi}\lambda F, \tag{S130}$$

which we can integrate as

$$\int \frac{dF}{F} = \int \frac{\lambda \xi dz}{1 - \xi z}. \tag{S131}$$

The initial conditions for generating functions can be subtle and confusing. The key fact follows from the definition $F(z,t) = \sum_m z^m p(m,t)$. Clearly normalization of the distribution requires that $F(z = 1, t) = \sum_m p(m,t) = 1$. A subtlety is that sometimes the generating function may be undefined *at* $z = 1$, in which case the limit as $z$ approaches 1 from below suffices to define the normalization condition. We also warn the reader that, while it is frequently convenient to change variables from $z$ to a different independent variable, one must carefully track how the normalization condition transforms.

Continuing on, we evaluate the integrals (producing a constant $c$) which gives

$$\ln F = -\lambda \ln(1 - \xi z) + c \tag{S132}$$

$$F = \frac{c}{(1 - \xi z)^{\lambda}}. \tag{S133}$$

Only one choice for $c$ can satisfy initial conditions, producing

$$F(z) = \left(\frac{1-\xi}{1-\xi z}\right)^{\lambda} = \left(\frac{\theta}{1 - z(1-\theta)}\right)^{\lambda}, \tag{S134}$$

##### S2.1.3 Recovering the steady-state probability distribution

To obtain the steady state mRNA distribution $p(m)$ we are aiming for we need to extract it from the generating function

$$F(z) = \sum_m z^m p(m). \tag{S135}$$

Taking a derivative with respect to $z$ results in

$$\frac{dF(z)}{dz} = \sum_m m z^{m-1} p(m). \tag{S136}$$

Setting $z = 0$ leaves one term in the sum when $m = 1$

$$\left.\frac{dF(z)}{dz}\right|_{z=0} = \left(0 \cdot 0^{-1} \cdot p(0) + 1 \cdot 0^0 \cdot p(1) + 2 \cdot 0^1 \cdot p(2) + \cdots\right) = p(1), \tag{S137}$$

since in the limit $\lim_{x \to 0^+} x^x = 1$. A second derivative of the generating function would result in

$$\frac{d^2F(z)}{dz^2} = \sum_{m=0}^{\infty} m(m-1) z^{m-2} p(m). \tag{S138}$$

Again evaluating at $z = 0$ gives

$$\left.\frac{d^2F(z)}{dz}\right|_{z=0} = 2p(z). \tag{S139}$$

In general any $p(m)$ is obtained from the generating function as

$$p(m) = \frac{1}{m!} \left.\frac{d^mF(z)}{dz}\right|_{z=0}. \tag{S140}$$

Let's now look at the general form of the derivative for our generating function in Eq. S134. For $p(0)$ we simply evaluate $F(z = 0)$ directly, obtaining

$$p(0) = F(z = 0) = \theta^\lambda. \tag{S141}$$

The first derivative results in

$$\begin{aligned} \frac{dF(z)}{dz} &= \theta^\lambda \frac{d}{dz}(1 - z(1-\theta))^{-\lambda} \\ &= \theta^\lambda \left[-\lambda(1 - z(1-f))^{-\lambda-1} \cdot (\theta - 1)\right] \\ &= \theta^\lambda \left[\lambda(1 - z(1-\theta))^{-\lambda-1}(1-\theta)\right]. \end{aligned} \tag{S142}$$

Evaluating this at $z = 0$ as required to get $p(1)$ gives

$$\left.\frac{dF(z)}{dz}\right|_{z=0} = \theta^\lambda \lambda (1-\theta) \tag{S143}$$

For the second derivative we find

$$\frac{d^2F(z)}{dz^2} = \theta^\lambda \left[\lambda(\lambda+1)(1 - z(1-\theta))^{-\lambda-2}(1-\theta)^2\right]. \tag{S144}$$

Again evaluating $z = 0$ gives

$$\left.\frac{d^2F(z)}{dz^2}\right|_{z=0} = \theta^\lambda \lambda(\lambda+1)(1-\theta)^2. \tag{S145}$$

Let's go for one more derivative to see the pattern. The third derivative of the generating function gives

$$\frac{d^3F(z)}{dz^3} = \theta^\lambda \left[\lambda(\lambda+1)(\lambda+2)(1-z(1-\theta))^{-\lambda-3}(1-\theta)^3\right], \tag{S146}$$

which again we evaluate at $z = 0$

$$\left.\frac{d^3F(z)}{dz^3}\right|_{z=1} = \theta^\lambda \left[\lambda(\lambda+1)(\lambda+2)(1-\theta)^3\right]. \tag{S147}$$

If $\lambda$ was an integer we could write this as

$$\left.\frac{d^3F(z)}{dz^3}\right|_{z=0} = \frac{(\lambda+2)!}{(\lambda-1)!}\theta^\lambda(1-\theta)^3. \tag{S148}$$

Since $\lambda$ might not be an integer we can write this using Gamma functions as

$$\left.\frac{d^3F(z)}{dz^3}\right|_{z=0} = \frac{\Gamma(\lambda+3)}{\Gamma(\lambda)}\theta^\lambda(1-\theta)^3. \tag{S149}$$

Generalizing the pattern we then have that the $m$-th derivative takes the form

$$\left.\frac{d^mF(z)}{dz^m}\right|_{z=0} = \frac{\Gamma(\lambda+m)}{\Gamma(\lambda)}\theta^\lambda(1-\theta)^m. \tag{S150}$$

With this result we can use Eq. S140 to obtain the desired steady-state probability distribution function

$$p(m) = \frac{\Gamma(m+\lambda)}{\Gamma(m+1)\Gamma(\lambda)}\theta^\lambda(1-\theta)^m. \tag{S151}$$

Note that the ratio of gamma functions is often expressed as a binomial coefficient, but since $\lambda$ may be non-integer, this would be ill-defined. Re-expressing this exclusively in our variables of interest, burst rate $\lambda$ and mean burst size $b$, we have

$$p(m) = \frac{\Gamma(m+\lambda)}{\Gamma(m+1)\Gamma(\lambda)}\left(\frac{1}{1+b}\right)^\lambda\left(\frac{b}{1+b}\right)^m. \tag{S152}$$

#### S2.2 Adding repression

##### S2.2.1 Deriving the generating function for mRNA distribution

Let us move from a one-state promoter to a two-state promoter, where one state has repressor bound and the other produces transcriptional bursts as above. A schematic of this model is shown as model 5 in Figure 1(C). Although now we have an equation for each promoter state, otherwise the master equation reads similarly to the one-state case, except with additional terms corresponding to transitions between promoter states, namely

$$\frac{d}{dt}p_R(m,t) = k_R^+ p_A(m,t) - k_R^- p_R(m,t) + (m+1)\gamma p_R(m+1,t) - m\gamma p_R(m,t) \tag{S153}$$

$$\begin{aligned}\frac{d}{dt}p_A(m,t) =& -k_R^+ p_A(m,t) + k_R^- p_R(m,t) + (m+1)\gamma p_A(m+1,t) - m\gamma p_A(m,t) \\ & - k_i p_A(m,t) + k_i \sum_{m'=0}^{m} \theta(1-\theta)^{m-m'} p_A(m',t),\end{aligned} \tag{S154}$$

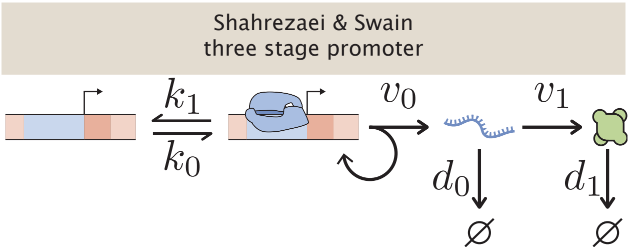

**Figure S4. Schematic of three-stage promoter from [23].** Adapted from Shahrezaei & Swain [23]. In their paper they derive a closed form solution for the protein distribution. Our two-state bursty promoter at the mRNA level can be mapped into their solution with some relabeling.

where $p_R(m, t)$ is the probability of the system having $m$ mRNA copies and having repressor bound to the promoter at time $t$, and $p_A$ is an analogous probability to find the promoter without repressor bound. $k_R+$ and $k_R^-$ are, respectively, the rates at which repressors bind and unbind to and from the promoter, and $\gamma$ is the mRNA degradation rate. $k_i$ is the rate at which bursts initiate, and as before, the geometric distribution of burst sizes has mean $b = (1 - \theta)/\theta$.

Interestingly, it turns out that this problem maps exactly onto the three-stage promoter model considered by Shahrezaei and Swain in [23], with relabelings. Their approximate solution for protein distributions amounts to the same approximation we make here in regarding the duration of mRNA synthesis bursts as instantaneous, so their solution for protein distributions also solves our problem of mRNA distributions. Let us examine the analogy more closely. They consider a two-state promoter, as we do here, but they model mRNA as being produced one at a time and degraded, with rates $v_0$ and $d_0$. Then they model translation as occurring with rate $v_1$, and protein degradation with rate $d_1$ as shown in Figure S4. Now consider the limit where $v_1, d_0 \to \infty$ with their ratio $v_1/d_0$ held constant. $v_1/d_0$ resembles the average burst size of translation from a single mRNA: these are the rates of two Poisson processes that compete over a transcript, which matches the story of geometrically distributed burst sizes. In other words, in our bursty promoter model we can think of the parameter $\theta$ as determining one competing process to end the burst and $(1-\theta)$ as a process wanting to continue the burst. So after taking this limit, on timescales slow compared to $v_1$ and $d_0$, it appears that transcription events fire at rate $v_0$ and produce a geometrically distributed burst of translation of mean size $v_1/d_0$, which intuitively matches the story we have told above for mRNA with variables relabeled.

To verify this intuitively conjectured mapping between our problem and the solution in [23], we continue with a careful solution for the mRNA distribution using probability generating functions, following the ideas sketched in [23]. It is natural to nondimensionalize rates in the problem by $\gamma$, or equivalently, this amounts to measuring time in units of $\gamma^{-1}$. We are also only interested in steady state, so we set the time derivatives to zero, giving

$$0 = k_R^+ p_A(m) - k_R^- p_R(m) + (m+1)p_R(m+1) - m p_R(m) \tag{S155}$$

$$\begin{aligned} 0 =& - k_R^+ p_A(m) + k_R^- p_R(m) + (m+1)p_A(m+1) - m p_A(m) \\ & - k_i p_A(m) + k_i \sum_{m'=0}^{m} \theta(1-\theta)^{m-m'} p_A(m'), \end{aligned} \tag{S156}$$

where for convenience we kept the same notation for all rates, but these are now expressed in units of mean mRNA lifetime $\gamma^{-1}$.

The probability generating function is defined as before in the constitutive case, except now we must introduce a generating function for each promoter state,

$$f_A(z) = \sum_{m=0}^{\infty} z^m p_A(m), \; f_R(z) = \sum_{m=0}^{\infty} z^m p_R(m). \tag{S157}$$

Our real objective is the generating function $f(z)$ that generates the mRNA distribution $p(m)$, independent of what state the promoter is in. But since $p(m) = p_A(m) + p_R(m)$, it follows too that $f(z) = f_A(z) + f_R(z)$.

As before we multiply both equations by $z^m$ and sum over all $m$. Each individual term transforms exactly as did an analogous term in the constitutive case, so the coupled ODEs for the generating functions read

$$0 = k_R^+ f_A(z) - k_R^- f_R(z) + \frac{\partial}{\partial z} f_R(z) - z \frac{\partial}{\partial z} f_R(z) \tag{S158}$$

$$\begin{aligned} 0 =& - k_R^+ f_A(z) + k_R^- f_R(z) + \frac{\partial}{\partial z} f_A(z) - z \frac{\partial}{\partial z} f_A(z) \\ & - k_i f_A(z) + k_i \frac{\theta}{1 - z(1-\theta)} f_A(z), \end{aligned} \tag{S159}$$

and after changing variables $\xi = 1 - \theta$ as before and rearranging, we have

$$0 = k_R^+ f_A(z) - k_R^- f_R(z) + (1-z) \frac{\partial}{\partial z} f_R(z) \tag{S160}$$

$$0 = -k_R^+ f_A(z) + k_R^- f_R(z) + (1-z) \frac{\partial}{\partial z} f_A(z) + k_i \frac{(z-1)\xi}{1 - z\xi} f_A(z), \tag{S161}$$

We can transform this problem from two coupled first-order ODEs to a single second-order ODE by solving for $f_A$ in the first and plugging into the second, giving

$$\begin{aligned} 0 =& (1-z) \frac{\partial f_R}{\partial z} + \frac{1-z}{k_R^+} \left( k_R^- \frac{\partial f_R}{\partial z} + \frac{\partial f_R}{\partial z} + (z-1) \frac{\partial^2 f_R}{\partial z^2} \right) \\ & + \frac{k_i}{k_R^+} \frac{(z-1)\xi}{1 - z\xi} \left( k_R^- f_R + (z-1) \frac{\partial f_R}{\partial z} \right), \end{aligned} \tag{S162}$$

where, to reduce notational clutter, we have dropped the explicit $z$ dependence of $f_A$ and $f_R$. Simplifying we have

$$0 = \frac{\partial^2 f_R}{\partial z^2} - \left(\frac{k_i \xi}{1 - z\xi} + \frac{1 + k_R^- + k_R^+}{1 - z}\right) \frac{\partial f_R}{\partial z} + \frac{k_i k_R^- \xi}{(1 - z\xi)(1 - z)} f_R. \tag{S163}$$

This can be recognized as the hypergeometric differential equation, with singularities at $z = 1$, $z = \xi^{-1}$, and $z = \infty$. The latter can be verified by a change of variables from $z$ to $x = 1/z$, being careful with the chain rule, and noting that $z = \infty$ is a singular point if and only if $x = 1/z = 0$ is a singular point.

The standard form of the hypergeometric differential equation has its singularities at 0, 1, and $\infty$, so to take advantage of the standard form solutions to this ODE, we first need to transform variables to put it into a standard form. However, this is subtle. While any such transformation should work in principle, the solutions are expressed most simply in the neighborhood of $z = 0$, but the normalization condition that we need to enforce corresponds to $z = 1$. The easiest path, therefore, is to find a change of variables that maps 1 to 0, $\infty$ to $\infty$, and $\xi^{-1}$ to 1. This is most intuitively done in two steps.

First map the $z = 1$ singularity to 0 by the change of variables $v = z - 1$, giving

$$0 = \frac{\partial^2 f_R}{\partial v^2} + \left(\frac{k_i \xi}{(1 + v)\xi - 1} + \frac{1 + k_R^- + k_R^+}{v}\right) \frac{\partial f_R}{\partial v} + \frac{k_i k_R^- \xi}{((1 + v)\xi - 1)v} f_R. \tag{S164}$$

Now two singularities are at $v = 0$ and $v = \infty$. The third is determined by $(1 + v)\xi - 1 = 0$, or $v = \xi^{-1} - 1$. We want another variable change that maps this third singularity to 1 (without moving 0 or infinity). Changing variables again to $w = \frac{v}{\xi^{-1} - 1} = \frac{\xi}{1 - \xi} v$ fits the bill. In other words, the combined change of variables

$$w = \frac{\xi}{1 - \xi}(z - 1) \tag{S165}$$

maps $z = \{1, \xi^{-1}, \infty\}$ to $w = \{0, 1, \infty\}$ as desired. Plugging in, being mindful of the chain rule and noting $(1 + v)\xi - 1 = (1 - \xi)(w - 1)$ gives

$$0 = \left(\frac{\xi}{1 - \xi}\right)^2 \frac{\partial^2 f_R}{\partial w^2} + \left(\frac{\xi k_i}{(1 - \xi)(w - 1)} + \frac{\xi(1 + k_R^- + k_R^+)}{(1 - \xi)w}\right) \frac{\xi}{1 - \xi} \frac{\partial f_R}{\partial w} + \frac{k_i k_R^- \xi^2}{(1 - \xi)^2 w(w - 1)} f_R. \tag{S166}$$

This is close to the standard form of the hypergeometric differential equation, and some cancellation and rearrangement gives

$$0 = w(w - 1)\frac{\partial^2 f_R}{\partial w^2} + \left(k_i w + (1 + k_R^- + k_R^+)(w - 1)\right) \frac{\partial f_R}{\partial w} + k_i k_R^- f_R. \tag{S167}$$

and a little more algebra produces

$$0 = w(1 - w)\frac{\partial^2 f_R}{\partial w^2} + \left(1 + k_R^- + k_R^+ - (1 + k_i + k_R^- + k_R^+)w\right) \frac{\partial f_R}{\partial w} - k_i k_R^- f_R, \tag{S168}$$

which is the standard form. From this we can read off the solution in terms of hypergeometric functions ${}_2F_1$ from any standard source, e.g. [80], and identify the conventional parameters in terms of our model parameters. We want the general solution in the neighborhood of $w = 0$ ($z = 1$), which for a homogeneous linear second order ODE must be a sum of two linearly independent solutions. More precisely,

$$f_R(w) = C^{(1)}{}_2F_1(\alpha, \beta, \delta; w) + C^{(2)} w^{1-\delta}{}_2F_1(1 + \alpha - \delta, 1 + \beta - \delta, 2 - \delta; w) \tag{S169}$$

with parameters determined by

$$\begin{aligned} \alpha\beta &= k_i k_R^- \\ 1 + \alpha + \beta &= 1 + k_i + k_R^- + k_R^+ \\ \delta &= 1 + k_R^- + k_R^+ \end{aligned} \tag{S170}$$

and constants $C^{(1)}$ and $C^{(2)}$ to be set by boundary conditions. Solving for $\alpha$ and $\beta$, we find

$$\begin{aligned} \alpha &= \frac{1}{2}\left(k_i + k_R^- + k_R^+ + \sqrt{(k_i + k_R^- + k_R^+)^2 - 4k_i k_R^-}\right) \\ \beta &= \frac{1}{2}\left(k_i + k_R^- + k_R^+ - \sqrt{(k_i + k_R^- + k_R^+)^2 - 4k_i k_R^-}\right) \\ \delta &= 1 + k_R^- + k_R^+. \end{aligned} \tag{S171}$$

Note that $\alpha$ and $\beta$ are interchangeable in the definition of ${}_2F_1$ and differ only in the sign preceeding the radical. Since the normalization condition requires that $f_R$ be finite at $w = 0$, we can immediately set $C^{(2)} = 0$ to discard the second solution. This is because all the rate constants are strictly positive, so $\delta > 1$ and therefore $w^{1-\delta}$ blows up as $w \to 0$. Now that we have $f_R$, we would like to find the generating function for the mRNA distribution, $f(z) = f_A(z) + f_R(z)$. We can recover $f_A$ from our solution for $f_R$, namely

$$f_A(z) = \frac{1}{k_R^+}\left(k_R^- f_R(z) + (z - 1)\frac{\partial f_R}{\partial z}\right) \tag{S172}$$

or

$$f_A(w) = \frac{1}{k_R^+}\left(k_R^- f_R(w) + w\frac{\partial f_R}{\partial w}\right), \tag{S173}$$

where in the second line we transformed our original relation between $f_R$ and $f_A$ to our new, more convenient, variable $w$. Plugging our solution for $f_R(w) = C^{(1)}{}_2F_1(\alpha, \beta, \delta; w)$ into $f_A$, we will require the differentiation rule for ${}_2F_1$, which tells us

$$\frac{\partial f_R}{\partial w} = C^{(1)}\frac{\alpha\beta}{\delta}{}_2F_1(\alpha + 1, \beta + 1, \delta + 1; w), \tag{S174}$$

from which it follows that

$$f_A(w) = \frac{C^{(1)}}{k_R^+}\left(k_R^-{}_2F_1(\alpha, \beta, \delta; w) + w\frac{\alpha\beta}{\delta}{}_2F_1(\alpha + 1, \beta + 1, \delta + 1; w)\right) \tag{S175}$$

and therefore

$$f(w) = C^{(1)}\left(1+\frac{k_R^-}{k_R^+}\right) {}_2F_1(\alpha,\beta,\delta;w) + w\frac{C^{(1)}}{k_R^+}\frac{\alpha\beta}{\delta} {}_2F_1(\alpha+1,\beta+1,\delta+1;w). \tag{S176}$$

To proceed, we need one of the (many) useful identities known for hypergeometric functions, in particular

$$w\frac{\alpha\beta}{\delta} {}_2F_1(\alpha+1,\beta+1,\delta+1;w) = (\delta-1)\left({}_2F_1(\alpha,\beta,\delta-1;w) - {}_2F_1(\alpha,\beta,\delta;w)\right). \tag{S177}$$

Substituting this for the second term in $f(w)$, we find

$$f(w) = \frac{C^{(1)}}{k_R^+}\left[\left(k_R^+ + k_R^-\right) {}_2F_1(\alpha,\beta,\delta;w) + (\delta-1)\left({}_2F_1(\alpha,\beta,\delta-1;w) - {}_2F_1(\alpha,\beta,\delta;w)\right)\right], \tag{S178}$$

and since $\delta - 1 = k_R^+ + k_R^-$, the first and third terms cancel, leaving only

$$f(w) = C^{(1)}\frac{k_R^+ + k_R^-}{k_R^+} {}_2F_1(\alpha,\beta,\delta-1;w). \tag{S179}$$

Now we enforce normalization, demanding $f(w=0) = f(z=1) = 1$. ${}_2F_1(\alpha,\beta,\delta-1;0) = 1$, so we must have $C^{(1)} = k_R^+/(k_R^+ + k_R^-)$ and consequently

$$f(w) = {}_2F_1(\alpha,\beta,k_R^+ + k_R^-;w). \tag{S180}$$

Recalling that the mean burst size $b = (1-\theta)/\theta = \xi/(1-\xi)$ and $w = \frac{\xi}{1-\xi}(z-1) = b(z-1)$, we can transform back to the original variable $z$ to find the tidy result

$$f(z) = {}_2F_1(\alpha,\beta,k_R^+ + k_R^-;b(z-1)), \tag{S181}$$

with $\alpha$ and $\beta$ given above by

$$\begin{aligned}\alpha &= \frac{1}{2}\left(k_i + k_R^- + k_R^+ + \sqrt{(k_i + k_R^- + k_R^+)^2 - 4k_ik_R^-}\right)\\ \beta &= \frac{1}{2}\left(k_i + k_R^- + k_R^+ - \sqrt{(k_i + k_R^- + k_R^+)^2 - 4k_ik_R^-}\right).\end{aligned} \tag{S182}$$

Finally we are in sight of the original goal. We can generate the steady-state probability distribution of interest by differentiating the generating function,

$$p(m) = m!\left.\frac{\partial^m}{\partial z^m}f(z)\right|_{z=0}, \tag{S183}$$

which follows easily from its definition. Some contemplation reveals that repeated application of the derivative rule used above will produce products of the form $\alpha(\alpha+1)(\alpha+2)\cdots(\alpha+m-1)$ in the expression for $p(m)$ and similarly for $\beta$ and $\delta$. These resemble ratios of factorials, but since $\alpha$, $\beta$, and $\delta$ are not necessarily integer, we should express the ratios using gamma functions instead. More precisely, one finds

$$p(m) = \frac{\Gamma(\alpha+m)\Gamma(\beta+m)\Gamma(k_R^+ + k_R^-)}{\Gamma(\alpha)\Gamma(\beta)\Gamma(k_R^+ + k_R^- + m)}\frac{b^m}{m!} {}_2F_1(\alpha+m,\beta+m,k_R^+ + k_R^- + m;-b) \tag{S184}$$

which is finally the probability distribution we sought to derive.

#### S2.3 Numerical considerations and recursion formulas

##### S2.3.1 Generalities

We would like to carry out Bayesian parameter inference on FISH data from [36], using Eq. (S184) as our likelihood. This requires accurate (and preferably fast) numerical evaluation of the hypergeometric function ${}_2F_1$, which is a notoriously hard problem [81], [82], and our particular needs here present an especial challenge as we show below.

The hypergeometric function is defined by its Taylor series as

$$ {}_2F_1(a,b,c;z) = \sum_{l=0}^{\infty} \frac{\Gamma(a+l)\Gamma(b+l)\Gamma(c)}{\Gamma(a)\Gamma(b)\Gamma(c+l)} \frac{z^l}{l!} \tag{S185} $$

for $|z| < 1$, and by analytic continuation elsewhere. If $z \lesssim 1/2$ and $\alpha$ and $\beta$ are not too large (absolute value below 20 or 30), then the series converges quickly and an accurate numerical representation is easily computed by truncating the series after a reasonable number of terms. Unfortunately, we need to evaluate ${}_2F_1$ over mRNA copy numbers fully out to the tail of the distribution, which can easily reach 50, possibly 100. From Eq. (S184), this means evaluating ${}_2F_1$ repeatedly for values of $a$, $b$, and $c$ spanning the full range from $\mathcal{O}(1)$ to $\mathcal{O}(10^2)$, even if $\alpha$, $\beta$, and $\delta$ in Eq. (S184) are small, with the situation even worse if they are not small. A naive numerical evaluation of the series definition will be prone to overflow and, if any of $a, b, c < 0$, then some successive terms in the series have alternating signs which can lead to catastrophic cancellations.

One solution is to evaluate ${}_2F_1$ using arbitrary precision arithmetic instead of floating point arithmetic, e.g., using the `mpmath` library in Python. This is accurate but incredibly slow computationally. To quantify how slow, we found that evaluating the likelihood defined by Eq. (S184) $\sim 50$ times (for a typical dataset of interest from [36], with $m$ values spanning 0 to $\sim 50$) using arbitrary precision arithmetic is 100-1000 fold slower than evaluating a negative binomial likelihood for the corresponding constitutive promoter dataset.

To claw back $\gtrsim 30$ fold of that slowdown, we can exploit one of the many catalogued symmetries involving ${}_2F_1$. The solution involves recursion relations originally explored by Gauss, and studied extensively in [81], [82]. They are sometimes known as contiguous relations and relate the values of any set of 3 hypergeometric functions whose arguments differ by integers. To rephrase this symbolically, consider a set of hypergeometric functions indexed by an integer $n$,

$$ f_n = {}_2F_1(a + \epsilon_i n, b + \epsilon_j n, c + \epsilon_k n; z), \tag{S186} $$

for a fixed choice of $\epsilon_i, \epsilon_j, \epsilon_k \in \{0, \pm 1\}$ (at least one of $\epsilon_i, \epsilon_j, \epsilon_k$ must be nonzero, else the set of $f_n$ would contain only a single element). Then there exist known recurrence relations of the form

$$ A_n f_{n-1} + B_n f_n + C_n f_{n+1} = 0, \tag{S187} $$

where $A_n, B_n$, and $C_n$ are some functions of $a, b, c$, and $z$. In other words, for fixed $\epsilon_i, \epsilon_j, \epsilon_k, a, b$, and $c$, if we can merely evaluate ${}_2F_1$ twice, say for $n'$ and $n' - 1$, then we can easily and rapidly generate values for arbitrary $n$.

This provides a convenient solution for our problem: we need repeated evaluations of ${}_2F_1(a+m, b+m, c+m; z)$ for fixed $a, b$, and $c$ and many integer values of $m$. They idea is that we can use arbitrary precision arithmetic to evaluate ${}_2F_1$ for just two particular values of $m$ and then generate ${}_2F_1$ for the other 50-100 values of $m$ using the recurrence relation. In fact there are even more sophisticated ways of utilizing the recurrence relations that might have netted another factor of 2 speed-up, and possibly as much as a factor of 10, but the method described here had already reduced the computation time to an acceptable $\mathcal{O}(1\text{ min})$, so these more sophisticated approaches did not seem worth the time to pursue.

However, there are two further wrinkles. The first is that a naive application of the recurrence relation is numerically unstable. Roughly, this is because the three term recurrence relations, like second order ODEs, admit two linearly independent solutions. In a certain eigenbasis, one of these solutions dominates the other as $n \to \infty$, and as $n \to -\infty$, the dominance is reversed. If we fail to work in this eigenbasis, our solution of the recurrence relation will be a mixture of these solutions and rapidly accumulate numerical error. For our purposes, it suffices to know that the authors of [82] derived the numerically stable solutions (so-called *minimal solutions*) for several possible choices of $\epsilon_i, \epsilon_j, \epsilon_k$. Running the recurrence in the proper direction using a minimal solution is numerically robust and can be done entirely in floating point arithmetic, so that we only need to evaluate ${}_2F_1$ with arbitrary precision arithmetic to generate the seed values for the recursion.

The second wrinkle is a corollary to the first. The minimal solutions are only minimal for certain ranges of the argument $z$, and not all of the 26 possible recurrence relations have minimal solutions for all $z$. This can be solved by using one of the many transformation formulae for ${}_2F_1$ to convert to a different recurrence relation that has a minimal solution over the required domain of $z$, although this can require some trial and error to find the right transformation, the right recurrence relation, and the right minimal solution.

##### S2.3.2 Particulars

Let us now demonstrate these generalities for our problem of interest. In order to evaluate the probability distribution of our model, Eq. (S184), we need to evaluate hypergeometric functions of the form ${}_2F_1(\alpha+m, \beta+m, \delta+m; -b)$ for values of $m$ ranging from 0 to $\mathcal{O}(100)$. The authors of [82] did not derive a recursion relation for precisely this case. We could follow their methods and do so ourselves, but it is much easier to convert to a case that they did consider. The strategy is to look through the minimal solutions tabulated in [82] and search for a transformation we could apply to ${}_2F_1(\alpha+m, \beta+m, \delta+m; -b)$ that would place the $m$'s (the variable being incremented by the recursion) in the same arguments of ${}_2F_1$ as the minimal solution. After some “guess and check,” we found that the transformation

$$ {}_2F_1(\alpha+m, \beta+m, \delta+m; -b) = (1+b)^{-\alpha-m} {}_2F_1\left(\alpha+m, \delta-\beta, \delta+m; \frac{b}{1+b}\right), \tag{S188} $$

produces a ${}_2F_1$ on the right hand side that closely resembles the minimal solutions $y_{3,m}$ and $y_{4,m}$ in Eq. 4.3 in [82]. Explicitly, these solutions are

$$ y_{3,m} \propto {}_2F_1\left(-\alpha'+\delta'-m, -\beta'+\delta', 1-\alpha'-\beta'+\delta'-m; 1-z\right) \tag{S189} $$
$$ y_{4,m} \propto {}_2F_1\left(\alpha'+m, \beta', 1+\alpha'+\beta'-\delta'+m; 1-z\right), \tag{S190} $$

where we have omitted prefactors which are unimportant for now. Which of these two we should use depends on what values $z$ takes on. Equating $1-z = b/(1+b)$ gives $z = 1/(1+b)$, and since $b$ is strictly positive, $z$ is bounded between 0 and 1. From Eq. 4.5 in [82], $y_{4,m}$ is the minimal solution for real $z$ satisfying $0 < z < 2$, so this is the only minimal solution we need.

Now that we have our minimal solution, what recurrence relation does it satisfy? Confusingly, the recurrence relation of which $y_{4,m}$ is a solution increments different arguments of ${}_2F_1$ that does $y_{4,m}$: it increments the first only, rather than first and third. This recurrence relation can be looked up, e.g., Eq. 15.2.10 in [80], which is

$$(\delta' - (\alpha' + m))f_{m-1} + (2(\alpha' + m) - \delta' + (\beta' - \alpha')z)f_m + \alpha'(z-1)f_{m+1} = 0. \tag{S191}$$

Now we must solve for the parameters appearing in the recurrence relation in terms of our parameters, namely by setting

$$\begin{aligned} \alpha' &= \alpha \\ \beta' &= \delta - \beta \\ 1 + \alpha' + \beta' - \delta' &= \delta \\ 1 - z &= \frac{b}{1+b} \end{aligned} \tag{S192}$$

and solving to find

$$\begin{aligned} \alpha' &= \alpha \\ \beta' &= \delta - \beta \\ \delta' &= 1 + \alpha - \beta \\ z &= \frac{1}{1+b}. \end{aligned} \tag{S193}$$

Finally we have everything we need. The minimal solution

$$y_{4,m} = \frac{\Gamma(1 + \alpha' - \delta' + m)}{\Gamma(1 + \alpha' + \beta' - \delta' + m)} \times {}_2F_1\left(\alpha' + m, \beta', 1 + \alpha' + \beta' - \delta' + m; 1 - z\right), \tag{S194}$$

where we have now included the necessary prefactors, is a numerically stable solution of the recurrence relation Eq. (S191) if the recursion is run from large $m$ to small $m$.

Let us finally outline the complete procedure as an algorithm to be implemented:

1. Compute the value of ${}_2F_1$ for the two largest $m$ values of interest using arbitrary precision arithmetic.
2. Compute the prefactors to construct $y_{4,\max(m)}$ and $y_{4,\max(m)-1}$.
3. Recursively compute $y_{4,m}$ for all $m$ less than $\max(m)$ down to $m = 0$.
4. Cancel off the prefactors of the resulting values of $y_{4,m}$ for all $m$ to produce ${}_2F_1$ for all desired $m$ values.

With ${}_2F_1$ computed, the only remaining numerical danger in computing $p(m)$ in Eq. (S184) is overflow of the gamma functions. This is easily solved by taking the log of the entire expression and using standard routines to compute the log of the gamma functions, then exponentiating the entire expression at the end if $p(m)$ is needed rather than $\log p(m)$.

### S3 Bayesian inference

#### S3.1 The problem of parameter inference

One could argue that the whole goal of formulating theoretical models about nature is to sharpen our understanding from qualitative statements to precise quantitative assertions about the relevant features of the natural phenomena in question [83]. It is in these models that we intend to distill the essential parts of the object of study. Writing down such models leads to a propagation of mathematical variables that parametrize our models. By assigning numerical values to these parameters we can compute concrete predictions that can be contrasted with experimental data. For these predictions to match the data the parameter values have to carefully be chosen from the whole parameter space. But how do we go about assessing the effectiveness of different regions of parameter space to speak to the ability of our model to reproduce the experimental observations? The language of probability, and more specifically of Bayesian statistics is – we think – the natural language to tackle this question.

##### S3.1.1 Bayes' theorem

Bayes' theorem is a simple mathematical statement that can apply to *any* logical conjecture. For two particular events $A$ and $B$ that potentially depend on each other Bayes' theorem gives us a recipe for how to update our beliefs about one, let us say $B$, given some state of knowledge, or lack thereof, about $A$. In its most classic form Bayes' theorem is written as

$$P(B \mid A) = \frac{P(A \mid B)P(B)}{P(A)}, \tag{S195}$$

where the vertical line | is read as "given that". So $P(B \mid A)$ is read as probability of $B$ given that $A$ took place. $A$ and $B$ can be any logical assertion. In particular the problem of Bayesian inference focuses on the question of finding the probability distribution of a particular parameter value given the data.

For a given model with a set of parameters $\vec{\theta} = (\theta_1, \theta_2, \ldots, \theta_n)$, the so-called *posterior distribution* $P(\vec{\theta} \mid D)$, where $D$ is the experimental data, quantifies the plausibility of a set of parameter values given our observation of some particular dataset. In other words, through the application of Bayes' formula we update our beliefs on the possible values that parameters can take upon learning the outcome of a particular experiment. We specify the word "update" as we come to every inference problem with prior information about the plausibility of particular regions of parameter space even before performing any experiment. Even when we claim as researchers that we are totally ignorant about the values that the parameters in our models can take, we always come to a problem with domain expertise that can be exploited. If this was not the case, it is likely that the formulation of our model is not going to capture the phenomena we claim to want to understand. This prior information is captured in the *prior probability* $P(\vec{\theta})$. The relationship between how parameter values can connect with the data is enconded in the *likelihood function* $P(D \mid \vec{\theta})$. Our theoretical model, whether deterministic or probabilistic, is encoded in this term that can be intuitively understood as the probability of having observed the particular experimental data we have at hand given that our model is parametrized with the concrete values $\vec{\theta}$. Implicitly here we are also conditioning on the fact that our theoretical model is "true," i.e. the model itself if evaluated or simulated in the computer is capable of generating equivalent datasets to the one we got to observe in an experiment. In this way Bayesian inference consists of applying Bayes' formula as

$$P(\vec{\theta} \mid D) \propto P(D \mid \vec{\theta}) P(\vec{\theta}). \tag{S196}$$

Notice than rather than writing the full form of Bayes' theorem, we limit ourselves to the terms that depend on our quantity of interest – that is the parameter values themselves $\vec{\theta}$ – as the denominator $P(D)$ only serves as a normalization constant.

We also emphasize that the dichotomy we have presented between prior and likelihood is more subtle. Although it is often stated that our prior knowledge is entirely encapsulated by the obviously named prior probability $P(\vec{\theta})$, this is usually too simplistic. The form(s) we choose for our likelihood function $P(D \mid \vec{\theta})$ also draw heavily on our prior domain expertise and the assumptions, implicit and explicit, that these choices encode are at least as important, and often inseparable from, the prior probability, as persuasively argued in [84].

##### S3.1.2 The likelihood function

As we alluded in the previous section it is through the likelihood function $P(D \mid \vec{\theta})$ that we encode the connection between our parameter values and the experimental observables. Broadly speaking there are two classes of models that we might need to encode into our likelihood function:

- Deterministic models: Models for which a concrete selection of parameter values give a single output. Said differently, models with a one-to-one mapping between inputs and outputs.
- Probabilistic models: As the name suggests, models that, rather than having a one-to-one input-output mapping, describe the full probability distribution of possible outputs.

In this paper we focus on inference done with probabilistic models. After all, the chemical master equations we wrote down describe the time evolutions of the mRNA probability distribution. So all our terms $P(\vec{\theta} \mid D)$ will be given by the steady-state solution of the corresponding chemical master equation in question. This is rather convenient as we do not have to worry about adding a statistical model on top of our model to describe deviations from the predictions. Instead our models themselves focus on predicting such variation in cell count.

##### S3.1.3 Prior selection

The different models explored in this work embraced different levels of coarse-graining that resulted in a diverse number of parameters for different models. For each of these model configurations Bayes' theorem demands from us to represent our preconceptions on the possible parameter values in the form of the prior $P(\vec{\theta})$. Throughout this work for models with $> 1$ parameter we assign independent priors to each of the parameters; this is

$$P(\vec{\theta}) = \prod_{i=1}^{n} P(\theta_i). \tag{S197}$$

Although it is not uncommon practice to use non-informative, or maximally uninformative priors, we are of the mindset that this is a disservice to the philosophical and practical implications of Bayes' theorem. It sounds almost contradictory to claim that can we represent our thinking about a natural phenomenon in the form of a mathematical model – in the context of Bayesian inference this means choosing a form for the likehihoods, and even making this choice presupposes prior understanding or assumptions as to the relevant features in the system under study – but that we have absolutely no idea what the parameter values could or could not be. We therefore make use of our own expertise, many times in the form of order-of-magnitude estimates, to write down weakly-informative prior distributions for our parameters.

For our particular case all of the datasets from [36] used in this paper have $\mathcal{O}(10^3)$ data points. What this implies is that our particular choice of priors will not significantly affect our inference as long as they are broad enough. A way to see why this is the case is to simply look at Bayes' theorem. For $N$ $1000 - 3000$ datum all of the independent of each other and $n \ll 10^3$ parameters Bayes' theorem reads as

$$P(\vec{\theta} \mid D) \propto \prod_{k=1}^{N} P(d_k \mid \vec{\theta}) \prod_{i=1}^{n} P(\theta_i), \tag{S198}$$

where $d_k$ represents the $k$-th datum. That means that if our priors span a wide range of parameter space, the posterior distribution would be dominated by the likelihood function.

##### S3.1.4 Expectations and marginalizations

For models with more than one or two parameters, it is generally difficult to visualize or reason about the full joint posterior distribution $P(\vec{\theta} \mid D)$ directly. One of the great powers of Bayesian analysis is *marginalization*, allowing us to reduce the dimensionality to only the parameters of immediate interest by averaging over the other dimensions. Formally, for a three dimensional model with parameters $\theta_1$, $\theta_2$, and $\theta_3$, we can for instance marginalize away $\theta_3$ to produce a 2D posterior as

$$P(\theta_1, \theta_2 \mid D) \propto \int_{\theta_3} d\theta_3 \, P(\theta_1, \theta_2, \theta_3 \mid D), \tag{S199}$$

or we can marginalize away $\theta_1$ and $\theta_3$ to produce the 1D marginal posterior of $\theta_2$ alone, which would be

$$P(\theta_2 \mid D) \propto \int_{\theta_1} d\theta_1 \int_{\theta_3} d\theta_3 \, P(\theta_1, \theta_2, \theta_3 \mid D). \tag{S200}$$

Conceptually, this is what we did in generating the 2D slices of the full 9D model in Figure 4(A). In practice, this marginalization is even easier with Markov Chain Monte Carlo samples in hand. Since each point is simply a list of parameter values, we simply ignore the parameters which we want to marginalize away [52].

###### S3.1.5 Markov Chain Monte Carlo

The theory and practice of Bayesian inference with Markov Chain Monte Carlo (MCMC) is a rich subject with fascinating and deep analogies to statistical mechanics, even drawing on classical Hamiltonian mechanics and general relativity in its modern incarnations. We refer the interested reader to [52] and [85] for excellent introductions. Here we merely give a brief summary of the MCMC computations carried out in this work.

We used the Python package `emcee` for most of the MCMC sampling in this work. For the constitutive promoter inference, we also ran sampling with the excellent Stan modeling language as a check. We did not use Stan for the inference of the simple repression model because implementing the gradients of the hypergeometric function ${}_2F_1$ appearing in Eq. S184, the probability distribution for our bursty model with repression, would have been an immensely challenging task. `emcee` was more than adequate for our purposes, and we were perhaps lucky that the 9-D posterior model for the model of simple repression with bursty promoter was quite well behaved and did not require the extra power of the Hamiltonian Monte Carlo algorithm provided by Stan [86]. Source code for all statistical inference will be made available at `https://github.com/RPGroup-PBoC/bursty_transcription`.

#### S3.2 Bayesian inference on constitutive promoters

Having introduced the ideas behind Bayesian inference we are ready to apply the theoretical machinery to our non-equilibrium models. In particular in this section we will focus on model 1 and model 5 in Figure 2(A). Model 1, the Poisson promoter, will help us build practical intuition into the implementation of the Bayesian inference pipeline as we noted in Section 3 of the main text that this model cannot be reconciled with experimental data from observables such as the Fano factor. In other words, we acknowledge that this model is "wrong," but we still see value in going through the analysis since the simple nature of the model translates into a neat statistical analysis.

###### S3.2.1 Model 1 - Poisson promoter

As specified in the main test, the mRNA steady-state distribution for model 1 in Figure 2(A) is Poisson with parameter $\lambda$. Throughout this Appendix we will appeal to the convenient notation for probability distributions of the form

$$m \sim \text{Poisson}(\lambda), \tag{S201}$$

where the simbol "∼" can be read as *is distributed according to.* So the previous equation can be read as: the mRNA copy number $m$ is distributed according to a Poisson distribution with parameter $\lambda$. Our objective then is to compute the posterior probability distribution $P(\lambda \mid D)$, where, as in the main text, $D = \{m_1, m_2, \ldots, m_N\}$ are the data consisting of single-cell mRNA counts. Since we can assume that each of the cells mRNA counts are independent of any other cells, our likelihood function $P(D \mid \lambda)$ consists of the product of $N$ Poisson distributions.

To proceed with the inference problem we need to specify a prior. In this case we are extremely data-rich, as the dataset from Jones et. al [36] has of order 1000-3000 single-cell measurements for each promoter, so our choice of prior matters little here, as long as it is sufficiently broad. A convenient choice for our problem is to use a *conjugate* prior. A conjugate prior is a special prior that causes the posterior to have the same functional form as the prior, simply with updated model parameters. This makes calculations analytically tractable and also offers a nice interpretation of the inference procedure as updating our knowledge about the model parameters. This makes conjugate priors very useful when they exist. The caveat is that conjugate priors only exist for a very limited number of likelihoods, mostly with only one or two model parameters, so in almost all other Bayesian inference problems, we must tackle the posterior numerically.

But, for the problem at hand, a conjugate prior does in fact exist. For a Poisson likelihood of identical and identically distributed data, the conjugate prior is a gamma distribution, as can be looked up in, e.g., [52], Section 2.6. Putting a gamma prior on $\lambda$ introduces two new parameters $\alpha$ and $\beta$ which parametrize the gamma distribution itself, which we use to encode the range of $\lambda$ values we view as reasonable. Recall $\lambda$ is the mean steady-state mRNA count per cell, which *a priori* could plausibly be anywhere from 0 to a few hundred. $\alpha = 1$ and $\beta = 1/50$ achieve this, since the gamma distribution is strictly positive with mean $\alpha/\beta$ and standard deviation $\sqrt{\alpha}/\beta$. To be explicit, then, our prior is

$$\lambda \sim \text{Gamma}(\alpha, \beta) \tag{S202}$$

As an aside, note that if we did not know that our prior was a conjugate prior, we could still write down our posterior distribution from its definition as

$$p(\lambda \mid D, \alpha, \beta) \propto p(D \mid \lambda) p(\lambda \mid \alpha, \beta) \propto \left( \prod_{k=1}^{N} \frac{\lambda^{m_k} e^{-\lambda}}{m_k!} \right) \frac{\beta}{\Gamma(\alpha)} (\beta\lambda)^{\alpha-1} e^{-\beta\lambda}. \tag{S203}$$

Without foreknowledge that this in fact reduces to a gamma distribution, this expression might appear rather inscrutable. When conjugate priors are unavailable for the likelihood of interest - which is almost always the case for models with > 1 model parameter - this inscrutability is the norm, and making sense of posteriors analytically is almost always impossible. Fortunately, MCMC sampling provides us a powerful method of constructing posteriors numerically which we will make use of extensively.

Since we did use a conjugate prior, we may simply look up our posterior in any standard reference such as [52], Section 2.6, from which we find that

$$\lambda \sim \text{Gamma}\left(\alpha + \bar{m}N, \beta + N\right), \tag{S204}$$

where we defined the sample mean $\bar{m} = \frac{1}{N}\sum_k m_k$ for notational convenience. A glance at the FISH data from [36] reveals that $N$ is $\mathcal{O}(10^3)$ and $\langle m \rangle \gtrsim 0.1$ for all constitutive strains in [36], so $\bar{m}N \gtrsim 10^2$. Therefore as we suspected, our prior parameters are completely overwhelmed by the data. The prior behaves, in a sense, like $\beta$ extra "data points" with a mean value of $(\alpha - 1)/\beta$ [52], which gives us some intuition for how much data is needed to overwhelm the prior in this case: enough data $N$ such that $\beta \ll N$ and $\alpha/\beta \ll \bar{m}$. In fact, $\bar{m}N$ and $N$ are so large that we can, to an excellent approximation, ignore the $\alpha$ and $\beta$ dependence and approximate the gamma distribution as a Gaussian with mean $\bar{m}$ and standard deviation $\sqrt{\bar{m}/N}$, giving

$$\lambda \sim \text{Gamma}\left(\alpha + \bar{m}N, \beta + N\right) \approx \text{Normal}\left(\bar{m}, \sqrt{\frac{\bar{m}}{N}}\right). \tag{S205}$$

As an example with real numbers, for the *lacUV5* promoter, Jones et. al [36] measured 2648 cells with an average mRNA count per cell of $\bar{m} \approx 18.7$. In this case then, our posterior is

$$\lambda \sim \text{Normal}\left(18.7, 0.08\right), \tag{S206}$$

which suggests we have inferred our model's one parameter to a precision of order 1%.

This is not wrong, but it is not the full story. The model's posterior distribution is tightly constrained, but is it a good generative model? In other words, if we use the model to generate synthetic data in the computer does it generate data that look similar to our actual data, and is it therefore plausible that the model captures the important features of the data generating process? This intuitive notion can be codified with *posterior predictive checks*, or PPCs, and we will see that this simple Poisson model fails badly.

The intuitive idea of posterior predictive checks is simple:

1. Make a random draw of the model parameter $\lambda$ from the posterior distribution.
2. Plug that draw into the likelihood and generate a synthetic dataset $\{m_k\}$ conditioned on $\lambda$.
3. Repeat many times.

More formally, the posterior predictive distribution can be thought of as the distribution of future yet-to-be-observed data, conditioned on the data we have already observed. Clearly if those data appear quite different, the model has a problem. Put another way, if we suppose the generative model is true, i.e. we claim that our model explains the process through which our observed experimental data was generated, then the synthetic datasets we generate should resemble the actual observed data. If this is not the case, it suggests the model is missing important features. All the data we consider in this work are 1D (distributions of mRNA counts over a population) so empirical cumulative distribution functions ECDFs are an excellent visual means of comparing synthetic and observed datasets. In general for higher dimensional datasets, much of the challenge is in merely designing good visualizations that can actually show if synthetic and observed data are similar or not.

For our example Poisson promoter model then, we merely draw many random numbers, say 1000, from the Gaussian posterior in Eq. S206. For each one of those draws, we generate a dataset from the likelihood, i.e., we draw 2648 (the number of observed cells in the actual dataset) Poisson-distributed numbers for each of the 1000 posterior draws, for a total of 2648000 samples from the posterior predictive distribution.

To compare so many samples with the actual observed data, one excellent visualization for 1D data is ECDFs of the quantiles, as shown for our Poisson model in Figure 3(B) in the main text.

##### S3.2.2 Model 5 - Bursty promoter

Let us now consider the problem of parameter inference from FISH data for model five from Figure 1(C). As derived in Appendix S2, the steady-state mRNA distribution in this model is a negative binomial distribution, given by

$$p(m) = \frac{\Gamma(m + k_i)}{\Gamma(m + 1)\Gamma(k_i)} \left(\frac{1}{1+b}\right)^{k_i} \left(\frac{b}{1+b}\right)^{m}, \tag{S207}$$

where $b$ is the mean burst size and $k_i$ is the burst rate nondimensionalized by the mRNA degradation rate $\gamma$. As sketched earlier, we can intuitively think about this distribution through a simple story. The story of this distribution is that the promoter undergoes geometrically-distributed bursts of mRNA, where the arrival of bursts is a Poisson process with rate $k_i$ and the mean size of a burst is $b$.

As for the Poisson promoter model, this expression for the steady-state mRNA distribution is exactly the likelihood we want to use in Bayes' theorem. Again denoting the single-cell mRNA count data as $D = \{m_1, m_2, \ldots, m_N\}$, here Bayes' theorem takes the form

$$p(k_i, b \mid D) \propto p(D \mid k_i, b) p(k_i, b), \tag{S208}$$

where the likelihood $p(D \mid k_i, b)$ is given by the product of $N$ negative binomials as in Eq. S207. We only need to choose priors on $k_i$ and $b$. For the datasets from [36] that we are analyzing, as for the Poisson promoter model above we are still data-rich so the prior's influence remains weak, but not nearly as weak because the dimensionality of our model has increased from one to two.

We follow the guidance of [52], Section 2.9 in opting for weakly-informative priors on $k_i$ and $b$ (conjugate priors do not exist for this problem), and we find "street-fighting estimates" [87] to be an ideal way of constructing such priors. The idea of weakly informative priors is to allow all remotely plausible values of model parameters while excluding the completely absurd or unphysical.

Consider $k_i$. Some of the strongest known bacterial promoters control rRNA genes and initiate transcripts no faster than $\sim 1$/sec. It would be exceedingly strange if any of the constitutive promoters from [36] were stronger than that, so we can take that as an upper bound. For a lower bound, if transcripts are produced too rarely, there would be nothing to see with FISH. The datasets for each strain contain of order $10^3$ cells, and if the $\langle m \rangle =$

$k_i b/\gamma \lesssim 10^{-2}$, then the total number of expected mRNA detections would be single-digits or less and we would have essentially no data on which to carry out inference. So assuming $b$ is not too different from 1, justified next, and an mRNA lifetime of $\gamma^{-1} \sim 3 - 5$ min, this gives us soft bounds on $k_i/\gamma$ of perhaps $10^{-2}$ and $3 \times 10^1$.

Next consider mean burst size $b$. This parametrization of the geometric distribution allows bursts of size zero (which could representing aborted transcripts and initiations), but it would be quite strange for the mean burst size $b$ to be below $\sim 10^{-1}$, for which nearly all bursts would be of size zero or one. For an upper bound, if transcripts are initiating at a rate somewhat slower than rRNA promoters, then it would probably take a time comparable to the lifetime of an mRNA to produce a burst larger than 10-20 transcripts, which would invalidate the approximation of the model that the duration of bursts are instantaneous compared to other timescales in the problem. So we will take soft bounds of $10^{-1}$ and $10^1$ for $b$.

Note that the natural scale for these "street-fighting estimates" was a log scale. This is commonly the case that our prior sense of reasonable and unreasonable parameters is set on a log scale. A natural way to enforce these soft bounds is therefore to use a lognormal prior distribution, with the soft bounds set $\pm 2$ standard deviations from the mean.

With this, we are ready to write our full generative model as

$$\begin{aligned} \ln k_i &\sim \text{Normal}(-0.5, 2), \\ \ln b &\sim \text{Normal}(0.5, 1), \\ m &\sim \text{NBinom}(k_i, b). \end{aligned} \tag{S209}$$

Section 4 in the main text details the results of applying this inference to the single-cell mRNA counts data. There we show the posterior distribution for the two parameters for different promoters. Figure S5 shows the so-called posterior predictive checks (see main text for explanation) for all 18 unregulated promoters shown in the main text.

#### S3.3 Bayesian inference on the simple-repression architecture

As detailed in 4 in the main text the inference on the unregulated promoter served as a stepping stone towards our ultimate goal of inferring repressor rates from the steady-state mRNA distributions of simple-repression architectures. For this we expand the one-state bursty promoter model to a two-state promoter as schematized in Figure 1(C) as model 5. This model adds two new parameters: the repressor binding rate $k^+$, solely function of the repressor concentration, and the repressor dissociation rate $k^-$, solely a function of the repressor-DNA binding affinity.

The structure of the data in [36] for regulated promoters tuned these two parameters independently. In their work the production of the LacI repressor was under the control of an inducible promoter regulated by the TetR repressor as schematized in Figre S6. When TetR binds to the small molecule anhydrotetracycline (aTc), it shifts to an inactive conformation unable to bind to the DNA. This translates into an increase in gene expression level. In other words, the higher the concentration of aTc added to the media, the less TetR repressors that

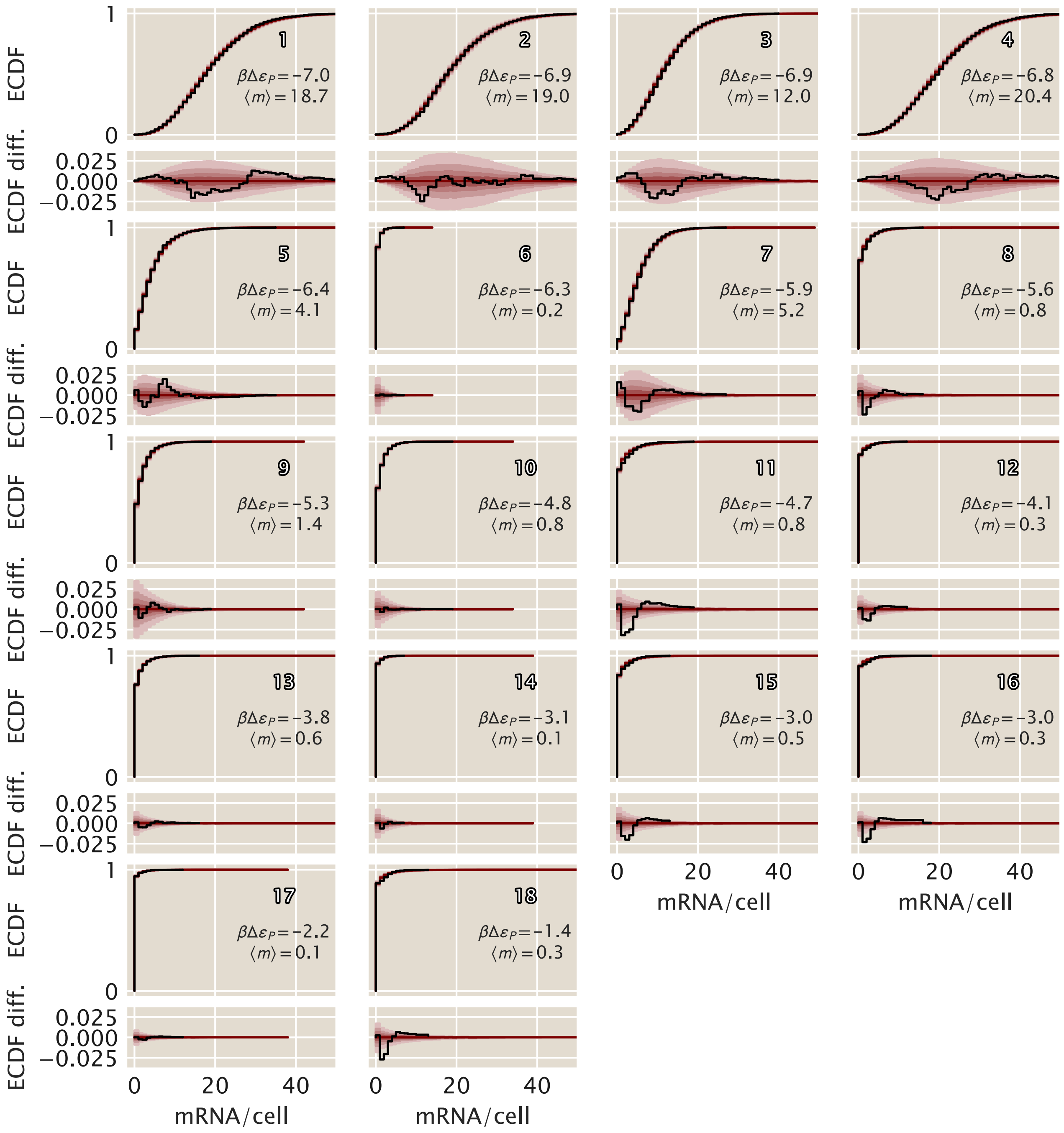

**Figure S5. Theory-data comparison of inference on unregulated promoters.** Comparison of the inference (red shaded area) vs the experimental measurements (black lines) for 18 different unregulated promoters with different mean mRNA expression levels from Ref. [36]. Upper panels show the empirical cumulative distribution function (ECDF), while the lower panels show the differences with respect to the median of the posterior samples. White numbers are the same as in Figure 1 for cross comparison. The predicted binding energies $\beta\Delta\varepsilon_p$ were obtained from the energy matrix model in Ref. [54]

can control the expression of the *lacI* gene, so the higher the concentration of LacI repressors in the cell. So by tuning the amount of aTc in the media where the experimental strains were grown they effectively tune $k^+$ in our simple theoretical model. On the other hand to tune $k^-$ the authors swap three different binding sites for the LacI repressor, each with different repressor-DNA binding affinities previously characterized [33].

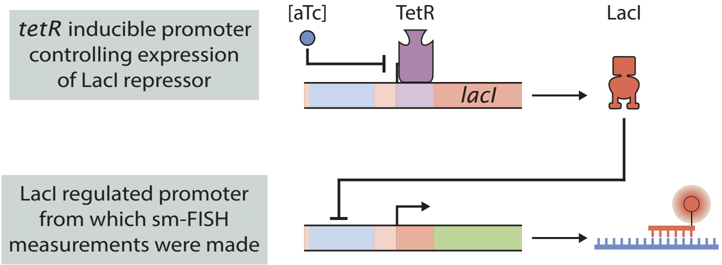

**Figure S6. aTc controlled expression of LacI repressor.** Schematic of the circuit used in [36] to control the expression of the LacI repressor. The *lacI* gene is under the control of the TetR repressor. As the TetR repressor is inactivated upon binding of anhydrotetracycline or aTc, the more aTc added to the media were cells are growing, the less TetR repressors available to control the expression of the *lacI* gene, resulting in more LacI repressors per cell. LacI simultaneously controls the expression of the mRNA on which single-molecule mRNA FISH was performed for gene expression quantification.

What this means is that we have access to data with different combinations of $k^-$ and $k^+$. We could naively try to fit the kinetic parameters individually for each of the datasets, but there is no reason to believe that the binding site identity for the LacI repressor somehow affects its expression level controlled from a completely different location in the genome, nor vice versa. In other words, what makes the most sense it to fit all datasets together to obtain a single value for each of the association and dissociation rates. What this means, as described in Section 4 of the main text is that we have a seven dimensional parameter space with four possible association rates $k^+$ given the four available aTc concentrations, and three possible dissociation rates $k^-$ given the three different binding sites available in the dataset.

Formally now, denote the set of seven repressor rates to be inferred as

$$\vec{k} = \{k_{Oid}^-, k_{O1}^-, k_{O2}^-, k_{0.5}^+, k_1^+, k_2^+, k_{10}^+\}. \tag{S210}$$

Note that since the repressor copy numbers are not known directly as explained before, we label their association rates by the concentration of aTc. Bayes theorem reads simply

$$p(\vec{k}, k_i, b \mid D) \propto p(D \mid \vec{k}, k_i, b) p(\vec{k}, k_i, b), \tag{S211}$$

where $D$ is the set of all $N$ observed single-cell mRNA counts across the various conditions. We assume that individual single-cell measurements are independent so that the likelihood factorizes as

$$p(D \mid \vec{k}, k_i, b) = \prod_{j=1}^{N} p(m \mid \vec{k}, k_i, b) = \prod_{j=1}^{N} p(m \mid k_j^+, k_j^-, k_i, b) \tag{S212}$$

where $k_j^{\pm}$ represent the appropriate binding and unbinding rates for the $j$-th measured cell. Our likelihood function, previously derived in Appendix S2, is given by the rather complicated result in Eq. S184, which for completeness we reproduce here as

$$\begin{aligned} p(m \mid k_R^+, k_R^-, k_i, b) = &\frac{\Gamma(\alpha+m)\Gamma(\beta+m)\Gamma(k_R^+ + k_R^-)}{\Gamma(\alpha)\Gamma(\beta)\Gamma(k_R^+ + k_R^- + m)} \frac{b^m}{m!} \\ &\times {}_2F_1(\alpha+m, \beta+m, k_R^+ + k_R^- + m; -b). \end{aligned} \tag{S213}$$

where $\alpha$ and $\beta$, defined for notational convenience, are

$$\begin{aligned} \alpha &= \frac{1}{2}\left(k_i + k_R^- + k_R^+ + \sqrt{(k_i + k_R^- + k_R^+)^2 - 4k_i k_R^-}\right) \\ \beta &= \frac{1}{2}\left(k_i + k_R^- + k_R^+ - \sqrt{(k_i + k_R^- + k_R^+)^2 - 4k_i k_R^-}\right). \end{aligned} \tag{S214}$$

Next we specify priors. As for the constitutive model, weakly informative lognormal priors are a natural choice for all our rates. We found that if the priors were too weak, our MCMC sampler would often become stuck in regions of parameter space with very low probability density, unable to move. We struck a balance in choosing our prior widths between helping the sampler run while simultaneously verifying that the marginal posteriors for each parameter were not artificially constrained or distorted by the presence of the prior. The only exception to this is the highly informative priors we placed on $k_i$ and $b$, since we have strong knowledge of them from our inference of constitutive promoters above.

With priors and likelihood specified we may write down our complete generative model as

$$\begin{aligned} \log_{10} k_i &\sim \text{Normal}(0.725, 0.025) \\ \log_{10} b &\sim \text{Normal}(0.55, 0.025) \\ \log_{10} k_{0.5}^+ &\sim \text{Normal}(-0.45, 0.3) \\ \log_{10} k_1^+ &\sim \text{Normal}(0.6, 0.3) \\ \log_{10} k_2^+ &\sim \text{Normal}(1.15, 0.3) \\ \log_{10} k_{10}^+ &\sim \text{Normal}(1.5, 0.3) \\ \log_{10} k_{Oid}^- &\sim \text{Normal}(-0.25, 0.3) \\ \log_{10} k_{O1}^- &\sim \text{Normal}(0.1, 0.3) \\ \log_{10} k_{O2}^- &\sim \text{Normal}(0.45, 0.3) \\ m &\sim \text{Likelihood}(k_R^+, k_R^-, k_i, b), \end{aligned} \tag{S215}$$

where the likelihood is specified by Eq. S213. We ran MCMC sampling on the full nine dimensional posterior specified by this generative model.

We found that fitting a single operator/aTc concentration at a time with a single binding and unbinding rate did not yield a stable inference for most of the possible operator/aTc combinations. In other words, a single dataset could not independently resolve the binding and unbinding rates, only their ratio as set by the mean fold-change in Figure 1 in the main text. Only by making the assumption of a single unique binding rate for each repressor copy number and a single unique unbinding rate for each binding site, as done in Figure 4(A), was it possible to independently resolve the rates and not merely their ratios.

We also note that we found it necessary to exclude the very weakly and very strongly repressed datasets from Jones et. al. [36]. In both cases there was, in a sense, not enough information in the distributions for our inference algorithm to extract, and their inclusion simply caused problems for the MCMC sampler without yielding any new insight. For the strongly repressed data (Oid, 10 ng/mL aTc), with $> 95\%$ of cells with zero mRNA, there was quite literally very little data from which to infer rates. And the weakly repressed data, all with the repressor binding site O3, had an unbinding rate so fast that the sampler essentially sampled from the prior; the likelihood had negligible influence, meaning the data was not informing the sampler in any meaningful way, so no inference was possible.
